# TISON: a next-generation multi-scale modeling theatre for *in silico* systems oncology

**DOI:** 10.1101/2021.05.04.442539

**Authors:** Mahnoor N Gondal, Muhammad U Sultan, Ammar Arif, Abdul Rehman, Hira A Awan, Zainab Arshad, Waleed Ahmed, Muhammad FA Chaudhary, Salaar Khan, Zain B Tanveer, Rida Nasir Butt, Risham Hussain, Huma Khawar, Bibi Amina, Rida Akbar, Fatima Abbas, Misha N Jami, Zainab Nasir, Osama S Shah, Hadia Hameed, Muhammad FA Butt, Ghulam Mustafa, Muhammad M Ahmad, Sameer Ahmed, Romena Qazi, Fayyaz Ahmed, Omer Ishaq, Syed W Nabi, Wim Vanderbauwhede, Bilal Wajid, Huma Shehwana, Emad Uddin, Muhammad Safdar, Irfan Javed, Muhammad Tariq, Amir Faisal, Safee U Chaudhary

## Abstract

Multi-scale models integrating biomolecular data from genetic, transcriptional, and translational levels, coupled with extracellular microenvironments can assist in decoding the complex mechanisms underlying system-level diseases such as cancer. To investigate the emergent properties and clinical translation of such cancer models, we present Theatre for *in silico* Systems Oncology (TISON, https://tison.lums.edu.pk), a next-generation web-based multi-scale modeling and simulation platform for *in silico* systems oncology. TISON provides a “zero-code” environment for multi-scale model development by seamlessly coupling scale-specific information from biomolecular networks, microenvironments, cell decision circuits, *in silico* cell lines, and organoid geometries. To compute the temporal evolution of multi-scale models, a simulation engine and data analysis features are also provided. Furthermore, TISON integrates patient-specific gene expression data to evaluate patient-centric models towards personalized therapeutics. Several literature-based case studies have been developed to exemplify and validate TISON’s modeling and analysis capabilities. TISON provides a cutting-edge multi-scale modeling pipeline for scale-specific as well as integrative systems oncology that can assist in drug target discovery, repositioning, and development of personalized therapeutics.

## Introduction

Biological systems are tightly regulated by a multifactorial interplay of biomolecular entities at both inter and intracellular scales together with extracellular environments (1,2). These entities including genes, transcripts, proteins, and metabolites, interact to function over a wide range of spatiotemporal scales (1,3) in well-orchestrated regulatory pathways (4,5). These pathways are further interlinked and form biomolecular interaction networks such as gene-regulatory, protein-protein interaction networks, and metabolic networks (6). These networks regulate the life cycle of cells by cell-type-dependent modulation of pathways resulting in programing of specific cell fates including cell death, proliferation, and differentiation (6,7). Moreover, living cells are assembled into tissues, which along with their extracellular environmental milieu create functional organs (8,9). This heterogeneous yet hierarchical nature of regulation in biological systems, accompanied by the spatial and temporal diversity at each scale, obscures the understanding of system-level manifestations of complex diseases such as cancer (10), Alzheimer’s (11), and diabetes (12).

Specifically, in the case of cancer, the disease begins with mutations at genetic and epigenetic levels (13,14), thereby dysregulating biomolecular pathways which then escalates up to tissues and organs. The diversity of these regulatory aberrations gives rise to vast genotypic and phenotypic heterogeneity (15), which is a major impediment in understanding and treatment of disease (16). This necessitates determining the role played by each biomolecule in bringing about holistic system-level outcomes (17). A key challenge in modern cancer biology is, therefore, to perform integrative investigations of complex inter-, intra-, and extracellular biomolecular regulations that give rise to system-level effects (1,2,18).

Rapid advancements in molecular biology, particularly in high-throughput genomics, transcriptomics, proteomics, and metabolomics have generated a vast amount of complex spatiotemporal data in both physiological and pathological contexts (19,20). Such experimental data now populates several online expression databases e.g. Catalogue of Somatic Mutations in Cancer (COSMIC) (21), Genotype-Tissue Expression (GTEx) (22), The Cancer Genome Atlas (TCGA) (23), Metabolic gEne Rapid Visualizer (MERAV) (24), The Cancer Proteome Atlas (TCPA) (25), Human Protein Atlas (HPA) (26), Human Metabolome Database (HMDB) (27), etc. Integrative computational models employing the multi-scale expression data from these repositories can help decode emergent system-level properties associated with cancer. The domain of *in silico* systems oncology (28), encompasses the utilization of such omics-based data from different spatiotemporal scales to study cancer growth, progression, and treatment using computational methods (29).

Until recently, numerous mathematical and computational models employing the aforementioned biological data have been reported for investigating multi-scale cancer systems biology (30–37). These include models of morphological development of solid tumors in normoxia and hypoxia (38), decoding the dynamics of homeostasis from cell-based multi-scale models of colon crypts (39), and model of glucose metabolism and its role in cancer growth and progression (40), alongside others (30–37). Subsequently, to facilitate the model development process, several multi-scale cancer modeling platforms have been developed (41–43). Amongst these, CompuCell (2), reported in 2004, allowed modeling of multi-cellular organisms by integrating gene regulatory networks, cell and extracellular matrix (ECM), and cell-cell and cell-microenvironment interactions. However, its use of the cellular potts model increased the computational cost for large-scale models while the calibration of the Monte Carlo time step to physical time step makes its employment in multi-scale modeling challenging (44). In comparison, ELECANS (Electronic Cancer System) (45) provided a feature-rich modeling platform for building multi-scale models to decode the multifactorial underpinnings of tumorigenesis. The platform’s Software Development Kit (SDK), however, also placed a heavy C# programming requirement for its users. Similarly, CHASTE (Cancer Heart and Soft Tissue Environment) (46) required test-driven development by using several mathematical modeling frameworks for solving Ordinary and Partial Differential Equations (ODEs/PDEs). Like ELECANS, the programming skills required for using CHASTE also hindered its utilization by conventional wet-lab biologists and clinicians. R/Repast (47), a recently reported platform, provided a High-Performance Computing (HPC) capability towards greater scalability but lacked a programming interface for implementing subcellular biomolecular models. Nevertheless, the lack of a generic and intuitive software providing a “zero-code” modeling environment continues to impede the development and employment of complex multi-scale biological models in research laboratories and clinical settings (48).

In this work, we propose a next-generation web-based multi-scale modeling platform “TISON” -Theatre for *in silico* Systems Oncology. TISON provides a “zero-code” modular environment, which is conveniently employable by modelers, experimental biologists, and clinicians, alike. The software comprises of eight scale-specific editors: (i) the Networks Editor (NE) allows for construction and analysis of rules and weight-based biomolecular networks, (ii) Therapeutics Editor (TE) helps develop therapeutic screens on biomolecular networks developed using NE towards identification of novel drug targets, drug repurposing, and personalized therapeutics, (iii) Environments Editor (EE) assists in the creation of diffusive microenvironments towards modeling the dynamical engagement of environmental cues with cellular organoids, (iv) Cell Circuits Editor (CCE) helps construct cell decision circuits as Finite State Machines (FSMs) (49) for computing the cell fate outcomes in light of biomolecular network regulation and microenvironment, (v) Cell Lines Editor (CLE) then assigns these cell circuits to *in silico* cell line models, (vi) Organoids Editor (OE) employs *in silico* cell lines to create tissue organoid systems, (vii) Simulations Editor (SE) simulates the tissue organoids to investigate their spatiotemporal evolution, and (viii) Analytics Editor (AE) queries simulation data and visualizes the analysis results.

To exemplify TISON’s features, we have replicated several case studies from published literature and compared their results thus validating each editor’s functionality (38,50–53). Taken together, TISON provides a next-generation multi-scale modeling platform for *in silico* systems oncology and can assist in unraveling the multifactorial interplay underpinning tumorigenesis. The software can provide significant impetus to the clinical applications of cancer systems biology by creating avenues for translation of molecular research towards targeted and personalized therapeutics.

## Results

Theatre for *in silico* Systems Oncology (TISON) platform is designed using the distributed three-tier software architecture consisting of components organized into front-end, middleware, and back-end. The front-end consists of eight web-based graphical user interfaces (GUIs) termed “*editors”*. Each editor provides scale-specific modeling features and setting up of associated parameters towards a scale-by-scale development of systems oncology models. The resulting models are taken up by the middleware that consists of a high-performance simulation engine, which has been implemented as a Microsoft® .Net Web Application Programming Interface (WebAPI). The back-end employs a Microsoft® SQL server for model data storage and retrieval. The resulting three-tier distributed software architecture is depicted in Figure 1 (see also Supplementary Material, Section 1).

**Figure 1.**
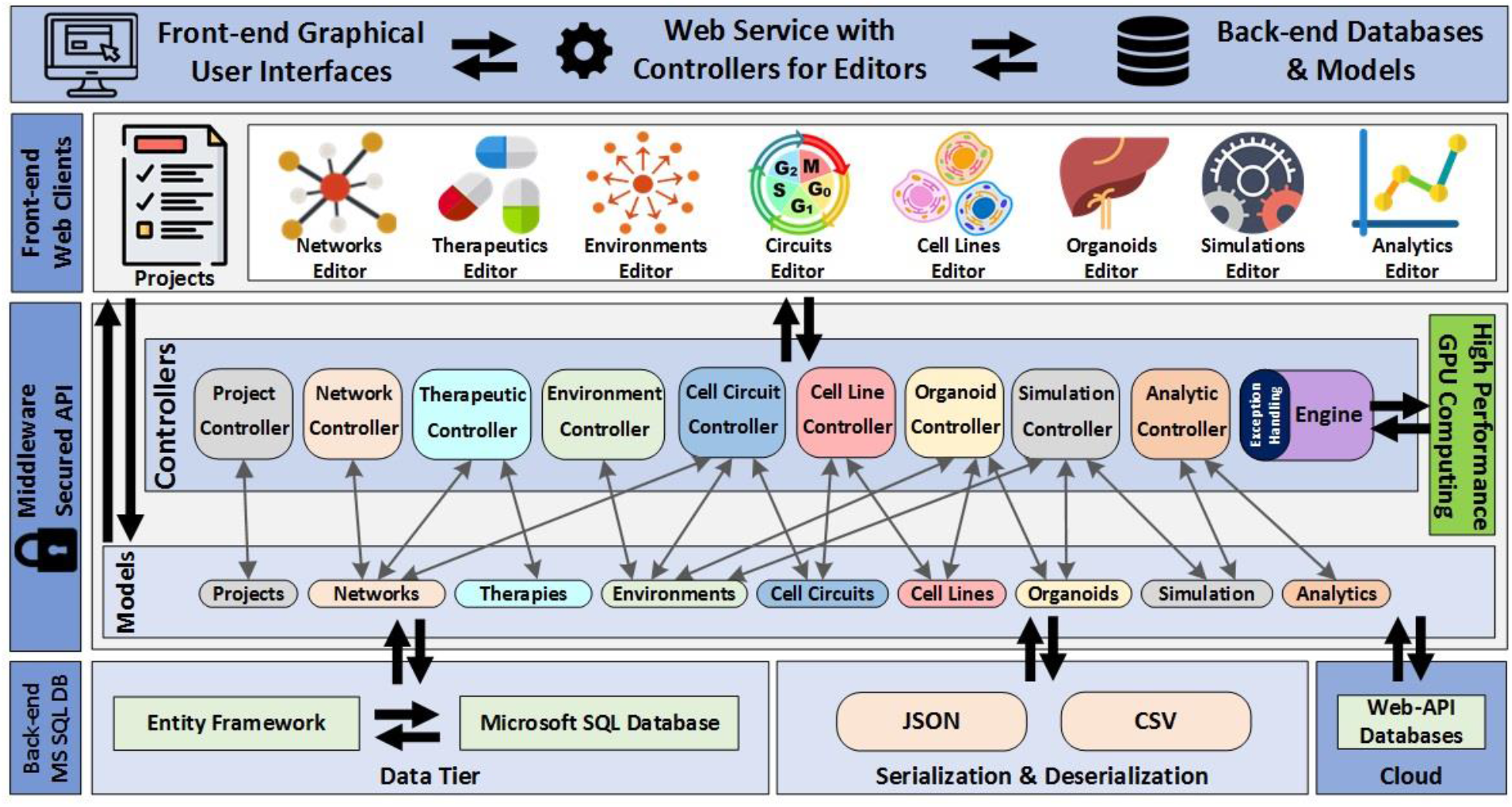
A Schematic of TISON Software Architecture. TISON is constructed as a three-layer application comprising of the front-end, middleware, and back-end. The front-end includes a web application that contains eight editors with corresponding GUI’s. The middleware consists of controllers and models, which take user-defined parameters from the GUIs, compute the simulation logic in light of model parameters, and store the results in the database at the back-end. The back-end stores serialized data in a Microsoft SQL database for provision to the middleware for onward processing.

TISON users can begin the model construction process by creating a project followed by defining the world size (see Supplementary Material, Section 1) following which scale-specific modeling components can be developed in TISON editors. In the following sub-sections, we elaborate on the salient features of TISON editors and exemplify their employment through literature-based case studies (see Supplementary Material, Section 2, Supplementary Figure S1).

### Networks Editor – Design and analysis of biomolecular regulatory networks

The process of multi-scale modeling in TISON begins with the construction of biomolecular networks using *Networks Editor* (NE). Users can choose between creating (i) rules-based, or (ii) weight-based network models for onward analyses. For rules-based networks, NE allows defining of Boolean rules as abstractions of biomolecular regulation. Deterministic analysis (DA) (54) can then be performed on these networks towards investigating their regulatory dynamics and cell fate outcomes. Results obtained from DA can be visualized as cell fate and attractor landscapes (51,55). NE also allows the conversion (56) of rules-based networks into weight-based networks. In weight-based networks, users can manually assign expression values to each node or import them from online databases that are readily available in NE. The databases include Metabolic gEne Rapid Visualizer (MERAV) (24), Human Proteome Atlas (HPA) (57), The Cancer Genome Atlas (TCGA) (58) through Firebrowse (59), and The Genotype-Tissue Expression project (GTEx) (22). The expression values can then be employed to calculate the basal level expression for each node in a weight-based network using an in-built feature in NE. This is especially useful in developing personalized cancer network models. For weight-based network analysis, NE allows its users to perform (i) DA, (ii) Probabilistic Analysis (PA) (60), or (iii) Ordinary Differential Equation (ODE) analysis (61). Results obtained can be downloaded and visualized as an attractor and cell fate landscape (51,55) for DA; probability, potential energy, and cell fate landscapes (52) for PA; and ODE landscapes (61) for ODE analysis. Further, NE users can also undertake multiple in tandem analyses towards evaluating robustness and parameter sensitivity of networks (62) along with results visualization using a variety of graphs and charts (see Supplementary Material, Section 2.1).

To exemplify and validate the functionality of NE, we have reconstructed four different case studies from published literature (Figure 2, see Supplementary Material, Section 2.1.2). In the first case study, we created a 197 node human colorectal tumorigenesis signaling network based on the work of Cho *et al*. (50), to validate NE’s rules-based DA pipeline (Figure 2A). Results from DA showed a normal response by the human signaling network in the absence of mutations and tallied with the published results (Figure 2B; Supplementary Figure S2). In the second case study, we constructed a 16 node p53 network of MCF-7 breast cancer cell lines (51) and evaluated NE’s weight-based DA pipeline (Figure 2C). TISON accurately reproduced the state transition dynamics of each cell fate outcome, in the absence and presence of DNA damage, in line with the published study (Figure 2D; Supplementary Figures S3-6). Towards assessing TISON’s PA functionality, a yeast cell cycle progression network consisting of 11 nodes (52) was reconstructed in NE (Figure 2E). Analysis of the network reproduced the global minimum, G1 state, as hypothesized in the original study (Figure 2F; Supplementary Figures S7-8). Lastly, to compare and validate NE’s ODE analysis pipeline, a 52 node human stem cell developmental network (53) was adopted for comparison with a published tool “NetLand” (61) (Figure 2G). Results from TISON’s ODE network analysis reported two embryonic stem cell marker genes, *Nanog* (stem cell marker gene) and *Gata6* (differentiation marker gene), along with the corresponding stem cell and differentiated state attractors as reported by NetLand (Figure 2H; Supplementary Figures S9-11).

**Figure 2.**
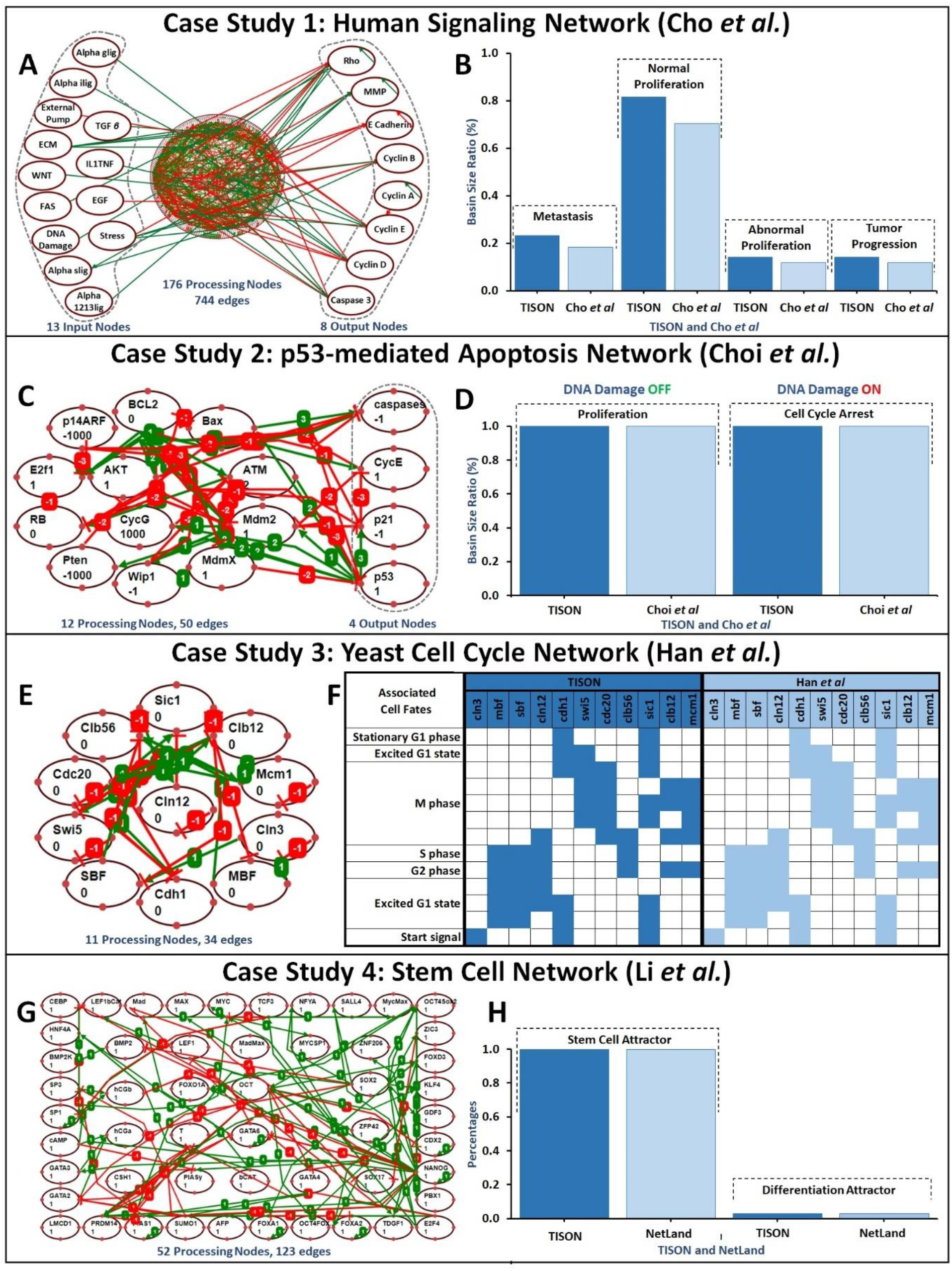
Case Studies for TISON’s Networks Editor. (**A**) Cho *et al*.*’s* rules-based Human Signalling Network constructed using TISON’s NE. The network contains 13 input nodes, 176 processing nodes, 8 output nodes, and 744 edges. (**B**) Comparison between Cho *et al*. and TISON’s deterministic analysis results. (**C**) Choi *et al*.*’s* weight-based p53 network containing 12 processing nodes, 4 output nodes, and 50 edges (**D**) Comparison between Choi *et al*. and TISON’s deterministic analysis results in the absence and presence of DNA damage. (**E**) Han *et al*.*’s* weight-based Yeast Cell Cycle Network containing 11 nodes and 34 edges. (**F**) Comparison between results reported by Han *et al*. and TISON’s probabilistic analysis pipeline. (**G**) Wang *et al*.*’s* weight-based Stem Cell Network containing 52 nodes and 123 edges. (**H**) Comparison between NetLand and TISON’s ODE analysis of Wang *et al*.’s network.

The aforementioned literature-based case studies validated the salient biomolecular network analysis modalities and associated landscape visualization functionalities of TISON’s NE (see Materials and Methods) (see Supplementary Material, Section 2.1.2).

### Therapeutics Editor – Developing therapeutic screens towards identification of novel drug targets, drug repurposing, and personalized therapeutics

TISON’s *Therapeutics Editor* (TE) assists in undertaking a therapeutic evaluation of biomolecular networks developed using the NE. Users can create “*therapies”* by employing information on drugs and their targets. Each therapy consists of a single or a combination of drugs and each drug may target one or more network nodes or edges. Drug action can involve: (i) the enhancement or suppression of node activity (i.e. gain of function, “*knock-up*” or loss of function “*knockdown*”), (ii) node removal (“*knock-out*”), (iii) node addition (“*knock-in*”), or (iv) regulation of target node expression by modifying edge interactions. For rules-based networks, node knock-up or knockdown is implemented by assigning a fixed user-defined value to the node thereby suppressing its upstream regulation. Wherein, in the case of rules-based node knock-out, the node is deleted from the network, and its rule is removed; whereas for the knock-in case, the new node along with its associated rule is added into the network by updating network rewiring. Similarly, in the case of weight-based networks, knock-up or knockdown is implemented by fixing the node expression value along with suppression of its upstream regulation and keeping its basal value at ‘0’; knock-out is performed by updating the node’s basal value and fixed node value to ‘0’, and deleting all of its outgoing as well as incoming interactions; weight-based node knock-in is implemented by defining a new node, assigning its upstream and downstream regulation and adding its basal value. Basal value in TE can be added using two ways, users can either directly assign a node’s basal value at the time of node creation or, it can be calculated using expression data provided by the user towards developing a personalized cancer network model. Lastly, users can also alter, knock-in, or knock-out the edge weights between source and target nodes in weight-based networks by updating or adding their interaction value. For such therapy-targeted nodes, specific scores can also be imported from the Drug-gene Interaction Database (DGIdb) (63), for ease in implementation.

Towards creating targeted therapy, TE users can proceed in two modalities i.e. *horizontal* or *vertical* therapy (see Supplementary Material, Section 2.2). A *horizontal therapy* is a single-drug therapy, wherein the drug may target one or more nodes or node interactions concurrently. Whereas, *vertical therapy* comprises multiple in tandem horizontal therapies. Upon creation of a therapy, users can analyze the network (detailed in NE, above) in the presence of the therapy towards computing cell fate outcomes. Results obtained from these analyses can be visualized using a variety of landscapes (see Supplementary Material, Section 2.2.1). Moreover, users can employ the inbuilt cell fate propensities comparison feature to compare the effect of drugs against the control case. Additionally, TE provides an *Exhaustive Screening* step, in which, users can exhaustively evaluate each or selected node in the network towards evaluating the most efficacious target nodes in the model system in light of patient-specific mutations. This feature is especially useful in predicting efficacious drug targets in light of patient’s mutation data towards developing personalized cancer therapeutic combinations. Gondal *et al*. (82) employed this feature to identify a synergistic combination of paclitaxel and pazopanib for treating colorectal cancer.

To validate and demonstrate TE’s functionality, we recreated two published case studies. In the first case study, we replicated the cellular response to DNA damage using a weight-based MCF-7 p53-mediated apoptosis network consisting of 16 nodes and 50 edges (51). Targeted therapies comprising of Wip1 and Nutlin were then introduced into the network. In both the absence and presence of DNA damage, the highest efficacy was exhibited by a combinatorial therapy (Nutlin+Wip1 inhibition) leading to increased levels of apoptosis and senescence (Figure 3A; Supplementary Figures S12-23). The results were comparable with those of the published study (51). In the second case study, TE was used to design a therapeutic screen customized to introduce genetic mutations associated with Colorectal Cancer (CRC) into a normal network to transform it into a cancerous network. For that, we introduced progression CRC mutations (APC, RAS, PTEN, and TP53) into a rules-based human signaling network comprising 197 nodes and 744 edges (50). The results, using DA, exhibited the emergence of CRC with enhanced oncogenic cell fate propensities, including abnormal proliferation and metastasis, in line with Cho *et al*. (50) (Figure 3B; Supplementary Figures S24-31, see Supplementary Material, Section 2.2.2).

**Figure 3.**
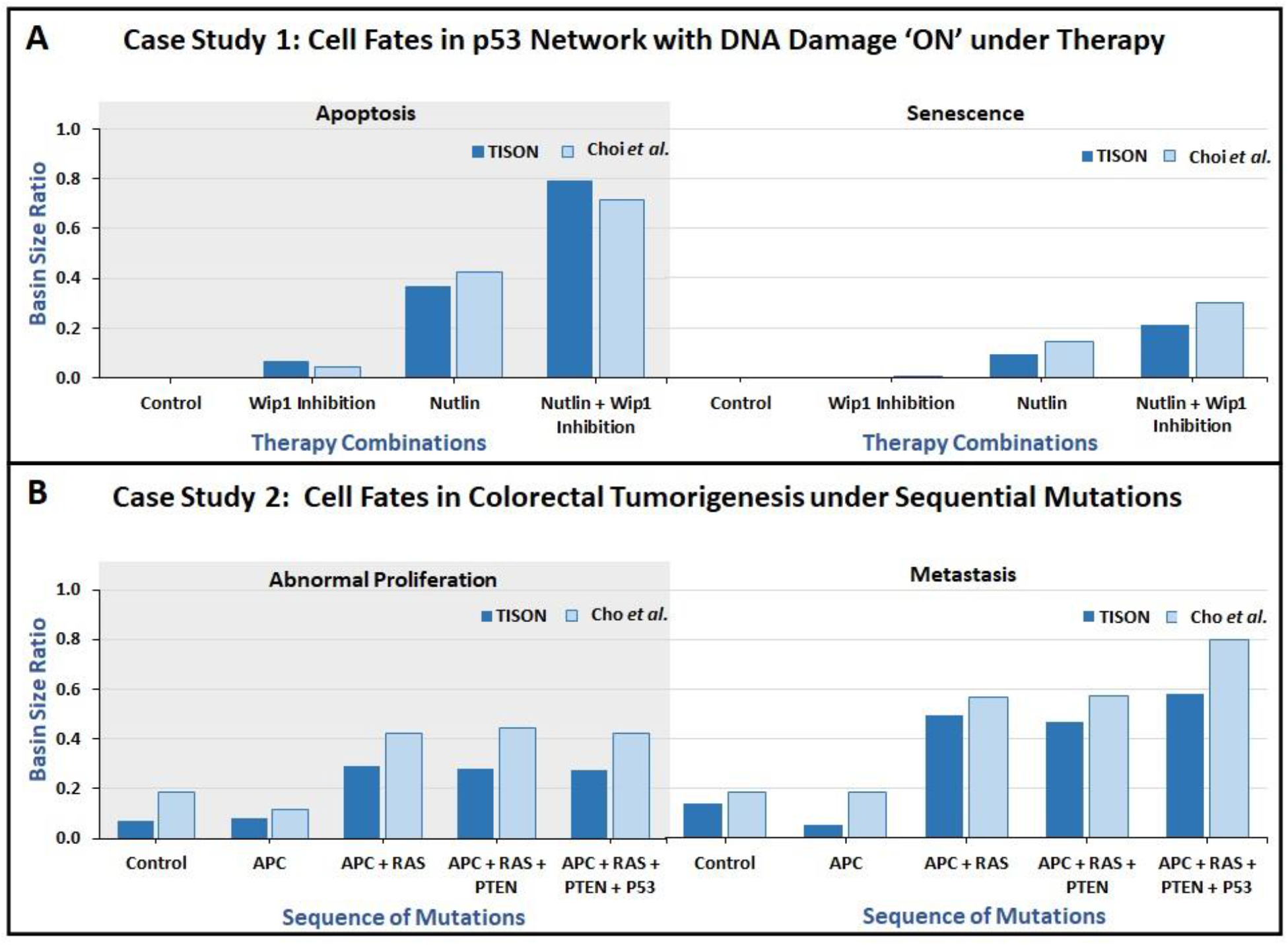
Case Studies for TISON’s Therapeutics Editor. (**A**) Comparison of drug screening results from Choi *et al*. and TISON’s TE with DNA damage ON. (**B**) Comparison of results from Cho *et al*. and TISON’s therapeutics editor for induction of CRC. TE was used to introduce successive mutations into the network (APC, RAS, PTEN, and P53) leading to the emergence of CRC.

### Environments Editor – Creating models of diffusive microenvironments models

TISON’s *Environments Editor* (EE) allows its users to design models of the cellular microenvironment (64) for setting up specific biological contexts such as tumor microenvironment (65) and hypoxia (66), etc. For that, EE models extracellular environments as a collection of “*layers*” wherein each layer is a continuous diffusive field of a specific type (64) such as a nutrient, morphogen, or cellular secretions. Each diffusive field is modeled using a partial differential equation (PDE) (67) and users can set up associated parameters such as diffusion constant, initial concentration, and boundary condition (68). To compute the spatiotemporal diffusion of biomolecules within each layer, an in-house implementation of the unconditionally stable Alternating Directions Implicit (ADI) method is provided for numerical estimation of the PDE models (69,70) (see Supplementary Material, Section 2.3 for details).

### Cell Circuits Editor – Constructing cell decision circuits as Finite State Machines

TISON’s *Cell Circuits Editor* (CCE) allows users to create cell state transition models in the form of finite state machines (FSM) (73) to help compute the overall cellular state (74– 76). CCE integrates NE constructed biomolecular network and EE designed extracellular entities such as drugs, nutrients, and signaling molecules, in the form of a “*Cell Circuit*”. Users can also incorporate intracellular processes such as cellular aging and growth into cell circuits by defining logical rules or mathematical equations for them. Each cell circuit maintains its internal state through a set of variables that contains associated networks’ node expressions, and the concentration of environmental biomolecules. Each variable can then be used in a “decision box” for choosing between different cell fates, under specific user-defined conditions. To further facilitate the users, several ready-to-use cell fates including mitosis, quiescence, cell death, migration, and differentiation have also been provided in CCE. Alongside this, an intuitive feature for the consumption and production of environmental biomolecules has been provided (see Supplementary Material, Section 2.4 for further details). Lastly, during the designing of cell circuits, users can simulate their circuits within a single time step to debug logical errors within their circuits.

To validate the functionality of CCE, we have reconstructed a literature-based case study by Gerlee and Anderson, to investigate the impact of oxygen concentration on cell population growth (38). The case study was undertaken in three stages (Figure 4); the first stage (Figure 4A) involved the development of a cell circuit to model cell death, mitosis, and quiescence, in the light of cell age and oxygen consumption (see Supplementary Figures S32, see Supplementary Material, Section 2.4.2 for further details). In the second stage (Figure 4B), an extracellular environment containing oxygen was incorporated into this circuit in the form of a diffusive environmental layer (see Supplementary Figures S33). Thirdly (Figure 4C), two variants of a biomolecular network regulating the p53-mediated apoptosis in MCF-7 breast cancer cell line (details in NE) having DNA damage ON and OFF (51) were incorporated into cell circuits. (see Supplementary Figures S34-35). The results obtained by simulating the complete cell circuit (from the third stage) exhibited proliferation to be the salient cell fate outcome in presence of DNA damage (i.e. without mutation), which is in agreement with Choi *et al*.*’s* network (51) (see Supplementary Material, Section 2.4.2 for further details).

**Figure 4.**
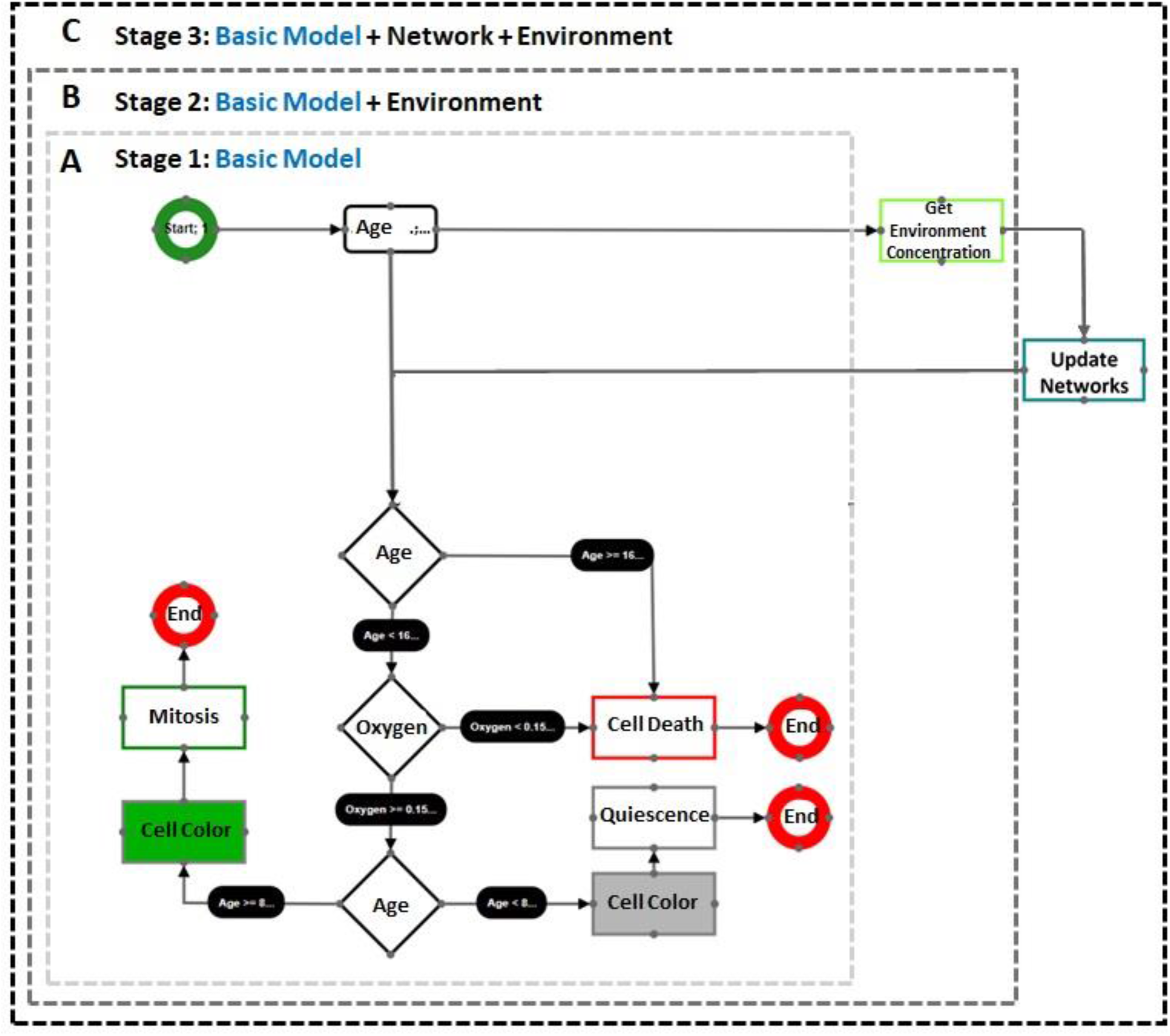
Case Study for TISON’s Cell Circuits Editor. CCE was used to design Gerlee and Anderson’s “*minimal*” model in three stages. **Stage 1:** Reconstruction of the “*minimal*” model. **Stage 2:** Incorporation of the extracellular environment into the model. **Stage 3:** Integration of biomolecular networks (DNA damage ON and OFF) into the model

### Cell Lines and Organoids Editor – Creating *in silico* cell lines towards developing tissue organoid systems

TISON’s *Cell Lines Editor* (CLE) assists in the creation of *in silico* counterparts of *in vitro* cell lines such as HCT-116 colon cancer cell lines (77) and MCF-7 breast cancer cell lines (78). The process involves the assignment of a cell circuit (designed using CCE) to a cell line. This coupling implicitly associates cell lines with an extracellular environment and biomolecular networks that are constructed earlier using EE and NE, respectively (see Supplementary Material, Section 2.5 for further details). *In silico* cells derived from these cell lines can then be assembled into three-dimensional tissues, termed “*organoids*” using TISON’s *Organoids Editor* (OE) (see Supplementary Material, Section 2.5 for further details).

Users can conveniently design and populate tissues by converting one or more monolayer cell lines into three-dimensional organoids and couple them with microenvironments (e.g. matrigel), to create a variety of tissue geometries representing different organs. The spatial coordinates of cells in an organoid can be defined in three ways: with a coordinate range, an equation, or with a custom coordinates file, allowing the users to create models of complex organoids models. Additionally, OE provides a resupply feature to simulate nutrient resupply and signaling molecule induction by coupling vessel and nerve elements with environmental layers from EE. OE further facilitates its users by providing predefined organoid geometries within the editor and the ability to individually modify any organoid element within an organoid through the “*Organoids Elements*” panel (see Supplementary Material, Section 2.6 for further details, see Supplementary Material, Section 2.5.2).

To exemplify and validate the functionality of CLE and OE, we have reconstructed two literature-based case studies investigating organoid growth in the presence and absence of DNA damage. In the first case study, we designed an MCF-7 breast epithelial *in silico* cell line that utilizes the DNA damage OFF network from Choi *et al*. (51) and cell circuit reported by Gerlee and Anderson (38) (see Supplementary Figures S36, see Supplementary Material, Section 2.6.2). The cell lines thus developed can be exported from the editor or conveniently employed in organoids editor for onward simulation. In the second case study, we employed an *in silico* MCF-7 breast epithelial cell line (see Supplementary Figures S45) with DNA damage ON followed by its integration into Gerlee and Anderson’s circuit (see Supplementary Figures S37). These *in silico* MCF-7 cell lines were used to create three-dimensional organoids representative of breast epithelial organoids in OE, ready for export as well as spatiotemporal simulations (see Supplementary Figures S38) (see Supplementary Material, Section 2.5.2 and 2.6.2 for further details).

### Simulations Editor – Investigating spatiotemporal evolution of resultant tissue organoid system

TISON’s *Simulations Editor* (SE) assists its users to simulate the spatiotemporal evolution of *in silico* organoids, developed using OE. Users can set the duration for which the organoid is to be simulated by providing the number of time steps. During organoid simulation, users can visually observe tissue evolution over space and time resulting from the complex multi-scalar interplay of the underlying biomolecular networks, extracellular environments, and cell decision circuits (detailed in the previous sections). Data generated during this process enables TISON users to decipher the multifactorial interplay at each scale within these models. SE, therefore, can provide scale-wise insights into the developmental evolution of tumor organoids and compare them with control cases (see Supplementary Material, Section 2.7 for further details).

To exemplify the features provided by SE, we have constructed four literature-based case studies on MCF-7 breast epithelium cell lines including (i) DNA damage ON, and (ii) DNA damage OFF. For that, two variants of Choi *et al*.*’s* p53 network with DNA damage ‘ON’ and DNA damage ‘OFF’ were created using NE. An oxygen microenvironment was designed using EE and coupled with the networks through a “*minimal”* cell circuit model reported by Gerlee and Anderson, using CCE. The resultant cell circuits (having networks with DNA damage ON and OFF, in the presence and absence of therapies) were used to create two *in silico* cell lines in CLE. Next, breast epithelium organoids were designed in OE using these *in silico* cell lines. Results from the simulation of these cell lines (see Supplementary Material, Section 2.7.2 for further details) showed that with DNA damage ON and OFF, the organoid failed to grow (see Supplementary Figures S39-40) as cells underwent cell cycle arrest.

### Analytics Editor – Quering simulation data and visualizing the analysis results

TISON’s *Analytics Editor* (AE) assists its users to query the organoid simulation data generated using the SE (detailed in the previous section) towards analysing the biomolecular, spatial, and temporal properties of an organoid. Users can query spatiotemporal data and plot (i) overall cell population, (ii) cell line-specific cell count, (iii) organoid center (iv) organoid mobility, (v) organoid specific variables, (vi) heatmaps of microenvironment consumption (vii) cell circuit-specific variables, as well as (viii) cell count by color for phenotypic visualization (see Supplementary Figures S41-44, Supplementary Material, Section 2.8 for further details).

## Discussion

Multi-scale modeling in systems biology involves developing integrative models of genes, transcripts, proteins, cells, and investigating their regulatory dynamics over diverse spatiotemporal scales. Simulations of such models can be used to decode the biomolecular foundations of emergent system-level properties as well as identifying novel therapeutic targets (79,80). Here, we propose “Theatre for *in silico* Systems Oncology” (TISON), which is a web-based next-generation platform for constructing multi-scale cancer systems biology models. The software embodies eight scale-specific editors featuring enriched graphical user interfaces (GUIs) for intuitive model development.

The multi-scale model building process begins with TISON’s *Networks Editor* (NE) wherein users can construct and analyze biomolecular regulatory networks. Users can also plugin expression data from online databases including Metabolic gEne Rapid Visualizer (MERAV) (24), Human Proteome Atlas (HPA) (57), The Cancer Genome Atlas (TCGA) (58) through Firebrowse (59), and The Genotype-Tissue Expression project (GTEx) (22) into networks. NE can perform deterministic, probabilistic, and ordinary differential equation model analyses to examine the cell fates programmed by biomolecular regulatory networks under specific conditions. Users can also analyze networks under input perturbations and noise towards mimicking stochasticity that exists in biological systems for evaluating network robustness (81). The pipeline can also be used to undertake parameter sensitivity analyses on the networks. Further, TISON’s *Therapeutics Editor* (TE) can help to develop therapeutic screens for the identification of novel drug targets, drug repurposing, and development of personalized therapeutics using the biomolecular networks constructed in NE. Users can design the extracellular matrix (ECM) (64) of a cell by utilizing TISON’s *Environments Editor* (EE), to create environmental models of normal or tumor microenvironments (65) for onward integration into cellular models. To couple the biomolecular networks with extra-cellular environmental milieu containing drugs, nutrients, and cellular secretions (ligands or ROS), etc, TISON allows the creation of cell decision circuits in TISON’s *Cell Circuits Editor* (CCE). Onwards, these cell circuits can be employed to design *in silico* cell lines using TISON’s *Cell Lines Editor* (CLE). Users can then employ the designed cell lines to develop three-dimensional tissue organoids using TISON’s *Organoids Editor* (OE). TISON’s *Simulations Editor* (SE) can simulate these organoids for evaluating their growth and evolution over time under user-specified environmental conditions and biomolecular regulation. Lastly, simulation data generated in SE can be queried in the *Analytics Editor* (AE) for cell population data analysis along with other advanced queries. TISON allows models to be simulated over time using a multi-agent simulation core which then serializes simulation data, produced during the process, to the backend databases. Note that, each editor implements data exchange in open data formats and is accompanied by detailed user manuals. Moreover, literature-based case studies are provided in each editor to exemplify and validate its functionality, besides helping users reconstruct the exemplars.

Specifically, in terms of platform adaptability, TISON provides a zero-code GUI which can be conveniently employed by systems biologists, experimental biologists, researchers and clinicians, alike. In comparison, CHASTE (46), a leading multi-scale modeling platform does not provide a GUI and can only be executed by command line. Furthermore, it requires recompilation on part of the modeler. Likewise, ELECANS (45) offers a rigid GUI, necessitating the modeler to write code in C++ or C# to program its software development kit (SDK) interface. CompuCell (2) provides an elaborative GUI, but the platform’s lattice and time step calibration limited its employment in multi-scale modeling. The focus of these modeling tools has been set on multi-agent simulations rather than multi-scale cancer modeling. The high-performance version of Repast (47) does not have any GUI or SDK interface for implementing subcellular mechanisms e.g. gene, protein, and metabolic networks.

Unlike these modeling platforms, TISON facilitates its users by providing ease of model customization and reproducibility, high-performance simulation core, and integration of expression databases. Normalized database schemas have been employed in TISON coupled with the model-view architecture for creating the web-based GUIs. In contrast with legacy modeling platforms, TISON’s salient software components are developed around the concept of Software as a Service (SaaS), while data import and export is completely performed in open data formats. Moreover, the low-cost and off-the-shelf GPU cluster enables fast model simulations. Another novel aspect of TISON is its feature to intuitively develop finite state machine models of cell decision circuits using drag-and-drop. To the best of our knowledge, no other contemporary cancer modeling tool provides such a feature. Lastly, TISON offers an enhanced simulation runtime performance in comparison to existing platforms. A feature-by-feature comparison of TISON with its contemporaries is tabulated in Figure 5, outlining the specific novelties offered by the platform.

**Figure 5.**
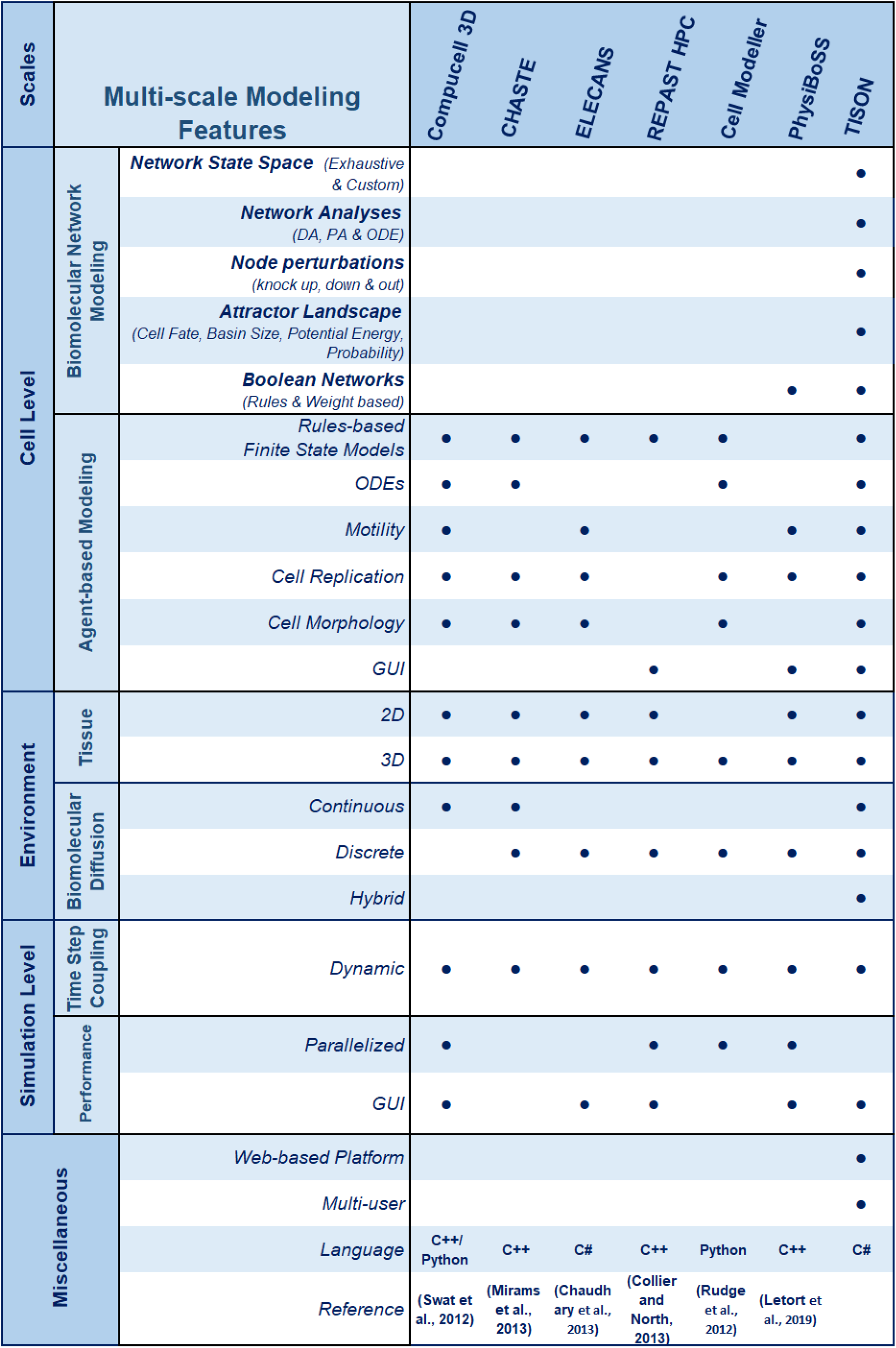
TISON Comparison with other Tools. Feature-by-feature comparison of TISON with other modeling tools.

**Figure 6.**
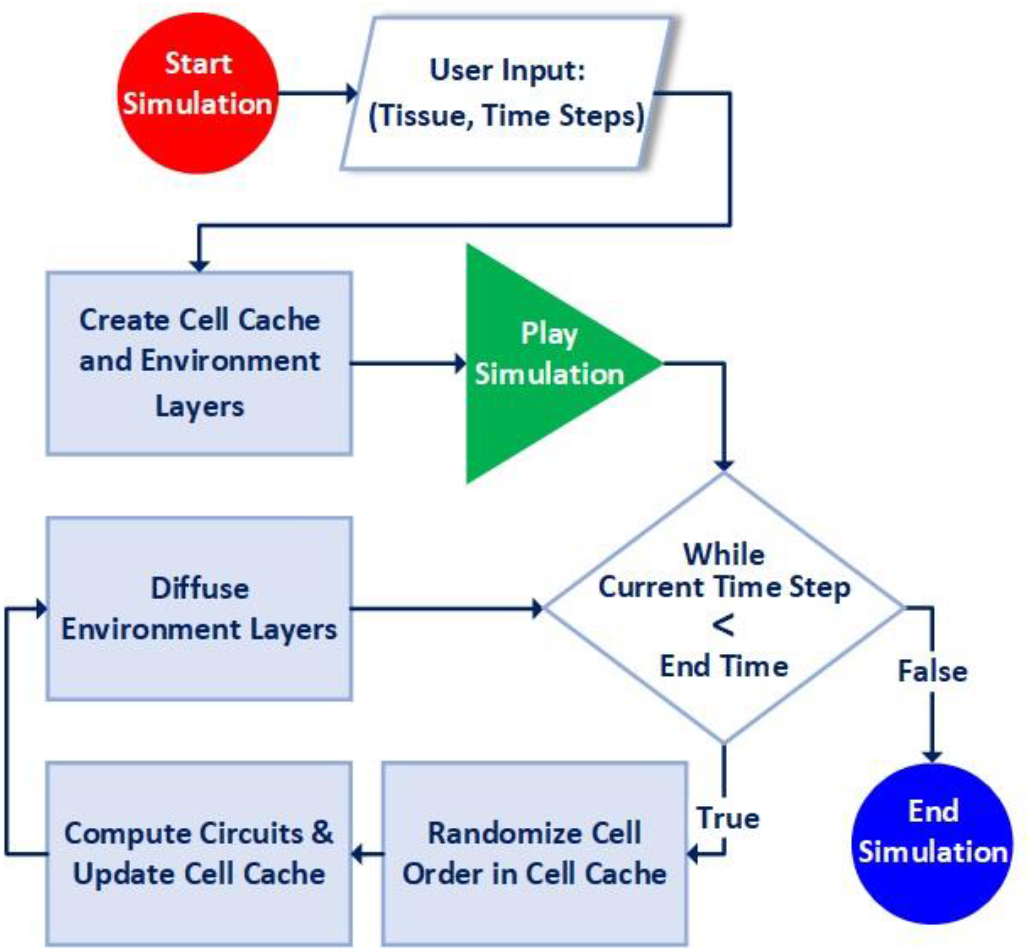
Multi-agent Modeling Engine Flowchart. Users can start the simulation by selecting the tissue and setting up the total time steps in the simulation. Next, cell cache and environment layers are created. The simulation is played and cell order is randomized in the cell cache. The cell circuits for each cell are computed and the cell cache is updated simultaneously. This is followed by environmental layer diffusion. The process is repeated at each time step and the resulting data is stored. This process iterates until the final time step of the simulation.

This work also exemplifies the reconstruction of several published case studies. The data generated from these case studies helped decode mechanisms driving the evolution of heterogeneous tumors which is currently difficult to undertake in wet lab settings. Newer case studies developed using TISON can revolutionize the pharmaceutical and industrial landscape by assisting in the identification and evaluation of personalized therapeutics in the light of patient-specific gene expression and mutation data.

In 2020, Gondal *et al*. (82) employed TISON’s Networks and Therapeutics Editors to construct five Boolean network models of biomolecular regulation in cells lining the *Drosophila* midgut epithelium. These networks were also annotated with colorectal cancer (CRC) patient-specific mutation data to develop an *in silico Drosophila Patient Model* (DPM). This computational framework in the form of an *in silico* DPM was used for decoding CRC along with the development of personalized combinatorial therapeutics for preclinical translational studies. The model was used to evaluate efficacious combinatorial therapies for individual CRC patients towards developing personalized cancer models. The personalized network models helped identify a synergistic combination of paclitaxel and pazopanib for treating colorectal cancer. These network models can also be extended to develop a multi-scale model of CRC towards incorporating spatiotemporal regulations of colorectal cancer using other editors in TISON. Similarly, several other cancer-specific case studies are currently in the process of being developed using networks and multi-scale capabilities in TISON.

Taken together, TISON is a robust and flexible platform for computational modeling and analysis towards developing a systems-level understanding of cellular pathways in the context of tumorigenesis and cancer at biomolecular, cellular, and tissue level. It advances the state-of-the-art in cancer systems oncology and stands to deliver significant advantages to cancer patients, clinicians, and the pharmaceutical industry. Besides the development of personalized therapeutics, the research community stands to gain by obtaining a modeling framework that can save precious material resources and time required for conducting wet-lab experiments. TISON can, therefore, help to decode mechanisms underpinning a variety of tumors, tailor efficacious therapeutic cocktails for cancer patients, elicit novel drug targets and combinations for onward evaluation.

## Materials and Methods

### System Architecture and Software Design

TISON is a web application that has been developed using a distributed three-tier Model-View-Controller (MVC) architecture (83). Microsoft® ASP.NET v5.2.3 MVC framework was employed to implement communication between each tier. The resultant architecture comprises a front-end, middleware, and back-end. The front-end consists of eight interactive web-based Graphical User Interfaces (GUIs) called “*editors*” that utilize Java script’s Jquery framework v1.12.3 (84) for communication with the middleware. Ajax (85) was used for client-server communication. Each front-end editor further utilizes third-party JavaScript libraries including JointJS v1.0.3 (86), PlotlyJS v1.39.2 (87–89), and JSTree v3.3.8 (90), to support GUI, data plotting, and visualization features in TISON. Web pages for each editor were developed using Hypertext Markup Language (HTML 3.0) (91), Cascading Style Sheets (CSS3) (92), and Bootstrap v3.3.7 (93). TISON’s middle-ware comprises of front-end controllers and a multi-agent simulation engine (94) that were both developed using ASP.NET Microsoft Framework v4.6 (95) in C#. ASP.NET Identity v2.2.1 (96) has been employed for user authentication and authorization. Entity Framework v6.1.3 (97) which is an Object Relational Mapper (ORM), was used to map database models. Lastly, the back-end was developed using Microsoft® SQL Server 2014 (98) to store, retrieve and manage data. Icons for GUIs were imported from flaticon.com (99).

### Networks Editor

TISON’s Networks Editor (NE) employs JointJS v1.0.3 (86) to create and draw networks on the canvas. Three types of Boolean network analysis methodologies have been implemented in the NE; (i) Deterministic (54), (ii) Probabilistic (60), and (iii) Ordinary Differential Equation (ODE) analysis (61). The Deterministic Analysis (DA) pipeline assists in analyzing closed systems wherein networks are not subjected to external stimuli or perturbations. Probabilistic Analysis (PA) pipeline caters to open system computations that take into account the intrinsic signaling perturbations in the presence of random noise. The algorithms for DA and PA pipelines have been derived from *ATLANTIS* (100). Results from network analyses can be visualized by using a variety of two-dimensional attractor and cell fate landscapes (51,52,55,61,101). Plotly v1.39.2 (87–89) was used for 2D plotting and visualization while ALGLIB (89) and Math.Numerics (102) were employed for network state matrix manipulation. The Ordinary Differential Equation (ODE) model analysis pipeline has been adopted from *NetLand* (61). To provide ready-support for tissue-specific gene expression data, Metabolic gEne Rapid Visualizer (MERAV) (24), Human Proteome Atlas (HPA) (57), The Cancer Genome Atlas (TCGA) (58) through Firebrowse (59), and The Genotype-Tissue Expression project (GTEx) (103) have been seamlessly integrated into the NE. Math.Lundin (104) was used as an expression parser to compute the Boolean rules for rules-based networks. Conversion from rules-based networks to weight-based networks has been implemented using the scheme reported by Kim *et* al (56).

### Therapeutics Editor

TISON’s Therapeutics Editor (TE) employs JointJS v1.0.3 (86) to visualize networks created earlier in NE. JSTree v3.3.8 (90) library facilitates the therapy construction process in the “Therapeutics Steps” panel. TE incorporates third-party APIs to integrate DGIdb (63) HPA (network DBs), TCGA (patient DBs), and DrugDB (drug DBs) for performing therapeutic evaluations. Network analyses for each therapy can employ DA, PA, and ODE (as described in NE, above). Additionally, Plotly v1.39.2 (87–89) was used to construct and plot stack and bar graphs for comparing analyses results as well as for landscape construction and visualization.

### Environments Editor

TISON’s Environments Editor (EE) implements continuum models of extracellular environments using Partial Differential Equations (PDE) (67). Alternating Direction Implicit (ADI) method (70,105) was used to obtain numerical estimates of the PDE. The tridiagonal matrix generated from the ADI method was solved using the Thomas algorithm (106). Dirichlet boundary condition (107) defines the diffusive behavior of each layer at the edges of the model domain. Plotly v1.54.1 (87–89) library was used for 2D plotting the environmental layers and D3.js (108) was used for their dynamic and interactive data visualization. The layer diffusion matrix’s z-index toggle bar is implemented using Ion.RangeSlider library (109).

### Cell Circuits Editor

TISON’s Cell Circuits Editor (CCE) was developed using the JointJS v1.0.3 (86) library for front-end visualization. Tipped.js (110) and jQuery.ui (111) were employed for front-end designing and parser.js (112) was used as a utility library for object structure conversions. Jsep parser (113) was used to parse inputs with algebraic expressions. To implement back-end information, Mathparser (104) was utilized for solving equations provided by the user. The resultant cell circuits are dynamically validated, computed (114), and then stored as Finite State Machines (FSM) (74,75).

### Cell Lines and Organoids Editor

Cell Lines Editor (CLE) assigns cell circuits developed using CCE to create specific cell types which are later used to created 3D organoids in the Organoids Editor (OE). A one-to-one entity relationship exists between cell circuits and cell lines. OE uses Plotly v1.54.1 (87–89) to draw cells on the specific Cartesian coordinates specified by the user within the world lattice space. jQuery.UI (111) was used to provide drag and drop feature for arranging organoid elements in the editor. Math.js (115) parser was utilized to solve user-provided equations for computing Cartesian coordinates of cells in organoids.

### Simulation Editor

Simulation Editor (SE) was developed using Plotly v1.54.1 (87–89) which helps draw cells on the user-specified Cartesian coordinates within the fixed lattice space (116). The resultant simulation at each time step can be viewed using a slider which is implemented using Ion.RangeSlider library (109). The results are downloaded in zip format using JSZip (117)

### Analytics Editor

Analytics Editor (AE) uses Plotly v1.54.1 (87–89) to plot simulation data in the form of graphs.

### Multi-agent Modeling Engine & Database Design Schema

In TISON, organoids are abstracted as a complex multi-agent system wherein each cell is an independent agent (118,119). At every time step, TISON’s simulation engine computes each cell in the organoid and stores its internal state using a set of variables. Note that the order of cells is randomized before computation. The computation of each cell involves a call to and calculation of the corresponding cell circuit that results in an update of the cell’s internal variables. Several cells can have the same cell circuit to represent the same cellular phenotype. Cells can also switch their cell circuits during a simulation (45,76). Simultaneously, the PDE models of extracellular environments are also computed, if included in the model. The simulation engine is implemented as a Web API that is connected to the front-end editors through corresponding controllers. The engine continues to perform these in tandem computations of cells and environments at each time step, until the time limit set by the user. The tissue organoids state is stored at each time step and is available for analytics at the end of the simulation. The simulation engine works off data models that are stored in a Microsoft SQL Server (98).

## Data and Software Availability

The software, case studies, and manual are freely available at http://tison.lums.edu.pk. The issues reporting and database are catered at https://github.com/BIRL/TISON/issues.

## Acknowledgments

This work was supported by the National ICT-R&D Fund (SRG-209), RF-NCBC-015, NGIRI-2020-4771, HEC (21-30SRGP/R&D/HEC/ 2014, 20-2269/NRPU/R&D/ HEC/12/ 4792 and 20-3629/NRPU/R&D/HEC/14/ 585), TWAS (RG 14-319 RG/ITC/AS_C) and LUMS (STG-BIO-1008, FIF-BIO-2052, FIF-BIO-0255, SRP-185-BIO, SRP-058-BIO and FIF-477-1819-BIO) grants.

## Author contributions

SUC designed and supervised the project. MNG led the project development and manuscript team. MNG, MUS, AR, AA, HAA, ZA, MFAC, WA, SK, FA, MNJ, HH, and MFAB implemented the software. MNG, RH, HK, SK, BA, RNB, RA, ZN, OSS, and MMA designed case studies and developed software manuals and videos. SUC, MNG, RH, HK, SK, BA, RNB, RA, and ZN developed the manuscript. SUC, SA, FA, OI, SWN, WV, BW, HS, EU, MS, IJ, MT, and AF revised the case studies and the manuscript.

## References

1. Rejniak KA, Anderson ARA. Hybrid models of tumor growth. Wiley Interdiscip Rev Syst Biol Med. 2011;3(1):115–25.

2. Izaguirre JA, Chaturvedi R, Huang C, Cickovski T, Coffland J, Thomas G, et al. CompuCell, a multi-model framework for simulation of morphogenesis. Bioinformatics. 2004;20(7):1129–37.

3. Yoshida A, Kohyama S, Fujiwara K, Nishikawa S, Doi N. Regulation of spatiotemporal patterning in artificial cells by a defined protein expression system. Chem Sci. 2019;11064–72.

4. Yamada T, Bork P. Evolution of biomolecular networks — lessons from metabolic and protein interactions. 2009;10(NovEmBER). Available from: http://dx.doi.org/10.1038/nrm2787

5. Liebert MA. Using Bayesian Networks to Analyze Expression Data. 2000;7:601– 20.

6. Wang E, Lenferink A, O’Connor-McCourt M. Genetic studies of diseases - Cancer systems biology: Exploring cancer-associated genes on cellular networks. Cell Mol Life Sci. 2007;64(14):1752–62.

7. Chalancon G, Ravarani CNJ, Balaji S, Martinez-Arias A, Aravind L, Jothi R, et al. Interplay between gene expression noise and regulatory network architecture. Trends Genet [Internet]. 2012;28(5):221–32. Available from: http://www.sciencedirect.com/science/article/pii/S0168952512000157

8. Bryant DM, Mostov KE. o E From cells to organs?: building polarized tissue. 2008;9(NovEMbER).

9. O’Brien LE, Zegers MMP, Mostov KE. Building epithelial architecture: insights from three-dimensional culture models. Nat Rev Mol Cell Biol [Internet]. 2002;3(7):531–7. Available from: https://doi.org/10.1038/nrm859

10. Bindea G, Mlecnik B, Tosolini M, Kirilovsky A, Waldner M, Obenauf AC, et al. Spatiotemporal Dynamics of Intratumoral Immune Cells Reveal the Immune Landscape in Human Cancer. Immunity [Internet]. 2013;39(4):782–95. Available from: http://www.sciencedirect.com/science/article/pii/S1074761313004378

11. Maestú F, Fernández A, Simos PG, Gil-Gregorio P, Amo C, Rodriguez R, et al. Spatio-temporal patterns of brain magnetic activity during a memory task in Alzheimer’s disease. Neuroreport. 2001;12(18):3917–22.

12. Duong Van Huyen J-P, Amri K, Bélair M-F, Vilar J, Merlet-Bénichou C, Bruneval P, et al. Spatiotemporal distribution of insulin-like growth factor receptors during nephrogenesis in fetuses from normal and diabetic rats. Cell Tissue Res [Internet]. 2003;314(3):367–79. Available from: https://doi.org/10.1007/s00441-003-0803-4

13. Chakraborty S, Rahman T. The difficulties in cancer treatment. Ecancermedicalscience. 2012;6:ed16.

14. Patel LR, Camacho DF, Shiozawa Y, Pienta KJ, Taichman RS. Mechanisms of cancer cell metastasis to the bone: a multistep process. Futur Oncol. 2011;7(11):1285–97.

15. Strauss R, Li Z-Y, Liu Y, Beyer I, Persson J, Sova P, et al. Analysis of Epithelial and Mesenchymal Markers in Ovarian Cancer Reveals Phenotypic Heterogeneity and Plasticity. PLoS One [Internet]. 2011;6(1):1–20. Available from: https://doi.org/10.1371/journal.pone.0016186

16. Mazio C, Casale C, Imparato G, Urciuolo F, Netti PA. Recapitulating spatiotemporal tumor heterogeneity in vitro through engineered breast cancer microtissues. Acta Biomater [Internet]. 2018;73:236–49. Available from: http://www.sciencedirect.com/science/article/pii/S1742706118302319

17. Archer TC, Fertig EJ, Gosline SJC, Hafner M, Hughes SK, Joughin BA, et al. Systems approaches to cancer biology. Cancer Res. 2016;76(23):6774–7.

18. Heiner M, Gilbert D. BioModel engineering for multiscale Systems Biology. Prog Biophys Mol Biol [Internet]. 2012;1–10. Available from: http://dx.doi.org/10.1016/j.pbiomolbio.2012.10.001

19. Manzoni C, Kia DA, Vandrovcova J, Hardy J, Wood NW, Lewis PA, et al. Genome, transcriptome and proteome: the rise of omics data and their integration in biomedical sciences. Brief Bioinform [Internet]. 2016;19(2):286–302. Available from: https://doi.org/10.1093/bib/bbw114

20. Stamatakos GS, Dionysiou DD, Graf NM, Sofra NA, Desmedt C, Hoppe A, et al. The “Oncosimulator”: a multilevel, clinically oriented simulation system of tumor growth and organism response to therapeutic schemes. Towards the clinical evaluation of in silico oncology. Conf Proc IEEE Eng Med Biol Soc. 2007;6629–32.

21. Bamford S, Dawson E, Forbes S, Clements J, Pettett R, Dogan A, et al. The COSMIC (Catalogue of Somatic Mutations in Cancer) database and website. Br J Cancer. 2004;91:355–8.

22. Lonsdale J, Thomas J, Salvatore M, Phillips R, Lo E, Shad S, et al. The Genotype-Tissue Expression (GTEx) project. Vol. 45, Nature Genetics. Nature Publishing Group; 2013. p. 580–5.

23. Lee HJ, Palm J, Grimes SM, Ji HP. The Cancer Genome Atlas Clinical Explorer: A web and mobile interface for identifying clinical-genomic driver associations. Genome Med [Internet]. 2015;7(1):1–14. Available from: http://dx.doi.org/10.1186/s13073-015-0226-3

24. Shaul YD, Yuan B, Thiru P, Nutter-Upham A, McCallum S, Lanzkron C, et al. MERAV: A tool for comparing gene expression across human tissues and cell types. Nucleic Acids Res. 2016;44(D1):D560–6.

25. Li J, Akbani R, Zhao W, Lu Y, Weinstein JN, Mills GB, et al. Explore, visualize, and analyze functional cancer proteomic data using the cancer proteome atlas. Cancer Res. 2017;77(21):e51–4.

26. Thul PJ, Lindskog C. The human protein atlas: A spatial map of the human proteome. Protein Sci. 2018;27(1):233–44.

27. Wishart DS, Feunang YD, Marcu A, Guo AC, Liang K, Vázquez-Fresno R, et al. HMDB 4.0: the human metabolome database for 2018. Nucleic Acids Res. 2018 Jan;46(D1):D608–17.

28. Powathil GG, Swat M, Chaplain MAJ. Systems oncology: Towards patient-specific treatment regimes informed by multiscale mathematical modelling. Semin Cancer Biol [Internet]. 2015;30:13–20. Available from: http://www.sciencedirect.com/science/article/pii/S1044579X14000273

29. Valladares-Ayerbes M, Haz-Conde M, Blanco-Calvo M. Systems oncology: toward the clinical application of cancer systems biology. Futur Oncol. 2015;11(4):553–5.

30. Anderson ARA, Rejniak KA, Gerlee P, Quaranta V. Microenvironment driven invasion: A multiscale multimodel investigation. J Math Biol. 2009;58(4–5):579– 624.

31. Stolarska MA, Yangjin KIM, Othmer HG. Multi-scale models of cell and tissue dynamics. Philos Trans R Soc A Math Phys Eng Sci. 2009;367(1902):3525–53.

32. Silva A, Anderson A, Gatenby R. A multiscale model of the bone marrow and hematopoiesis. Math Biosci Eng. 2011;8(2):643–58.

33. Chaudhary SU, Shin S, Won J, Cho K, Member S. Multiscale Modeling of Tumorigenesis Induced by Mitochondrial Incapacitation in Cell Death. IEEE Trans Biomed Eng. 2011;58(2):3028–32.

34. Barbarroux L, Michel P, Adimy M, Crauste F. Multi-scale modeling of the CD8 immune response. AIP Conf Proc. 2016;1738.

35. Szabó A, Merks RMH. Blood vessel tortuosity selects against evolution of aggressive tumor cells in confined tissue environments: A modeling approach. Vol. 13, PLoS Computational Biology. 2017. 1–32 p.

36. Kumar S, Das A, Sen S. Multicompartment cell-based modeling of confined migration: regulation by cell intrinsic and extrinsic factors. Mol Biol Cell. 2018;29(13):1599–610.

37. Unni P. Mathematical Modeling, Analysis, and Simulation of Tumor. 2019;2019.

38. Gerlee P, Anderson ARA. An evolutionary hybrid cellular automaton model of solid tumour growth. J Theor Biol. 2007 Jun;246(4):583–603.

39. Fletcher AG, Murray PJ, Maini PK. Multiscale modelling of intestinal crypt organization and carcinogenesis. Math Model Methods Appl Sci. 2015;25(13):2563–85.

40. Shamsi M, Saghafia M, Dejam M, Sanati-nezhad A. Mathematical Modeling of the Function of Warburg Effect in Tumor Microenvironment. 2018;(May):1–13.

41. Ion I. Moraru, James C. Schaff, Boris M. Slepchenko, Michael Blinov, Frank Morgan, Anuradha Lakshminarayana, Fei Gao, Ye Li and LML. The Virtual Cell Modeling and Simulation Software Environment. PMC. 2008;

42. Eissing T, Kuepfer L, Becker C, Block M, Coboeken K, Gaub T, et al. A computational systems biology software platform for multiscale modeling and simulation: Integrating whole-body physiology, disease biology, and molecular reaction networks. Front Physiol. 2011;FEB(February):1–10.

43. Hoops S, Gauges R, Lee C, Pahle J, Simus N, Singhal M, et al. COPASI - A COmplex PAthway SImulator. Bioinformatics. 2006;22(24):3067–74.

44. Metzcar J, Wang Y, Heiland R, Macklin P. A Review of Cell-Based Computational Modeling in Cancer Biology. JCO Clin Cancer Informatics. 2019 Nov;3(3):1–13.

45. Chaudhary SU, Shin S-Y, Lee D, Song J-H, Cho K-H. ELECANS--an integrated model development environment for multiscale cancer systems biology. Bioinformatics. 2013;29(7):957–9.

46. Mirams GR, Arthurs CJ, Bernabeu MO, Bordas R, Cooper J, Corrias A, et al. Chaste: An Open Source C++ Library for Computational Physiology and Biology. PLoS Comput Biol. 2013;9(3).

47. Prestes A, Alfonso G. Sensitivity analysis of Repast computational ecology models with R / Repast. 2016;(June):8811–31.

48. Phan JH, Quo CF, Cheng C, Wang MD. Multiscale integration of -omic, imaging, and clinical data in biomedical informatics. IEEE Rev Biomed Eng. 2012;5:74–87.

49. Chow TS. Testing software design modeled by finite-state machines. IEEE Trans Softw Eng. 1978;(3):178–87.

50. Cho S-H, Park S-M, Lee H-S, Lee H-Y, Cho K-H. Attractor landscape analysis of colorectal tumorigenesis and its reversion. BMC Syst Biol. 2016;10(1):96.

51. Choi M, Shi J, Jung SH, Chen X, Cho K-H. Attractor Landscape Analysis Reveals Feedback Loops in the p53 Network That Control the Cellular Response to DNA Damage. Sci Signal [Internet]. 2012;5(251):ra83.-ra83. Available from: https://stke.sciencemag.org/content/5/251/ra83

52. Han B, Wang J. Quantifying robustness and dissipation cost of yeast cell cycle network: the funneled energy landscape perspectives. Biophys J. 2007;92(11):3755–63.

53. Li C, Wang J. Quantifying Cell Fate Decisions for Differentiation and Reprogramming of a Human Stem Cell Network?: Landscape and Biological Paths. 2013;9(8).

54. Glass L, Kauffman SA. The logical analysis of continuous, non-linear biochemical control networks. J Theor Biol. 1973 Apr;39(1):103–29.

55. Huang S, Ernberg I, Kauffman S. Cancer attractors: A systems view of tumors from a gene network dynamics and developmental perspective. Semin Cell Dev Biol [Internet]. 2009;20(7):869–76. Available from: http://www.sciencedirect.com/science/article/pii/S1084952109001499

56. Kim Y, Choi S, Shin D, Cho KH. Quantitative evaluation and reversion analysis of the attractor landscapes of an intracellular regulatory network for colorectal cancer. BMC Syst Biol. 2017;11(1):1–22.

57. Uhlén M, Fagerberg L, Hallström BM, Lindskog C, Oksvold P, Mardinoglu A, et al. Tissue-based map of the human proteome. Science (80-). 2015;347(6220).

58. Zhang K, Wang H. Cancer Genome Atlas Pan-cancer analysis project. Chinese J Lung Cancer. 2015;18(4):219–23.

59. Kryukov I. Original article FirebrowseR?: an R client to the Broad Institute ‘ s Firehose Pipeline. 2017;1–6.

60. Shmulevich I, Dougherty ER, Kim S, Zhang W. Probabilistic Boolean networks: a rule-based uncertainty model for gene regulatory networks. Bioinformatics. 2002;18(2):261–74.

61. Guo J, Lin F, Zhang X, Tanavde V, Zheng J. NetLand: Quantitative modeling and visualization of Waddington’s epigenetic landscape using probabilistic potential. Bioinformatics. 2017;33(10):1583–5.

62. Kitano H. Biological robustness. Nat Rev Genet [Internet]. 2004;5(11):826–37. Available from: https://doi.org/10.1038/nrg1471

63. Cotto KC, Wagner AH, Feng Y-Y, Kiwala S, Coffman AC, Spies G, et al. DGIdb 3.0: a redesign and expansion of the drug–gene interaction database. Nucleic Acids Res. 2018;46(D1):D1068–73.

64. Warrick JW, Murphy WL, Beebe DJ. Screening the cellular microenvironment: a role for microfluidics. IEEE Rev Biomed Eng. 2008;1(1):75–93.

65. Whiteside TL. The tumor microenvironment and its role in promoting tumor growth. Oncogene. 2008 Oct;27(45):5904–12.

66. Heddleston JM, Li Z, McLendon RE, Hjelmeland AB, Rich JN. The hypoxic microenvironment maintains glioblastoma stem cells and promotes reprogramming towards a cancer stem cell phenotype. Cell cycle. 2009;8(20):3274–84.

67. Villadsen J, Michelsen ML. Solution of differential equation models by polynomial approximation. Vol. 7. Prentice-Hall Englewood Cliffs, NJ; 1978.

68. Cheng AHD, Cheng DT. Heritage and early history of the boundary element method. Eng Anal Bound Elem. 2005;29(3):268–302.

69. Chang MJ, Chow LC, Chang WS. Improved alternating-direction implicit method for solving transient three-dimensional heat diffusion problems. Numer Heat Transf Part B Fundam. 1991;19(1):69–84.

70. 3-D Heat Equation Numerical Solution [Internet]. Available from: https://www.mathworks.com/matlabcentral/fileexchange/59336-3-d-heat-equation-numerical-solution

71. Comsol AB. COMSOL multiphysics reference manual. Version. 2007;

72. Rivera MJ, Molina JAL, Trujillo M, Romero-García V, Berjano EJ. Analytical validation of COMSOL Multiphysics for theoretical models of Radiofrequency ablation including the Hyperbolic Bioheat transfer equation. In: 2010 Annual International Conference of the IEEE Engineering in Medicine and Biology. IEEE; 2010. p. 3214–7.

73. Mohri M. On some applications of finite-state automata theory to natural language processing. Nat Lang Eng. 1996;2(1):61–80.

74. Chow TS. Testing Software Design Modeled by Finite-State Machines. IEEE Trans Softw Eng. 1978;SE-4(3):178–87.

75. Li JH, Dai GX, Li HH. Mutation analysis for testing finite state machines. 2nd Int Symp Electron Commer Secur ISECS 2009. 2009;1:620–4.

76. Cho K-H, Chaudhary SU, Lee D. Biosimulation method and computing device with high expandability. Korea; Korea Patent No. 10-2013-0033839, 2013.

77. Ignatenko NA, Yerushalmi HF, Pandey R, Kachel KL, Stringer DE, Marton LJ, et al. Gene expression analysis of HCT116 colon tumor-derived cells treated with the polyamine analog PG-11047. Cancer Genomics Proteomics. 2009;6(3):161– 75.

78. Vantangoli MM, Madnick SJ, Huse SM, Weston P, Boekelheide K. MCF-7 Human Breast Cancer Cells Form Differentiated Microtissues in Scaffold-Free Hydrogels. PLoS One [Internet]. 2015 Aug 12;10(8):e0135426–e0135426. Available from: https://pubmed.ncbi.nlm.nih.gov/26267486

79. Wooley JC, Lin HS, Council NR. Computational modeling and simulation as enablers for biological discovery. In: Catalyzing inquiry at the interface of computing and biology. National Academies Press (US); 2005.

80. Li XL, Oduola WO, Qian L, Dougherty ER. Integrating Multiscale Modeling with Drug Effects for Cancer Treatment. Cancer Inform. 2015;14(Suppl 5):21–31.

81. Cosentino DGBC. Validation and invalidation of systems biology models using robustness analysis. 2011;(August 2010):229–44.

82. Gondal MN, Butt RN, Shah OS, Nasir Z, Hussain R, Khawar H, et al. In silico Drosophila Patient Model Reveals Optimal Combinatorial Therapies for Colorectal Cancer. bioRxiv [Internet]. 2020; Available from: https://www.biorxiv.org/content/early/2020/09/01/2020.08.31.274829

83. Gupta P, Govil MC. MVC Design Pattern for the multi framework distributed applications using XML, spring and struts framework. Int J Comput Sci Eng. 2010;2(04):1047–51.

84. Bibeault B, Kats Y. jQuery in Action. Dreamtech Press; 2008.

85. Khosravi S. ASP. NET AJAX: PROGRAMMER’S REFERENCE. John Wiley & Sons; 2007.

86. Joint API [Internet]. Available from: https://resources.jointjs.com/docs/jointjs/v3.1/joint.html

87. Sievert C, Parmer C, Hocking T, Chamberlain S, Ram K, Corvellec M, et al. plotly: Create Interactive Web Graphics via ‘plotly. js.’ R Packag version. 2017;4(1):110.

88. Ullrich A, Eckelmann F, Ghozzi S. Dashboards as strategy to integrate multiple data streams for real time surveillance. Online J Public Health Inform. 2019;11(1):3–5.

89. ALGLIB [Internet]. Available from: https://www.alglib.net/

90. jsTree [Internet]. Available from: https://www.jstree.com/docs/config/

91. Introduction to HTML 3.0 [Internet]. Available from: https://www.w3.org/MarkUp/html3/intro.html

92. CSS: Cascading Style Sheets [Internet]. Available from: https://developer.mozilla.org/en-US/docs/Web/CSS

93. Mark Otto JT, Rebert C, Thilo J, Xhmikos FH, Lauke PH. Bootstrap Release 3.3. 7; 2016.

94. Sycara KP. Multiagent systems. AI Mag. 1998;19(2):79.

95. Troelsen A, Japikse P. C# 6.0 and the. NET 4.6 Framework. Apress; 2015.

96. Rastogi P, Anderson R, Dykstra T, Galloway J. Introduction to ASP .NET Identity. Microsofts,[Online] Tillgänglig http://www.aspnet/identity/overview/gettingstarted/introduction-to-aspnet-identity[Hämtad2015-02-27]. 2013;

97. Troelsen A, Japikse P. ADO. NET Part III: Entity Framework. In: C# 60 and the NET 46 Framework. Springer; 2015. p. 929–99.

98. Mistry R, Misner S. Introducing Microsoft SQL Server 2014. Microsoft Press; 2014.

99. Flaticon [Internet]. Available from: flaticon.com

100. Shah OS, Chaudhary MFA, Awan HA, Fatima F, Arshad Z, Amina B, et al. ATLANTIS - Attractor Landscape Analysis Toolbox for Cell Fate Discovery and Reprogramming. Sci Rep [Internet]. 2018;8(1):1–11. Available from: http://dx.doi.org/10.1038/s41598-018-22031-3

101. Zheng D, Yang G, Li X, Wang Z, Liu F, He L. An Efficient Algorithm for Computing Attractors of Synchronous And Asynchronous Boolean Networks. PLoS One. 2013;8(4):1–7.

102. Math.NET Numerics [Internet]. Available from: https://numerics.mathdotnet.com/

103. Carithers LJ, Moore HM. The Genotype-Tissue Expression (GTEx) Project. Biopreserv Biobank. 2015;13(5):307–8.

104. Lundin P. Math Parser. 2016.

105. Wang T-Y, Chen CC-P. Thermal-ADI-a linear-time chip-level dynamic thermal-simulation algorithm based on alternating-direction-implicit (ADI) method. IEEE Trans very large scale Integr Syst. 2003;11(4):691–700.

106. Zhang R. Applied Contaminant Transport Modeling: Theory and Practice. Vol. 25, Journal of Environment Quality.1996. p. 927.

107. Maini PK, Myerscough MR. Boundary-driven instability. Appl Math Lett. 1997;10(1):1–4.

108. D3.js [Internet]. Available from: https://d3js.org/

109. Ion.RangeSlider [Internet]. Available from: http://ionden.com/a/plugins/ion.rangeSlider/index.html

110. Tipped.js [Internet]. Available from: https://www.tippedjs.com/

111. jQuery. Available from: https://jqueryui.com/

112. parse-js [Internet]. Available from: https://www.npmjs.com/package/parse-js

113. jsep parser [Internet]. Available from: https://github.com/soney/jsep

114. Cho KH, Chaudhary SU, Lee D. Biosimulation method and computing device using compiler embedded there within. South Korea: Korea Patent Office; 10-2013–0033837, 2013.

115. math.js. Available from: https://mathjs.org/docs/expressions/parsing.html

116. Trahan CJ, Wyatt RE. Classical and quantum phase space evolution: fixed-lattice and trajectory solutions. Chem Phys Lett [Internet]. 2004;385(3):280–5. Available from: http://www.sciencedirect.com/science/article/pii/S0009261403021833

117. JSZip [Internet]. Available from: https://stuk.github.io/jszip/

118. Uhrmacher AM, Weyns D. Multi-Agent systems: Simulation and applications. CRC press; 2009.

119. Khan S, Makkena R, McGeary F, Decker K, Gillis W, Schmidt C. A multi-agent system for the quantitative simulation of biological networks. In: Proceedings of the second international joint conference on Autonomous agents and multiagent systems. 2003. p. 385–92.

