## Supplementary Information for "TISON: a next-generation multi-scale modeling theatre for *in silico* systems oncology"

[1] Biomedical Informatics Research Laboratory, Department of Biology, Lahore University of Management Sciences, Lahore, Pakistan, [2] Department of Biostatistics and Bioinformatics, School of Medicine, Duke University, Durham, USA, [3] The Roy J. Carver Department of Biomedical Engineering, The University of Iowa, Iowa City, USA, [4] Department of Pathology, Shaukat Khanum Memorial Cancer Hospital and Research Centre, Lahore, Pakistan, [5] Department of Statistics, University of Gujrat, Gujrat, Pakistan, [6] Department of Computer Science, National University of Computer and Emerging Sciences, Islamabad, Pakistan, [7] School of Computing Science, University of Glasgow, Glasgow, United Kingdom, [8] University of Management and Technology, Lahore, Pakistan, [9] Sabz Qalam, Lahore, Pakistan, [10] National University of Medical Sciences, Islamabad, Pakistan, [11] National University of Science and Technology, Islamabad, Pakistan, [12] Pakistan Institute of Engineering and Applied Sciences, Islamabad, Pakistan, and [13] Department of Biology, Lahore University of Management Sciences, Lahore, Pakistan.

* Corresponding author. Department of Biology, Lahore University of Management Sciences, DHA Phase 5, Opposite Sector U, Khayaban e Jinnah، Road، Cantt، Lahore, Punjab 54792 Lahore, Pakistan. Tel.: +92 321 4255171; Fax: +92 42 3560 8317;

### Section 1. Overview of the Platform

Theater for *in silico* Systems Oncology (TISON) is a web application that provides its users with an intuitive “zero-code” multi-scale model development environment for application in cancer systems biology. The platform is freely available as an online portal where users can create or upload their biological models of biomolecular networks, perform therapeutic screens, construct *in silico* cellular phenotypes, and multi-cellular organoids for onward spatiotemporal simulation and data analysis. Users can develop these models by creating a project in TISON followed by setting up a “world size” which allows users to model up to 1 million cells using fixed lattice (1) space.

The platform is designed using a three-tier software architecture consisting of front-end, middleware, and back-end. The front-end hosts information on web clients and consists of eight web-based graphical user interfaces (GUI) termed *“editors”*. These editors are interconnected, and each editor provides scale-specific modeling features towards the stepwise development of complex systems oncology models. The resulting models are taken up by the middle-ware comprising of front-end controllers and a high-performance multi-agent simulation engine. Editors pass information to the corresponding controllers and models in the middleware through a secure API. This information comprising of model properties and associated user-defined parameters is utilized by the controllers to generate simulation logic and compute results. The results are then passed on to the database at the back-end. The back-end is therefore used to store, retrieve, and manage model data. The information contained is then serialized and de-serialized at the back-end for provision to middle-ware for onwards processing.

### Section 2. Modeling Editors in TISON

TISON is available to the users as an online portal wherein they can freely register to create, upload, simulate, or analyze their multi-scale models in the form of projects. TISON envisages an intuitive interface for the construction of multi-scale models with the provision of eight scale-specific editors. These editors have been designed following the biological structural organization, starting from a genomic scale up to the cell lines and organoids scales. Multi-scale model development in TISON is performed by using one or more of its eight scale-specific editors (Figure S1).

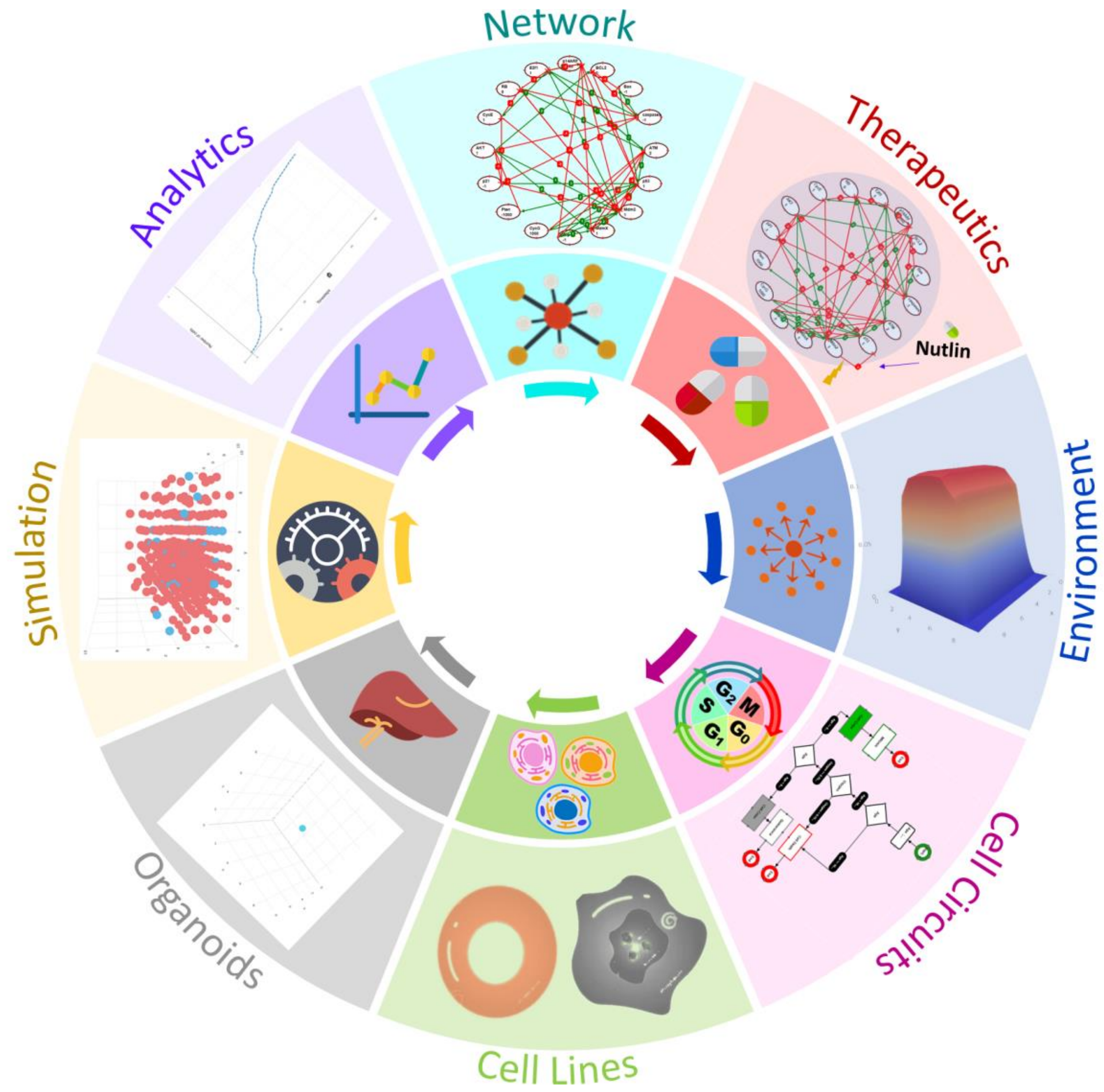

**Figure S1- Schematics of Information Flow between TISON Editors.** TISON's front-end comprises 8 editors for stepwise multi-scale model development. These include (1) networks editor to design biomolecular regulatory networks, (2) therapeutics editor for the mutational and therapeutic screening of biomolecular networks, (3) environments editor to model extracellular environments for providing inputs to biomolecular networks, (4) cell circuits editor to create finite state machines which integrate biomolecular networks, therapies and extracellular networks into cell fate decision flowcharts, (5) cell lines editor to assign cell circuits to phenotypes, (6) organoids editor to design organoids, (7)

simulations editor to simulate organoid models for studying their temporal development, and (8) analytics editor to perform quantitative analysis of simulation data.

These editors include (i) Networks Editor (NE) for biomolecular network construction and analysis, (ii) Therapeutics Editor (TE) for designing therapeutic interventions, (iii) Environments Editor (EE) for creating tumor microenvironments, (vi) Cell Circuits Editor (CCE) to design cell decision circuits, (v) Cell Lines Editor (CLE) for cell circuit assignment to cell types, (vi) Organoids Editor (OE) for assembling three-dimensional organoids from cell lines, (vii) Simulations Editor (SE) to undertake spatiotemporal simulation of organoids, and lastly, (viii) Analytics Editor (AE) for analysis of simulation results.

In the following sub-sections, a description of software features and functionalities, software development methodology, as well as sample case studies are provided for each editor to assist TISON users in the employment of the platform.

### Section 2.1. Networks Editor – An Overview

#### Section 2.1.1. Software Features & Functionalities

Users can initiate the multi-scale model development process in TISON by building biomolecular interaction networks of biomolecules such as genes and proteins in the Networks Editor (NE). Users can enter Networks Editor from the Project Home Page or the toolbar at the bottom of the graphical user interface (GUI) after opening a project.

- **Creating Networks**

NE assists in the construction of biomolecular interaction networks in multiple modalities. Users can draw weight-based networks by dragging and dropping elements such as genes, proteins, ligands, and receptors from the editor's toolbar onto the drawing canvas. Rules-based networks can be created by writing rules in the rules editor. Weight-based and rules-based networks can also be created by uploading CSV (comma separated values) files containing the network. Both types of networks can be exported from or imported into the NE. Networks can be merged, divided into sub-networks, zoomed-in, and duplicated. Additionally, NE allows the seamless conversion of rules-based networks into weight-based networks (2). Expression data from online databases can be integrated into network models by creating nodes from databases. This expression can be employed to calculate the basal expression for weight-based networks using an in-built basal value calculation feature in NE.

To further assist the users and to validate the functioning of the editor, four biomolecular network case studies have been provided which can be conveniently loaded for use as exemplars (see Section 2.1.3 for NE's User Manual). These case studies can be accessed through the "Upload Case Study" button in NE or through the template projects tab on the TISON's home page (see Section 2.1.3 for NE's User Manual).

- **Analyzing Networks**

Once a network has been created in NE, three types of network analyses are available depending on the network. Deterministic Analysis (DA) can be performed on rules-based networks whereas, DA, Probabilistic Analysis (PA), and Ordinary Differential Equation (ODE) analysis can be performed on weight-based networks. Input files containing one or more network inputs can be consumed by NE and analysis results can be viewed or downloaded by the user. Analysis results are available in both summary as well as detailed formats depending on the type of network and analysis (see Section 2.1.3 for NE's User Manual).

- **Visualizing Networks**

Depending on the type of network being analyzed, results can be viewed as cell fate landscapes, attractor landscapes, potential energy landscapes, probability landscapes, and ODE landscapes. Furthermore, a summary of analysis results obtained using combinatorial or batch inputs can be visualized in the form of line graphs (for row-wise batch inputs) and bar charts (for combinatorial inputs) (see Section 2.1.3 for NE's User Manual).

#### Section 2.1.2. Functional Validation of the Networks Editor

The network modeling and analysis features of TISON's NE were evaluated by reconstructing four published case studies including (i) Rules-Based Human Signaling Network for Deterministic Analysis (DA) (3), (ii) Weight-Based p53 Network for DNA damage ON and OFF for DA (4), (iii) Weight-Based Yeast Cell Cycle Network for Probabilistic Analysis (PA) (5) and, (iv) Weight-Based Stem Cell Network for Ordinary Differential Equation (ODE) Analysis (6,7). Results for deterministic and probabilistic analyses, attractor landscapes, and basin ratios were compared with the published literature for each of these case studies. Supplementary results from each of the aforementioned case studies are provided below.

#### Case Study 1 – Investigation of Colorectal Tumorigenesis in Human Signaling Network

In the first case study, we reconstructed the network employed by Cho *et al.* for investigating colorectal tumorigenesis in a Human Signaling Network (3). The network used Boolean rules and contained 197 nodes and 744 edges (Figure S2A). The cell fate outputs included apoptosis, metastasis, and proliferation. TISON's NE was used to perform deterministic analysis on the network under control conditions (see Supplementary Data – NE for further details).

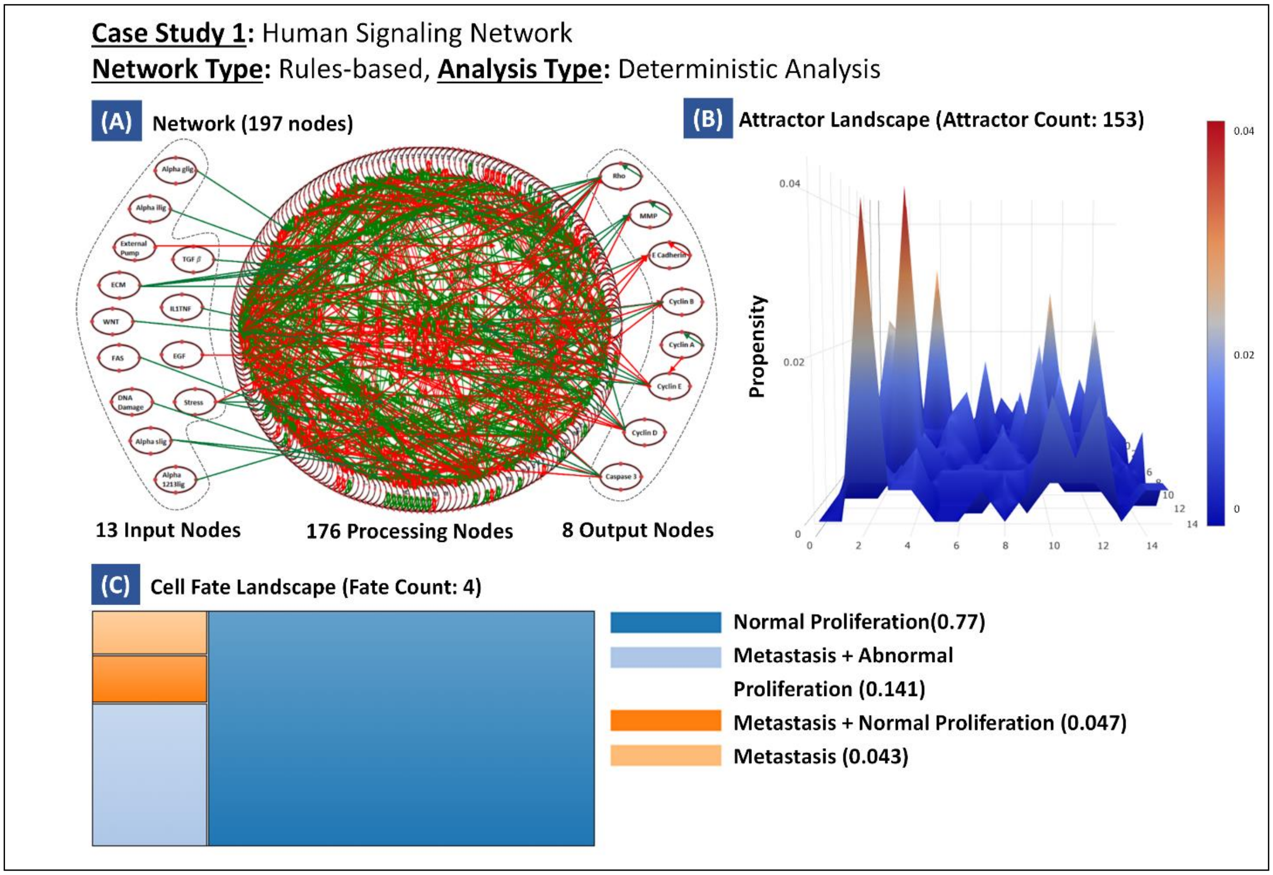

**Figure S2 – Case Study 1: Human Signaling Network Analysis under Control Conditions.** (A) The network consisting of 197 nodes and 744 edges with 13 input nodes, 8 output nodes, and 176 processing nodes constructed using TISON's NE, (B) Attractor

Landscape with 153 cell fate attractors, and (C) Cell Fate Landscape representing four most likely attractors.

- **Analysis Results**

Network analysis of the human signaling network generated 153 attractors (Figure S2B), corresponding to the most probable network steady states, which were then mapped onto cell fates. The cell fate landscape exhibited four major attractors in response to the normal input conditions (Figure S2C) (i) normal proliferation with a propensity of 0.77 (ii) metastasis and abnormal proliferation with a propensity of 0.141 (iii) metastasis and normal proliferation with a propensity of 0.047, and (iv) metastasis with a propensity of 0.043. These results were in agreement with those reported by Cho *et al.* (see Supplementary Data – NE for further details).

#### Case Study 2 – p53-mediated Apoptosis Network in MCF-7 Breast Cancer Cell Lines

The second case study reconstructed a p53 network to investigate the cellular response to DNA damage, performed by Choi *et al.* (4). The network had 16 nodes and 50 edges. TISON was employed to compute the cell fate propensities after the introduction of DNA damage, using DA. DNA damage ON and OFF states was created by setting the basal value of ATM node; -1 (Figure S3A) for DNA damage OFF and 2 for DNA damage ON (Figure S4A) (see Supplementary Data – NE for further details).

- **Analysis Results**

The weight-based deterministic analysis was employed to model the dynamics of the p53 network (Figure S5A) in response to DNA damage ON and OFF. The results had one attractor state with a basin size ratio of 1 (Figure S3B) in response to DNA damage OFF. This corresponded to proliferation in the cell fate landscape (Figure S3C and Figure S4). For DNA damage ON, DA showed the emergence of one cell cycle arrest attractor (Figure S5B and Figure S6). Hence, NE correctly reproduced the attractor states for both DNA damage ON and OFF as reported by Choi *et al.* (see Supplementary Data – NE for further details).

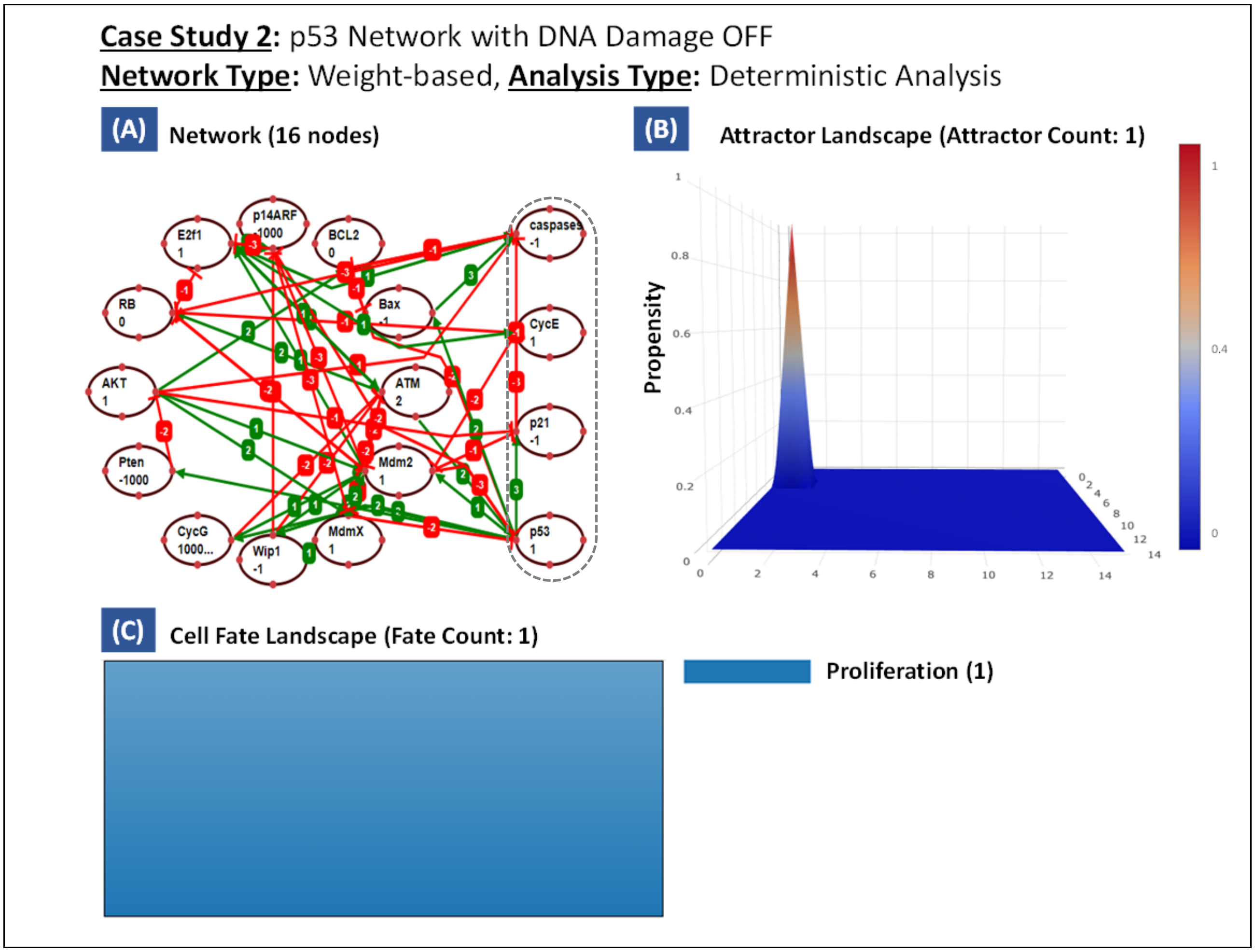

**Figure S3 – Case Study 2: p53 Network with DNA Damage OFF.** (A) The network containing 16 nodes and 50 edges with 4 output nodes and 12 processing nodes constructed using TISON's NE, (B) Attractor Landscape with 1 cell fate attractor, and (C) Cell Fate Landscape with 1 cell fate representing the most probable attractor.

| Deterministic Analysis Results | | | | | | | | | | | | | | | | | | | |
|---|---|---|---|---|---|---|---|---|---|---|---|---|---|---|---|---|---|---|---|
| Attractor | Associated Cell Fate | Basin Size Ratio | Attractor Type | atm | p53 | mdm2 | mdmx | wip1 | cycg | pten | p21 | akt | cyce | rb | e2f1 | p14arf | bcl2 | bax | caspases |
| 1 | Proliferation | 1 | Point | | | ■ | | | ■ | | | ■ | ■ | | ■ | | ■ | | |

**Figure S4 – Case Study 2: DA Results for p53 Network with DNA Damage OFF.** The DA results representing the activity state of each node in the absence of DNA damage. The black and white boxes depict the ON and OFF states of nodes, respectively.

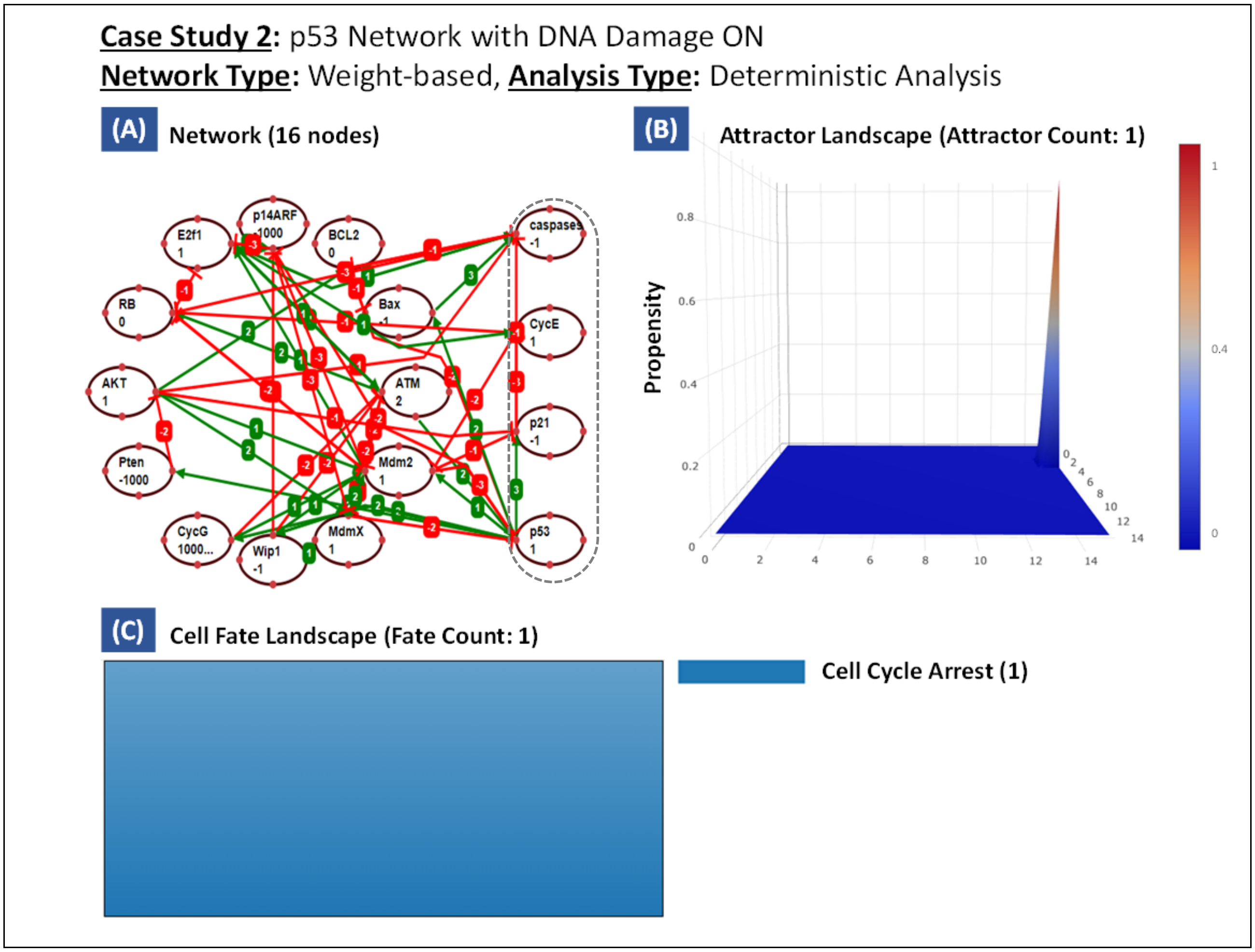

**Figure S5 – Case Study 2: p53 Network with DNA Damage ON.** (A) The network consisting of 16 nodes and 50 edges with 4 output nodes and 12 processing nodes constructed using TISON's NE, (B) Attractor Landscape with 1 cell fate attractor, and (C) Cell Fate Landscape with 1 cell fate representing the most probable attractor.

| Deterministic Results | | | | | | | | | | | | | | | | | | | |
|---|---|---|---|---|---|---|---|---|---|---|---|---|---|---|---|---|---|---|---|
| Attractor | Associated Cell Fate | Basin Size Ratio | Attractor Type | atm | p53 | mdm2 | mdmx | wip1 | cycg | pten | p21 | akt | cyce | rb | e2f1 | p14arf | bcl2 | bax | caspases |
| 1 | Cell Cycle Arrest | 1 | Cyclic | □ | □ | ■ | ■ | □ | ■ | □ | □ | ■ | □ | □ | ■ | □ | ■ | □ | □ |
| | | | | □ | □ | ■ | ■ | ■ | ■ | □ | ■ | ■ | □ | □ | ■ | □ | ■ | □ | □ |
| | | | | □ | ■ | ■ | ■ | ■ | ■ | □ | ■ | ■ | □ | ■ | ■ | □ | ■ | □ | □ |
| | | | | ■ | □ | □ | □ | □ | ■ | □ | □ | ■ | ■ | □ | ■ | □ | ■ | □ | □ |
| | | | | ■ | □ | ■ | □ | □ | ■ | □ | □ | ■ | ■ | □ | ■ | □ | ■ | □ | □ |
| | | | | ■ | ■ | □ | ■ | □ | ■ | □ | □ | ■ | ■ | ■ | ■ | □ | ■ | □ | □ |
| | | | | ■ | ■ | □ | ■ | ■ | ■ | □ | ■ | ■ | ■ | ■ | ■ | □ | ■ | □ | □ |

**Figure S6 – Case Study 2: DA Results for p53 Network with DNA Damage ON.** The DA results represent the activity state of each node in the presence of DNA damage. The black and white boxes reflect the ON and OFF states of nodes, respectively.

**Case Study 3 – Decoding Yeast Network for Cell Cycle Progression**

The third case study was on the yeast cell cycle network developed by Han *et al.* (5). The network contained 11 nodes and 34 edges (Figure S7A). NE's probabilistic analysis pipeline was employed to compute the basin size ratios followed by the construction of the cell fate landscape (see Supplementary Data – NE for further details).

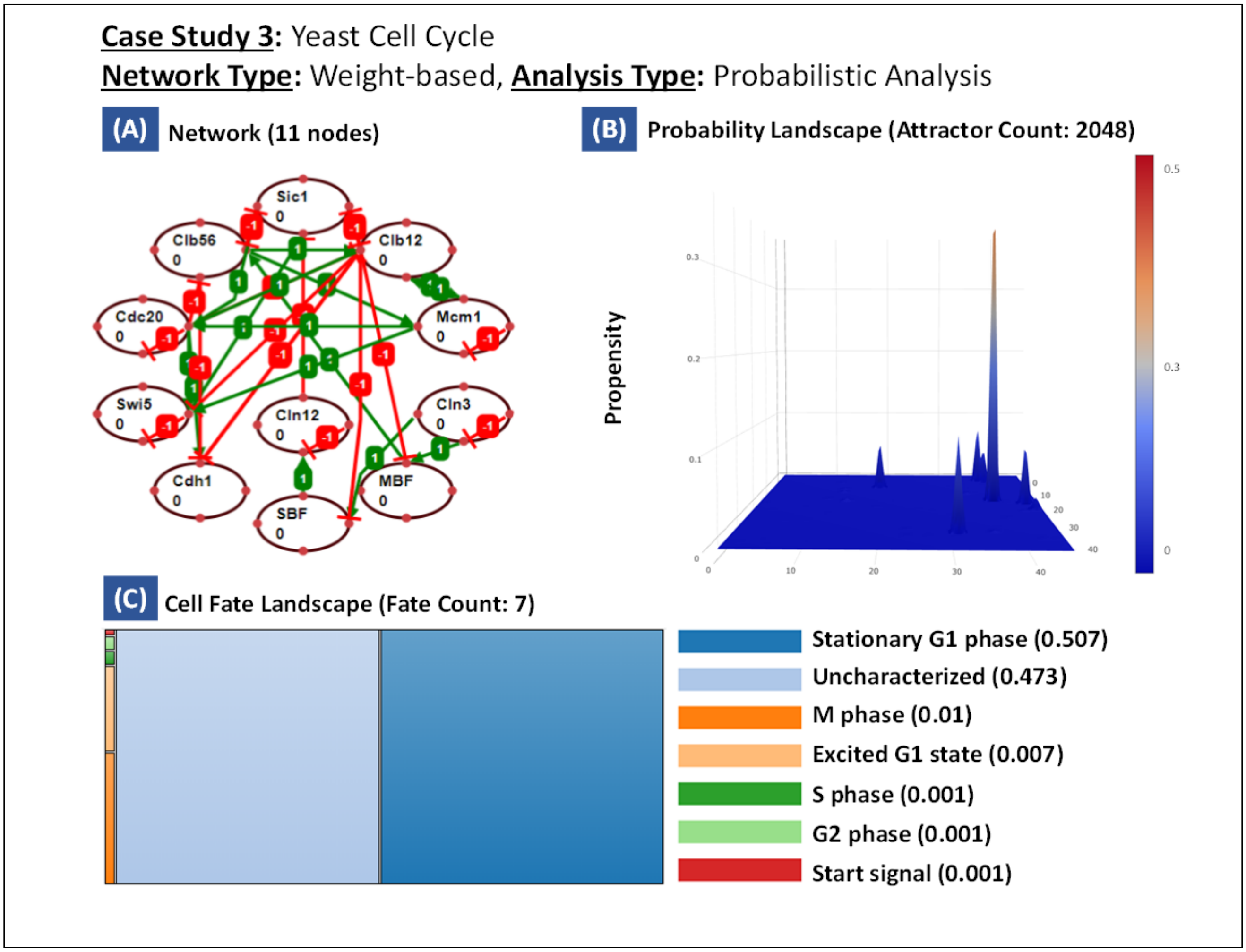

**Figure S7 – Case Study 3: Yeast Cell Cycle Progression Network.** (A) The network consisting of 11 nodes and 34 edges constructed using TISON's NE, (B) Probability Landscape with 2048 cell fate attractors, and (C) Cell Fate Landscape with 7 cell fate representing the most likely attractors.

- **Analysis Results**

To decode the dynamics of the yeast cell cycle progression network, we employed weight-based probabilistic analysis. 7 cell fate attractors (Figure S7B) were reported and the highest probability (0.507) was observed for the ‘stationary G1 phase’ state (Figure S7C). The cell cycle trajectory was computed to begin at ‘start signal’ and terminated at the ‘Stationary G1 phase’ (Figure S8). The basin size ratios reported by TISON were in agreement with the published study (see Supplementary D

| Probabilistic Results | | | | | | | | | | | | | | |
|---|---|---|---|---|---|---|---|---|---|---|---|---|---|---|
| Attractor | State Order in Trajectory | Associated Cell Fate | Probability | cln3 | mbf | sbf | cln12 | cdh1 | swi5 | cdc20 | clb56 | sic1 | clb12 | mcm1 |
| 69 | 13 | Stationary G1 phase | 0.539 | | | | | ■ | | | | ■ | | |
| 101 | 12 | Excited G1 state | 0.004 | | | | | ■ | ■ | | | ■ | | |
| 117 | 11 | M phase | 0.003 | | | | | ■ | ■ | ■ | | ■ | | |
| 52 | 10 | | 0.002 | | | | | | ■ | ■ | | | ■ | ■ |
| 56 | 9 | | 0.002 | | | | | | ■ | ■ | | ■ | ■ | ■ |
| 54 | 8 | | 0.002 | | | | | | ■ | ■ | | ■ | | ■ |
| 156 | 7 | | 0.001 | | | | ■ | | | ■ | ■ | | ■ | ■ |
| 905 | 6 | S phase | 0.001 | | ■ | ■ | ■ | | | | ■ | | | |
| 908 | 5 | G2 phase | 0.001 | | ■ | ■ | ■ | | | | ■ | | ■ | ■ |
| 897 | 4 | Excited G1 state | 0.001 | | ■ | ■ | ■ | | | | | | | |
| 965 | 3 | | 0.001 | | ■ | ■ | ■ | ■ | | | | ■ | | |
| 837 | 2 | | 0.001 | | ■ | ■ | | ■ | | | | ■ | | |
| 1093 | 1 | Start signal | 0.001 | ■ | | | | ■ | | | | ■ | | |

**Figure S8 – Case Study 3: PA Results File for Yeast Cell Cycle Progression Network.** The PA results representing the activity state of each node as the cell cycle progresses. Results from the trajectory represent the seven stages of a cell cycle (from 1 to 13 state transitions).

**Case Study 4 – Utilizing the Stem Cell Network to Investigate Cell Proliferation and Differentiation**

In the fourth case study, we validated NE’s ODE analysis feature by analyzing a human stem cell network published by Wang *et al.* (6). The network consisted of 52 nodes and 123 edges (Figure S9A). TISON was used to construct the ODE landscape using its ordinary differential equation analysis feature (see Supplementary Data – NE for further details).

- **Analysis Results**

To investigate the regulatory dynamics of a stem cell network, we employed ODE analysis to our reconstructed 52-node network. The ODE landscape exhibited the emergence of two attractors representing a stem cell and a differentiated cell (Figure S9B). The gene expression comparison between TISON and NetLand (7) was also performed and the results are shown in (Figure S10-11) (see Supplementary Data – NE for further details).

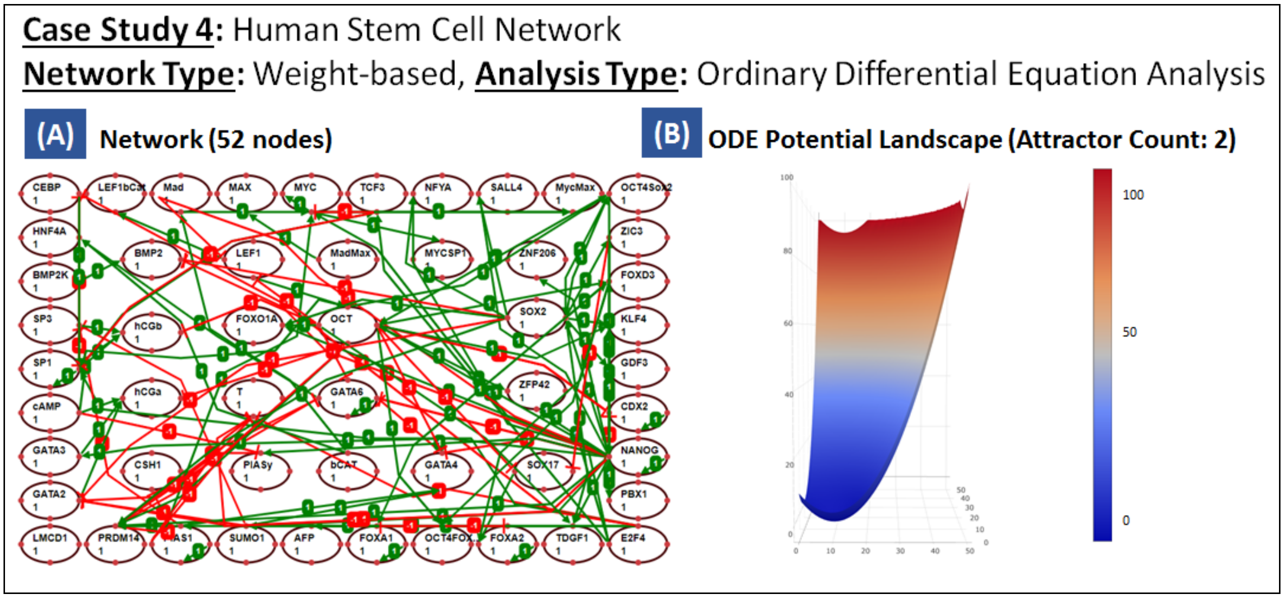

**Figure S9 – Case Study 4: Human Stem Cell Network.** (A) The network containing 52 nodes and 123 edges constructed using TISON's NE, and (B) ODE Potential Energy Landscape with 2 attractors.

- **Comparison with Published Literature**

To validate NE's ODE modeling feature, we compared it with "NetLand" (7). The results obtained provide individual gene expression values for both stem cell and differentiation attractors (Figure S10 and S11).

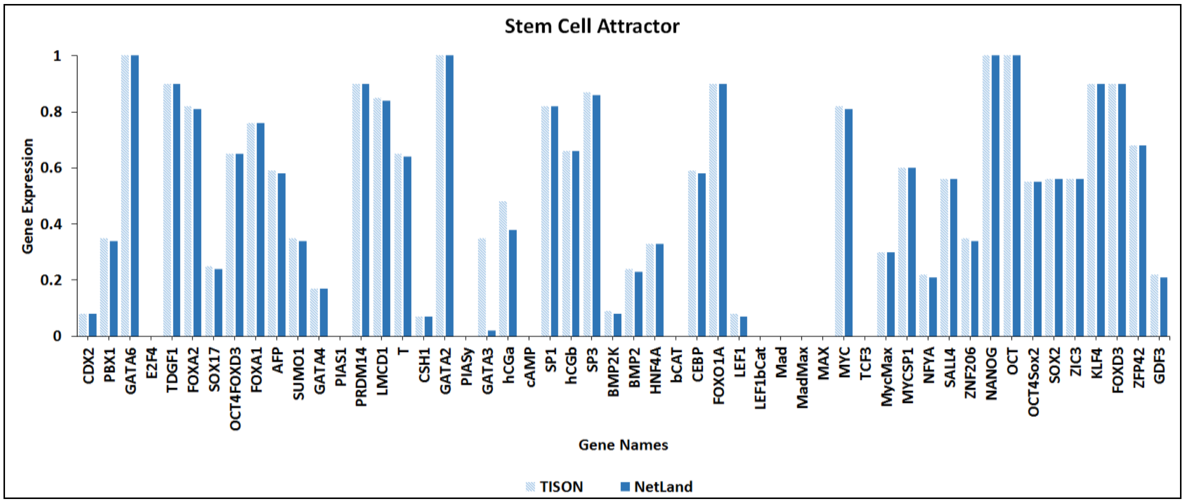

**Figure S10 – Case Study 4: Stem Cell Attractor.** The stem cell attractor representing 52 genes and their relative expression values. The gene expression values from NetLand were compared with the ones generated after ODE analysis in TISON.

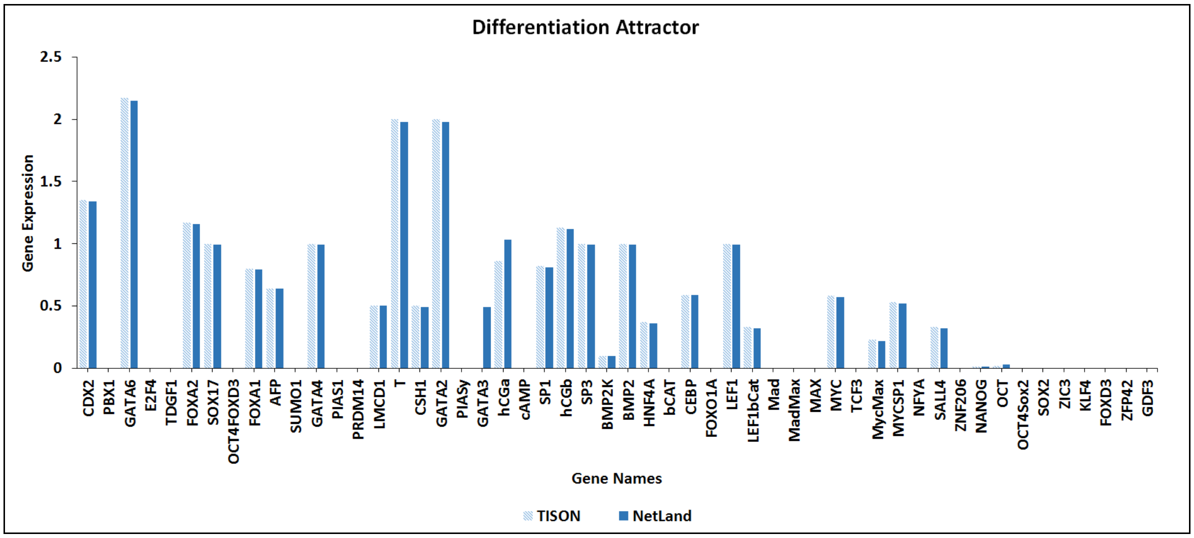

**Figure S11 – Case Study 4: Differentiation Attractor**. The differentiation attractor representing 52 genes and their relative expression values. The gene expression values from NetLand were compared with the ones generated after ODE analysis in TISON.

#### Section 2.1.3. User Manual & Video Tutorials for Networks Editor

**Table S1 - Reference Table for NE.** Description of supporting information along with the web links have been tabulated below for the TISON home page, user manual, issues table, datasets, and video tutorials on YouTube as a playlist.

| Item | Description | Link |
|---|---|---|
| **Project Home** | URL link for TISON project home | https://tison.lums.edu.pk/ |
| **User's Manual** | Editor's user manual can be found here. | https://tison.lums.edu.pk/Manuals/NetworksManual.pdf |
| **Issues** | Issues and related discussions can be added at the issues page. | https://github.com/BIRL/TISON/issues |
| **Datasets** | Sample Data files are available at the following link. | https://github.com/BIRL/TISON |
| **Playlist of Video Tutorials** | Video tutorials for step-by-step execution of case studies as well as employment of salient features | https://www.youtube.com/watch?v=oMv53VibuOg&list=PLaNVq-kFOn0YTO6RCa5mr2jwIkkD-uv9X&ab_channel=BiomedicalInformaticsResearchLab |

### Section 2.2. Therapeutics Editor – An Overview

#### Section 2.2.1. Software Features & Functionalities

To begin the process of therapeutic screening of networks using Therapeutics Editor (TE), TISON users first have to create a project (using Project Explorer) followed by designing of biomolecular networks (in Networks Editor, see Section 2.1 for details). Users can access TE from the Project Home Page or the editor bar at the bottom of the GUI. Networks developed in NE become automatically available in TE for the development and evaluation of therapies. TE can then assist its users in evaluating existing therapies from literature as well as eliciting novel drug targets (details in Section 2.2.2). Within each TE therapy, a single drug or multi-drug combination can be employed followed by their application on a biomolecular network.

- **Creating Therapies**

Users can create a therapy by selecting a pre-existing network and incorporating the effect of drugs by using the "*Therapeutics Steps*" panel. Each drug can act on a network by (i) node knock-up or knock-down, (ii) node knock-out, (iii) node knock-in, or (iv) altering edge interactions. Specifically, for rules-based networks, node knock-up or knock-down is implemented by assigning a new fixed user-defined value to the node along with the suppression of its upstream regulations (i.e. deleting the rule defining the regulation of a node). Knock-out for a rules-based network is implemented by deleting the node itself as well as its rule from the rest of the network. Knock-in, in contrast, is performed by adding the new node name alongside its defining rules in the network towards a seamless network rewiring. In the case of weight-based networks, node knock-up or knock-down is implemented by setting the incoming interaction weights and its basal values to zero (i.e. suppression of its upstream regulation) and assignment of a fixed (new) value for the target node. For node knock-out on a weight-based network is performed by setting the node value, as well as basal to zero and deleting all of its outgoing and incoming edges. Knock-in is implemented by incorporating node name, incoming and outgoing edges as well as setting up its basal value. This basal value can be either directly assigned by the user to calculated using expression values towards developing personalized cancer models. The alteration of edge interactions can only be implemented on weight-based networks, wherein the interaction weight value between two network nodes is replaced with a user-defined value. Additionally, edges can also be knocked out by setting their value to 0 or knocked in by defining its up and downstream nodes in the network.

These features have been incorporated into TE to facilitate the user in creating single-drug therapy as well as multi-drug therapy which may contain multiple drugs. Importantly, the therapy construction process can be undertaken in the form of a "*Horizontal therapy*" or a "*Vertical therapy*". In horizontal therapy, each drug can be programmed to target multiple nodes or node interactions, concurrently. In this case, multiple network nodes' values or interactions can be altered by a single drug, and the impact of the drug on the target node(s) is implemented independently, within an analysis. In a vertical therapy, multiple drugs with distinct node targets can be collated to act in tandem. This involves a sequential implementation of the effect of each drug. The node propensities obtained from one drug screening step are used as inputs for the next drug screening step using the heuristic DA pipeline. The resulting output contains the cumulative effect of each therapeutic step performed on the network. Furthermore, therapies can be exported from or imported into the TE. Currently, the only integrated drug database is *The Drug Gene*

*Interaction Database* (DGIdb) to provide drug score to users, for ease. Users can select a target node along with a drug of choice and its node expression value is integrated directly into the therapy model, from the database. Additionally, users can perform *Exhaustive Screening* (ES) on the network which allows them to exhaustively evaluate each or any set of nodes in the network towards investigating the effect of each node in the network. This is especially useful in the case of developing personalized therapeutics, wherein, users can evaluate the most efficacious set of nodes using the ES feature in TE. To validate the functioning of the editor as well as to facilitate user employment of TE, two template case studies have been provided including a rules-based and a weight-based therapy. These case studies can be accessed through the "Upload Case Study" button in TE or through the template project tab on the TISON's home page.

- **Analyzing Therapies**

Once therapy has been created in TE, three types of network analyses are available depending upon the type of therapy. For therapeutic screening of weight-based networks, deterministic analysis, probabilistic analysis, and ordinary differential equation analysis can be performed. For rules-based networks, only deterministic analysis can be performed (see Section 2.2.3 for TE's User Manual).

- **Visualizing Therapies**

Users can view as well as download results directly from the therapeutics editor. Result visualization is available in the form of cell fate landscape, attractor landscape, potential energy landscape, probability landscape, and ODE landscape depending on the type of analysis done. TE also offers a comparative analysis feature, which allows the user to compare the results of therapeutic evaluation between drugs and the control case. These results can be visualized in the form of bar charts or stack charts and are downloadable (see Section 2.2.3 for TE's User Manual).

#### Section 2.2.2. Functional Validation of the Therapeutics Editor

We have undertaken functional validation of TE by reconstructing two published case studies which include (i) therapeutic strategies for p53-mediated apoptosis network under DNA damage ON and OFF (4), and (ii) induction of colorectal tumorigenesis through mutation implementation in the human signaling network (3). These case studies show that TE can be employed for adding therapy as well as introducing mutations. Results for deterministic and probabilistic analyses, attractor landscapes, and basin ratios were compared with the published literature for both case studies. The sub-sections below elaborate on the results obtained from the analyses.

**Case Study 1 – Therapeutic strategies for p53-mediated Apoptosis Network in MCF7 Breast Cancer Cell Line**

The first case study for validation of TE is built upon Choi *et al.'s* weight-based p53 network to identify the interactions which determine the cellular response to DNA damage. The reconstructed network contains 16 nodes and 50 edges. TISON was used to perform DA on the p53 network with both DNA damage ON and OFF, by incorporating various mutations and corresponding therapies. The results obtained were compared with Choi *et al.* and found to be in agreement (see Supplementary Data – TE for further details).

- **Analysis Results (DNA Damage OFF)**

To investigate the cell fate programming dynamics of the mutated p53 network (with DNA damage OFF), DA was performed. Under control conditions, the analysis generated only one attractor, corresponding to proliferation (Figure S12). Similarly, after inhibiting the Wip1 node in the network, we obtained the attractor for proliferation (Figure S13). However, in the case of Nutlin, three attractors emerged: two correspondings to the senescence cell fate with a basin size ratio of 0.777 and one attractor corresponding to apoptosis with a basin size ratio of 0.223 (Figure S14). When the network was treated with both Wip1 and Nutlin inhibitors, two attractors remained; one for apoptosis with a basin size ratio of 0.578 and the other for senescence with a basin size ratio of 0.422 (Figure S15) (see Supplementary Data – TE for further details).

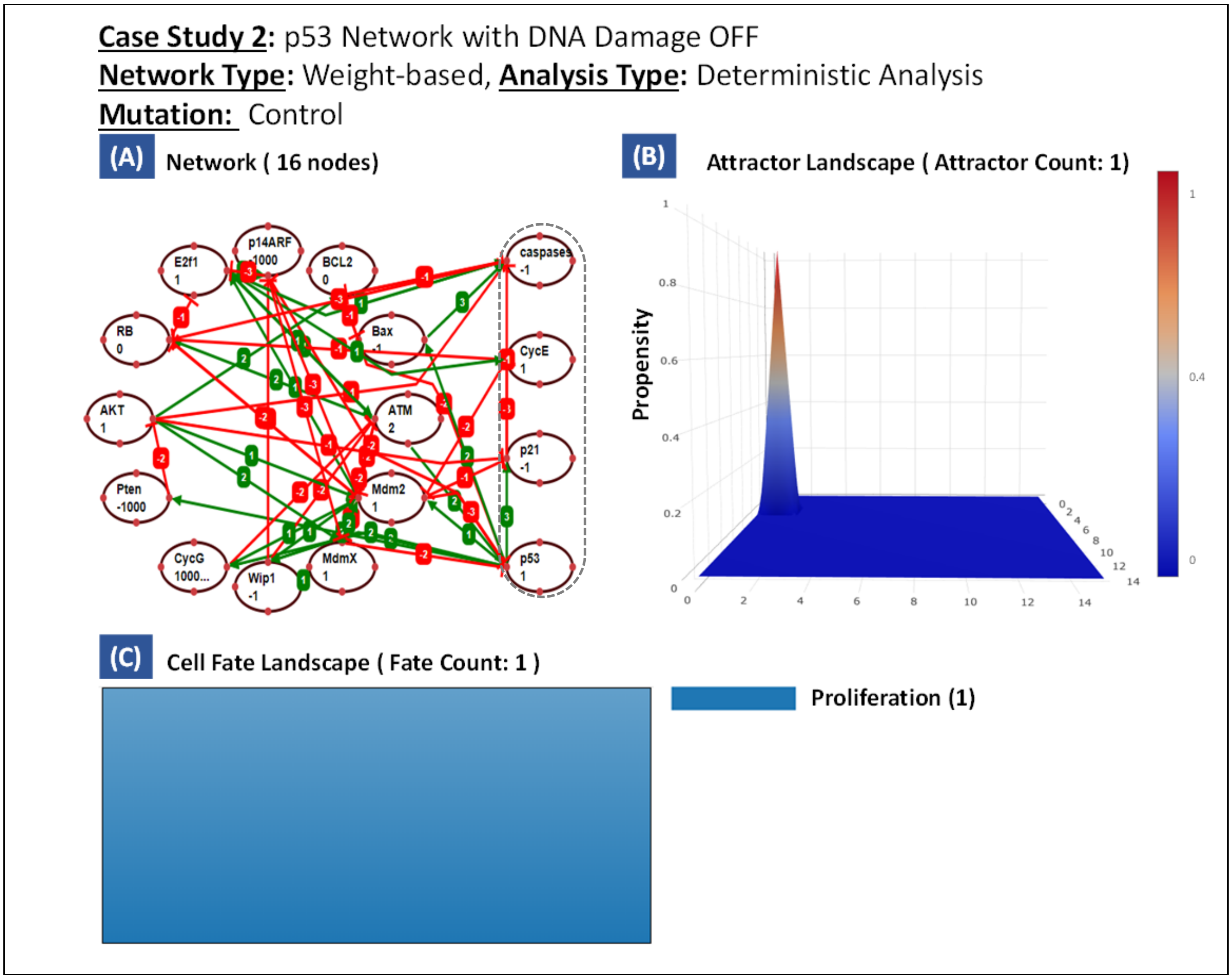

**Figure S12 – Case Study 1: p53-Mediated Apoptosis Network under Control Conditions.** (A) Network constructed using TISON's NE containing 12 processing nodes, 4 output nodes, and 50 edges, (B) Attractor Landscape with 1 cell fate attractor, and (C) Cell Fate Landscape representing 1 cell fate.

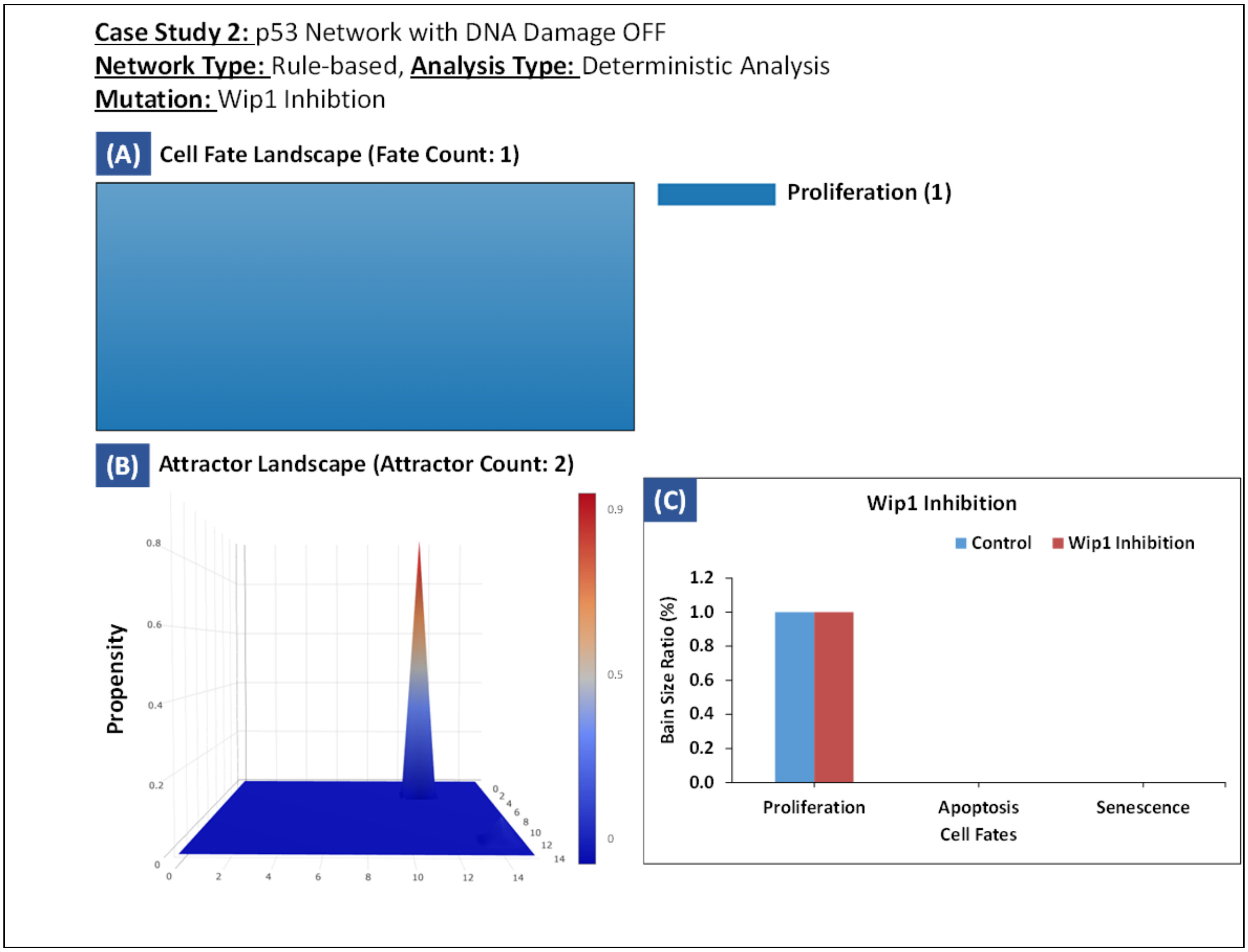

**Figure S13 – Case Study 1: p53-Mediated Apoptosis Network (DNA Damage OFF) with WIP1 Inhibition.** (A) Cell Fate Landscape with 1 cell fate representing the most likely attractor, (B) Attractor Landscape with 2 cell fate attractors, and (C) Comparison between control and Wip1 inhibition therapy.

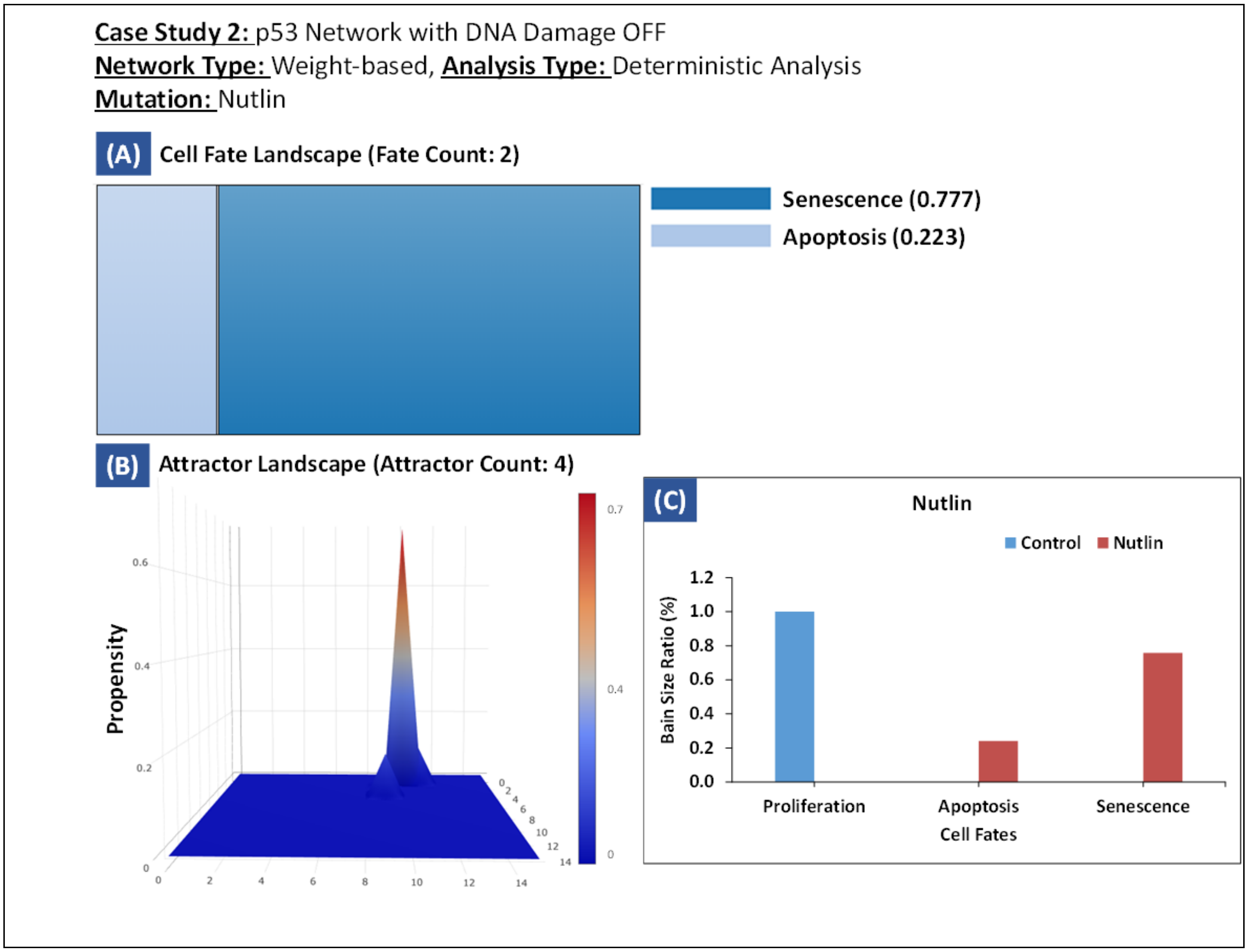

**Figure S14 – Case Study 1: p53-Mediated Apoptosis Network (DNA damage OFF) with Nutlin.** (A) Cell Fate Landscape with 2 cell fates representing the most likely attractors, (B) Attractor Landscape with 4 cell fate attractors, and (C) Comparison between control and Nutlin therapy.

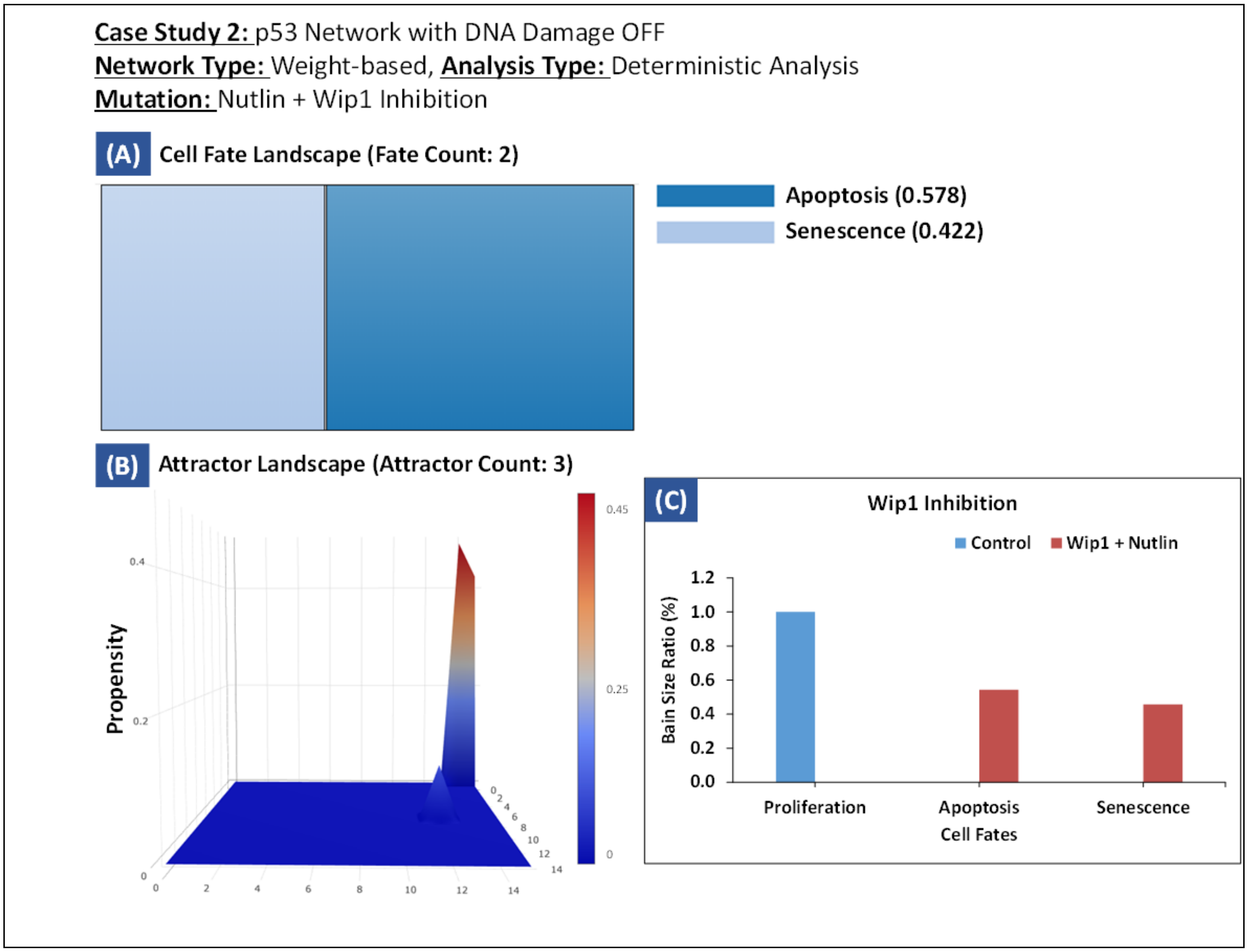

**Figure S15 – Case Study 1: p53-Mediated Apoptosis Network (DNA Damage OFF) with Nutlin and Wip inhibition.** (A) Cell Fate Landscape with 2 cell fates representing the most likely attractors, (B) Attractor Landscape with 3 cell fate attractors, and (C) Comparison between control and Nutlin + Wip Inhibition therapy.

| Conditions | Cell Fate | Basin Size Ratio (%) | Attractor Type | atm | p53 | mdm2 | mdmx | wip1 | cycg | pten | p21 | akt | cyce | rb | e2f1 | p14arf | bcl2 | bax | caspases |
|---|---|---|---|---|---|---|---|---|---|---|---|---|---|---|---|---|---|---|---|
| Control | Proliferation | 1 | Point | | | ■ | | | ■ | | | ■ | ■ | | ■ | | ■ | | |
| Wip1 Inhibition | Proliferation | 0.94531 | Point | | | ■ | | | ■ | | | ■ | ■ | | ■ | | ■ | | |
| Nutlin | Senescence | 0.78906 | Point | | ■ | ■ | | ■ | ■ | | | ■ | ■ | | ■ | | ■ | | |
| | Apoptosis | 0.10547 | | | ■ | ■ | | ■ | ■ | | ■ | ■ | | | ■ | | | ■ | ■ |
| | | 0.10547 | | | ■ | ■ | | ■ | ■ | | | ■ | ■ | | ■ | | | ■ | ■ |
| Wip1 + Nutlin | Senescence | 0.45703 | Point | | ■ | ■ | | | ■ | | | ■ | ■ | | ■ | | ■ | | |
| | Apoptosis | 0.44141 | | | ■ | ■ | | | ■ | | | ■ | ■ | | ■ | | | ■ | ■ |
| | | 0.09375 | | | ■ | ■ | | | ■ | | ■ | ■ | | | ■ | | | ■ | ■ |
| | Senescence | 0.00391 | | | ■ | ■ | | | ■ | | ■ | ■ | | | ■ | | ■ | ■ | |
| | | 0.00391 | | | ■ | ■ | | | ■ | | ■ | ■ | | | ■ | | ■ | | |

**Figure S16 – Case Study 1: Drug Screening Results from p53-Mediated Apoptosis Network (DNA Damage OFF).** The cell fates, basin size ratios, and attractor types of each node were obtained from control and under three therapies: Wip1 inhibition, Nutlin, and Wip1 inhibition + Nutlin.

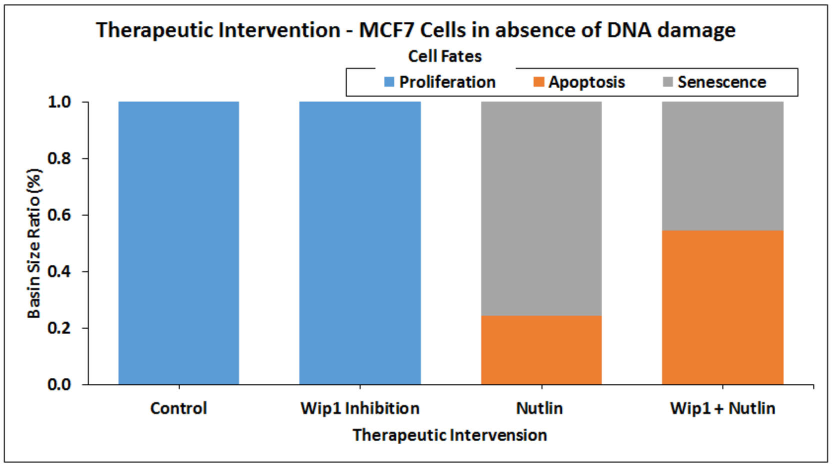

**Figure S17 – Case Study 1: p53-Mediated Apoptosis Network Analysis.** Representing cell fates in the system with DNA damage OFF in the case of Wip1 inhibition, Nutlin, and Wip1 inhibition + Nutlin.

- **Analysis Results (DNA Damage ON)**

To investigate the dynamics of the weight-based p53 network (with DNA damage ON) under mutations, DA was performed. Under control conditions, the analysis generated only one cell cycle arrest attractor (Figure S18). Similarly, after inhibiting the Wip1 node, two attractors emerged: one representing cell cycle arrest with a basin size ratio of 0.941 and the other representing apoptosis with a basin size ratio of 0.059 (Figure S19). However, after inhibiting Nutlin, the network generated three attractors corresponding to senescence, cell cycle arrest, and apoptosis with a basin size ratio of 0.148, 0.57, and 0.281, respectively (Figure S20). Upon a concurrent inhibition of Wip1 and Nutlin, two attractors representing apoptosis with a basin size ratio of 0.723 and senescence with a basin size ratio of 0.277 were obtained (Figure S21) (see Supplementary Data – TE for further details).

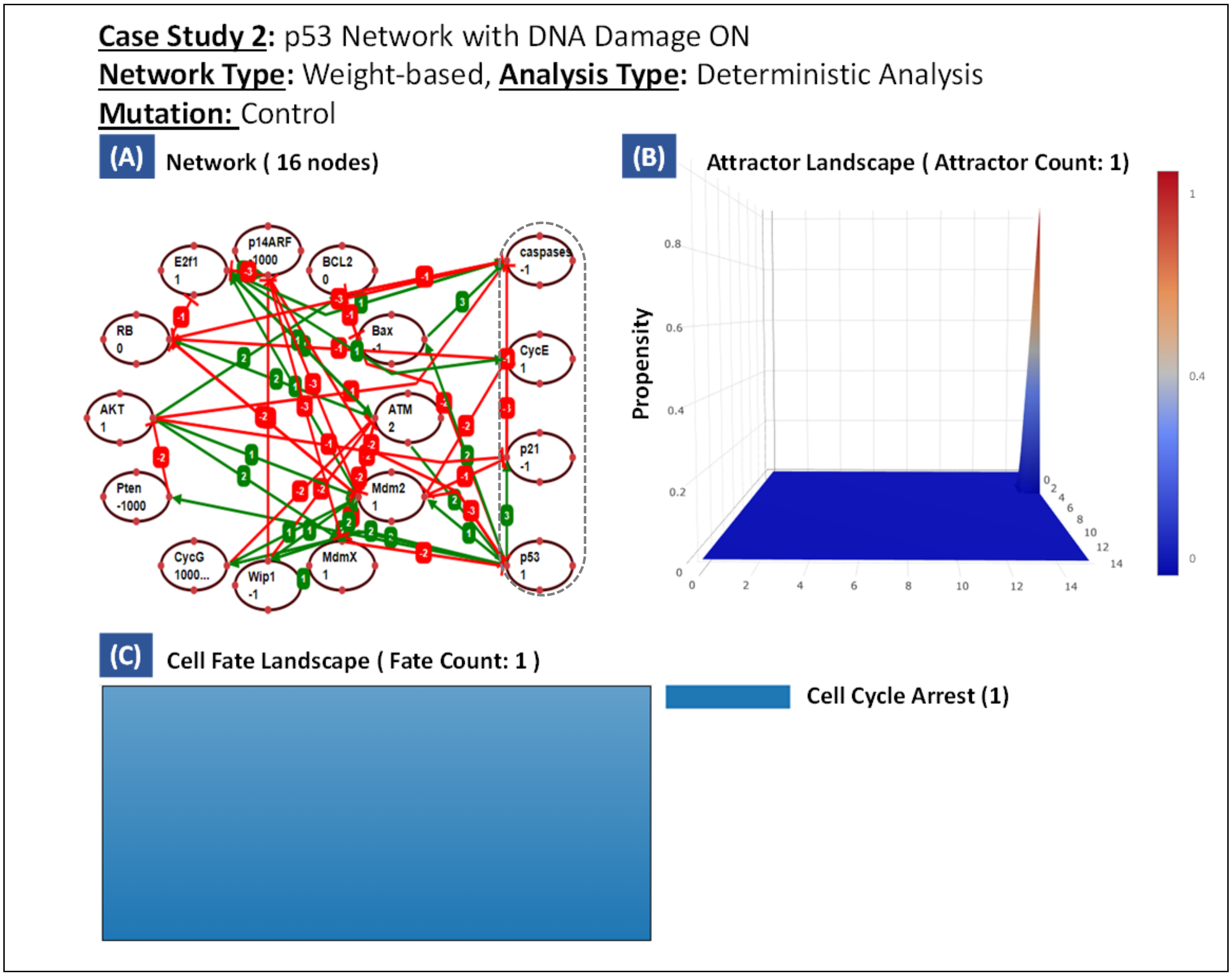

**Figure S18 – Case Study 1: p53-Mediated Apoptosis Network with DNA Damage ON under Control Conditions.** (A) Network constructed using TISON's NE containing 12 processing nodes, 4 output nodes, and 50 edges, (B) Attractor Landscape with 1 cell fate attractor, and (C) Cell Fate Landscape representing 1 cell fate.

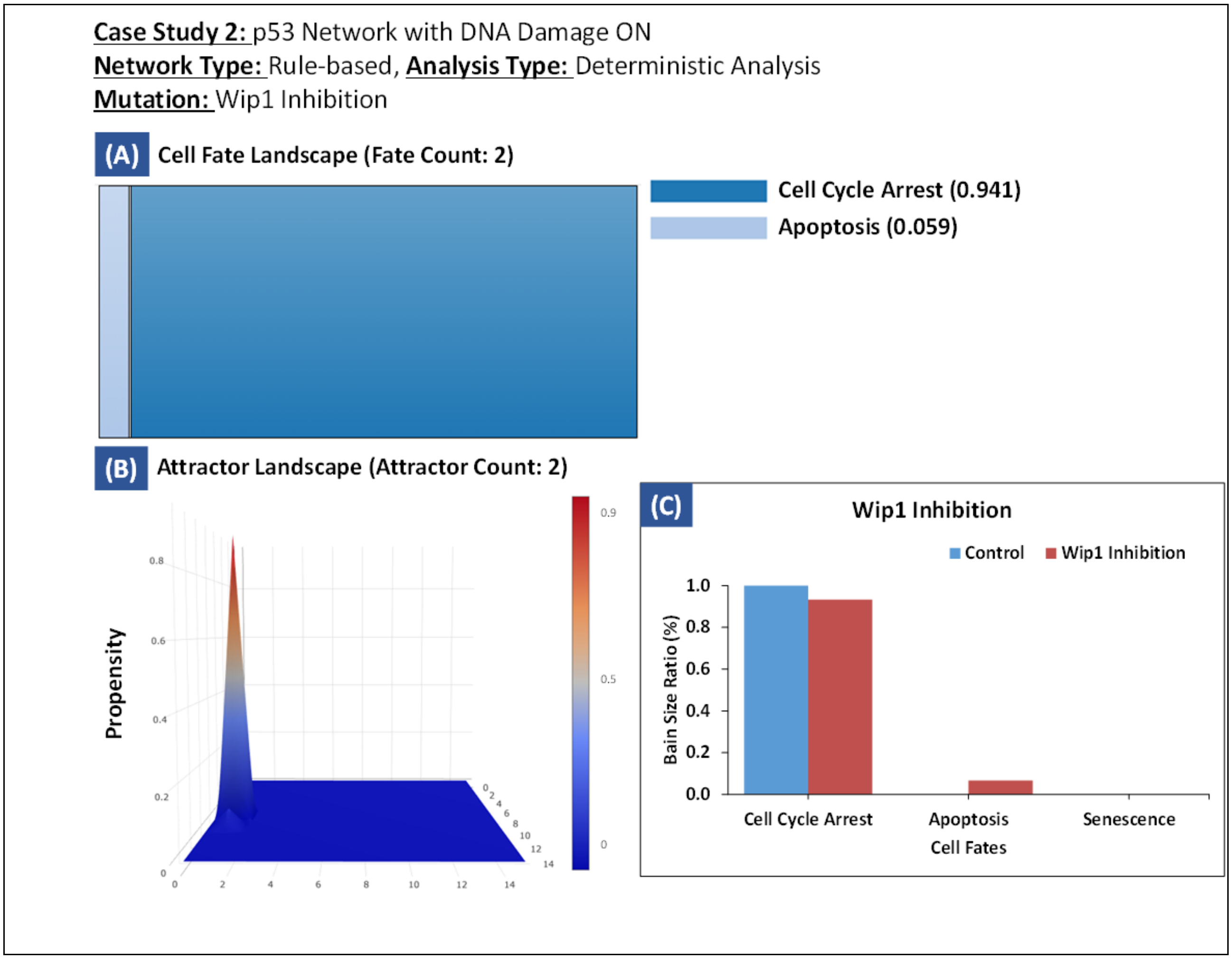

**Figure S19 – Case Study 1: p53-Mediated Apoptosis Network (DNA Damage ON) with WIP1 Inhibition.** (A) Cell Fate Landscape with 2 cell fates representing the most likely attractors, (B) Attractor Landscape with 2 cell fate attractors, and (C) Comparison between control and Wip1 inhibition therapy.

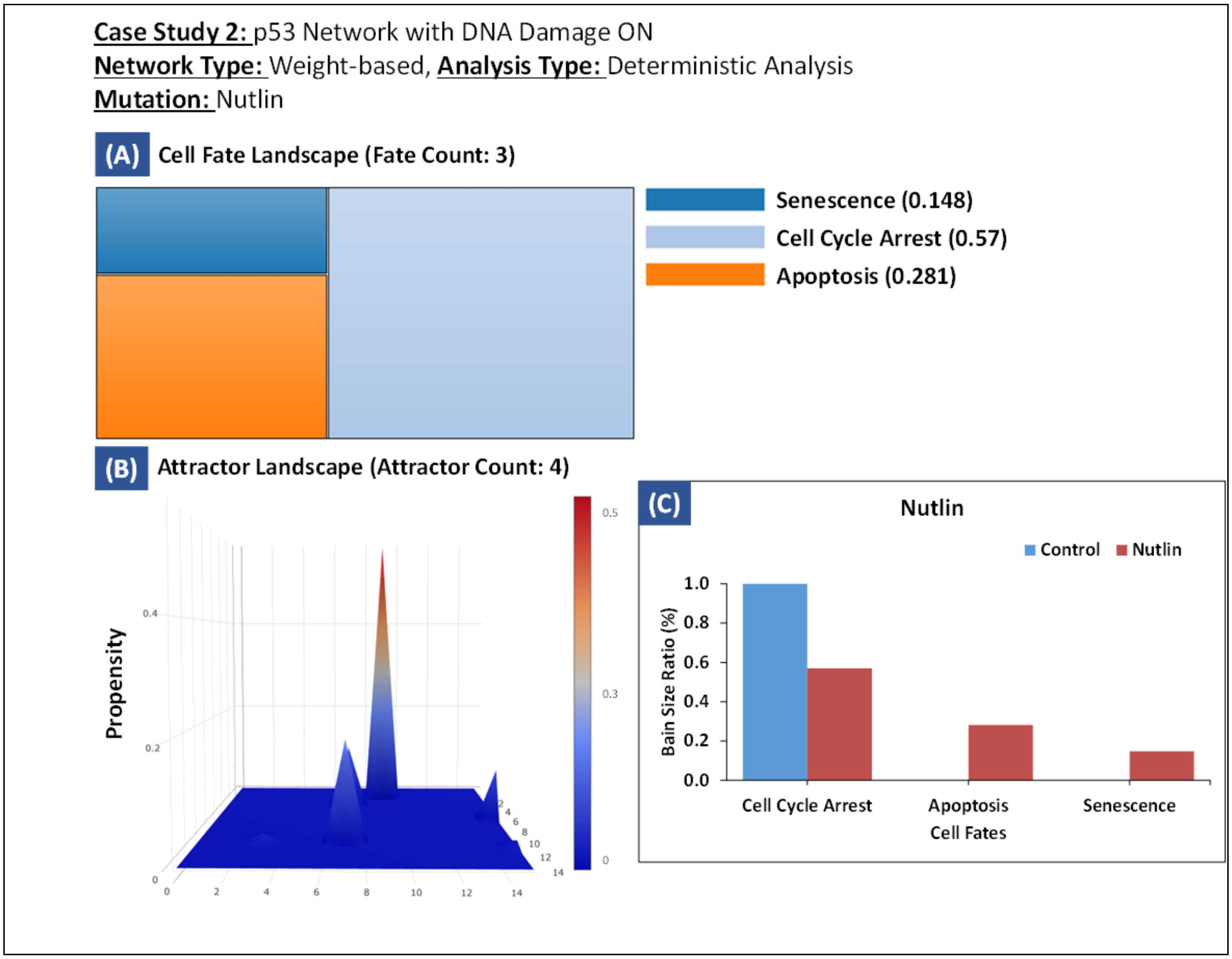

**Figure S20 – Case Study 1: p53-Mediated Apoptosis Network (DNA Damage ON) with Nutlin**. (A) Cell Fate Landscape with 3 cell fates representing the most likely attractors, (B) Attractor Landscape with 4 cell fate attractors, and (C) Comparison between control and Nutlin therapy.

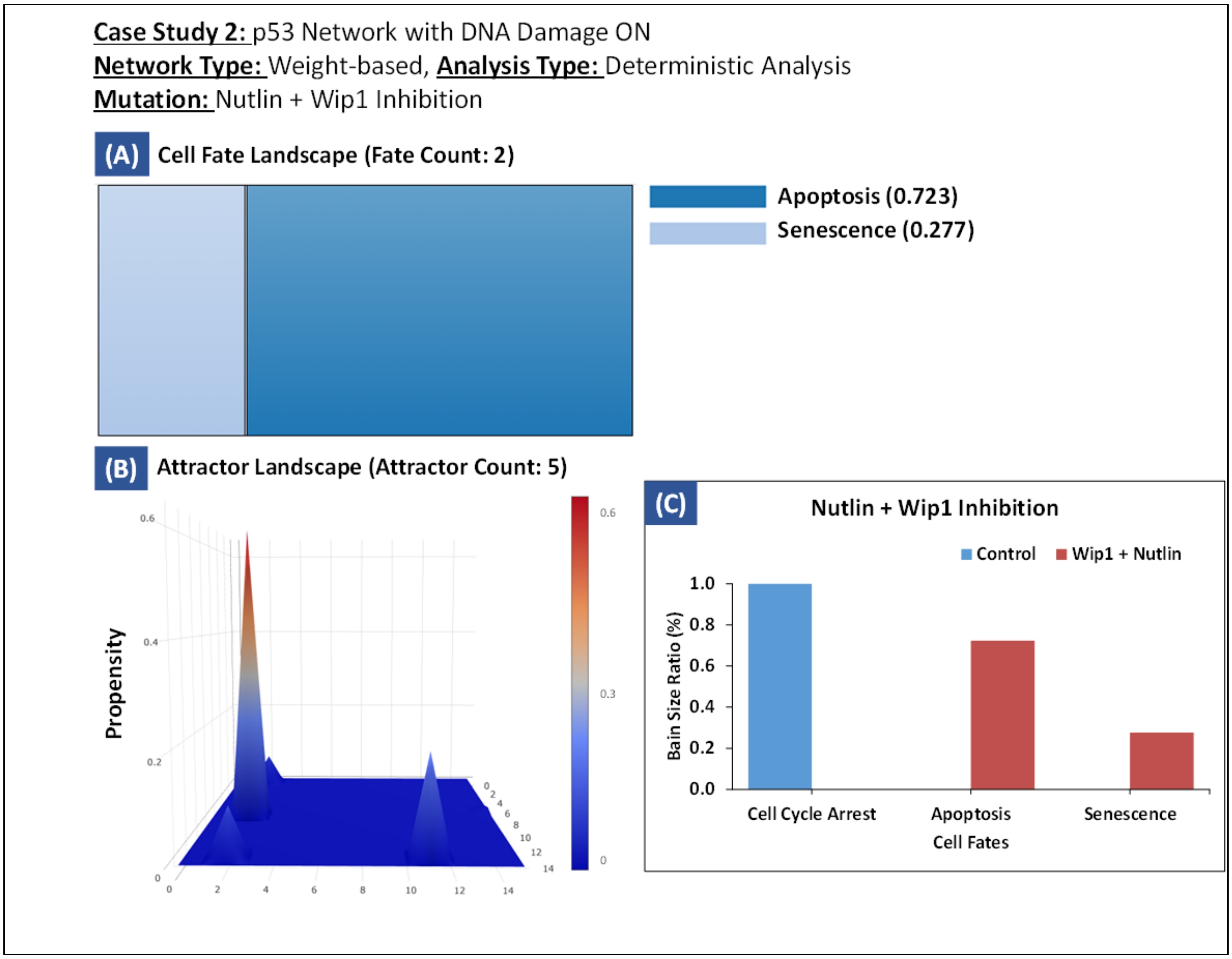

**Figure S21 – Case Study 1: p53-Mediated Apoptosis Network (DNA Damage ON) with Nutlin and Wip1 Inhibition.** (A) Cell Fate Landscape with 2 cell fates representing the most likely attractors, (B) Attractor Landscape with 5 cell fate attractors, and (C) Comparison between control and Nutlin + Wip Inhibition therapy.

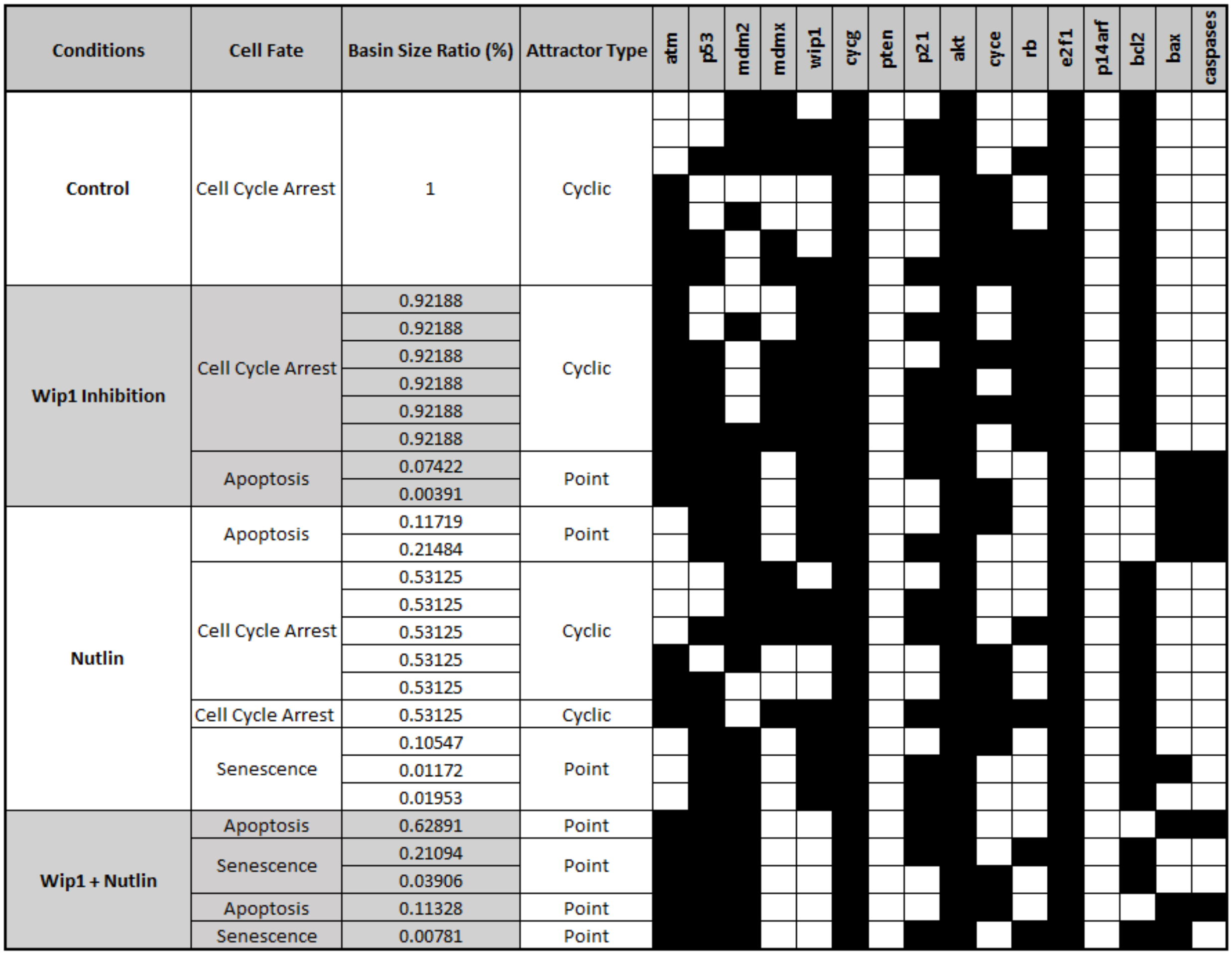

**Figure S22 – Case Study 1: p53-Mediated Apoptosis (DNA Damage ON) Network Analysis Summary.** The figure shows states of each node in the system with DNA damage ON in case of control, Wip1 inhibition, Nutlin, and Wip1 inhibition + Nutlin.

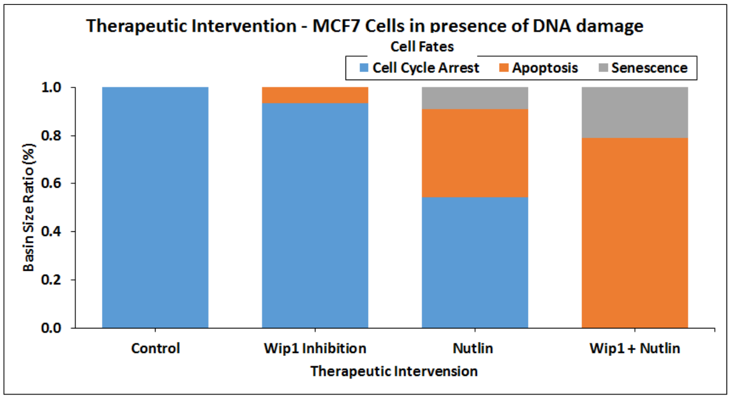

**Figure S23 – Case Study 1: p53-Mediated Apoptosis (DNA Damage ON) Network Analysis**. The figure represents cell fates in the system with DNA damage ON in case of control, Wip1 inhibition, Nutlin, and Wip1 inhibition + Nutlin.

**Case Study 2 – Investigating Colorectal Tumorigenesis by Induction of Sequential Mutations in Human Signaling Network**

In the second case study, we reconstructed a rules-based human signaling network reported by Cho *et al.* for observing colorectal tumorigenesis under sequential mutations (3). The network consisted of 197 nodes and 744 edges (Figure S24A). TE was employed to design a therapy by incorporating sequential mutations into the constructed network which resulted in colorectal tumorigenesis. Both the control and therapy cases were analyzed using DA followed by a comparison of resultant cell fates with those reported by Cho *et al.* (see Supplementary Data – TE for further details).

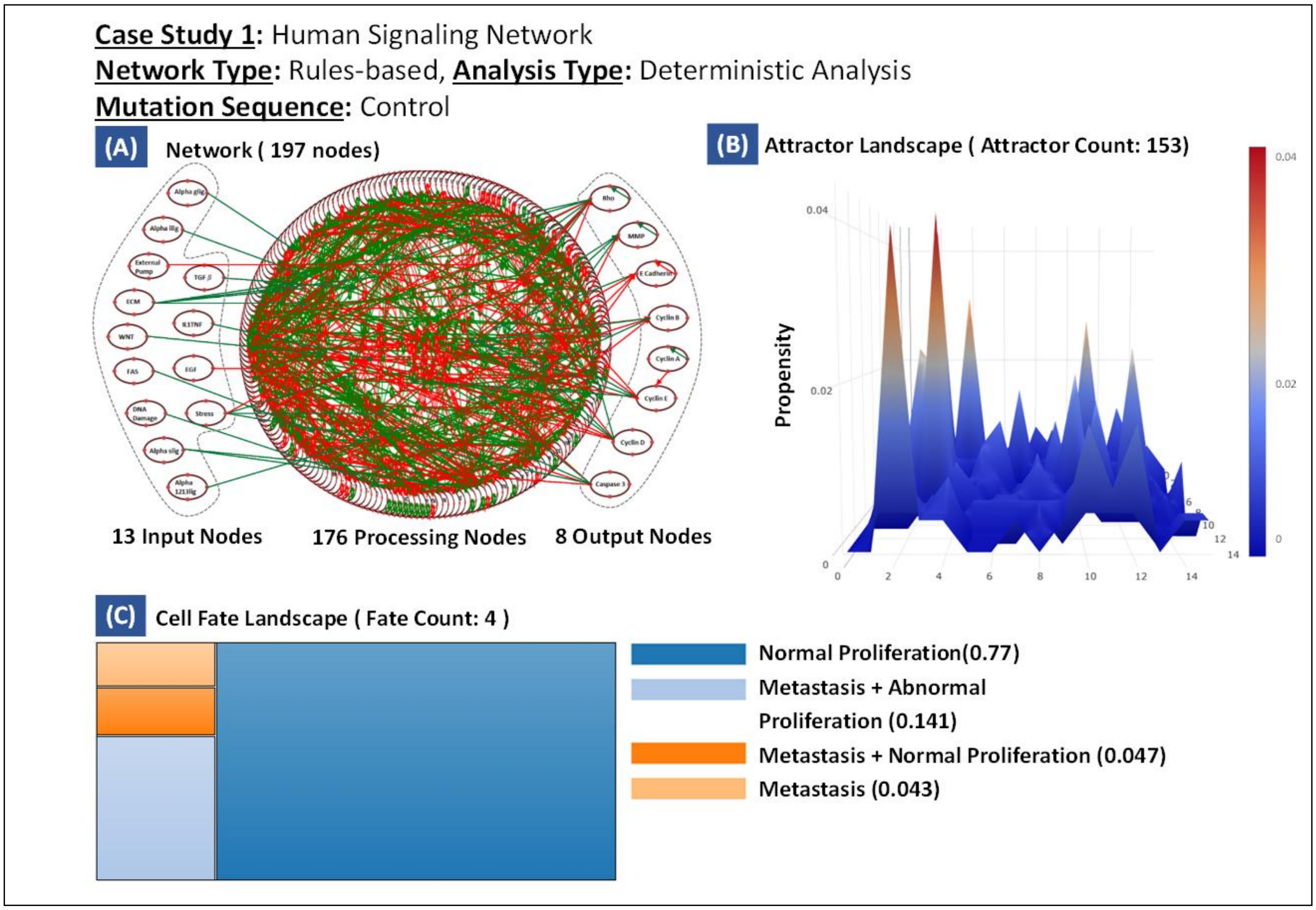

**Figure S24 – Case Study 2: Human Signaling Network Analysis in Control Conditions.** (A) Network constructed using TISON's NE consisting of 197 nodes and 744 edges with 13 input nodes, 8 output nodes, and 176 processing nodes, (B) Attractor Landscape with 153 cell fate attractors, and (C) Cell Fate Landscape representing the most likely attractor states.

- **Analysis Results**

To model the induction of colorectal tumorigenesis under sequential mutations, we used the rules-based DA pipeline in TE (Figure S24A). The resulting landscape for control therapy exhibited four major attractors in response to normal input conditions (Figure S24B) and their corresponding cell fates are shown in Figure S24C. Next, we incorporated sequential mutations which included (1) APC mutation (Figure S25), (2) APC and RAS mutations (Figure S26), (3) APC, RAS and PTEN mutations (Figure S27), and (4) APC, RAS, PTEN and P53 mutations (Figure S28). The results indicated the emergence of cancerous cell fates such as abnormal proliferation and metastasis, as a result of mutations.

A cell fate comparison between the control and mutations induced therapies was also undertaken (Figure S29). The basin size ratio comparison within each cell fate between TISON and Cho *et al.* is provided in Figure S30. Additionally, Figure S31 contains the mutation-specific basin size ratio comparison reported by TISON and Cho *et al.* for each of the cell fates. In conclusion, the results obtained from TISON tallied well with Cho *et al.'s* case study (see Supplementary Data – TE for further details).

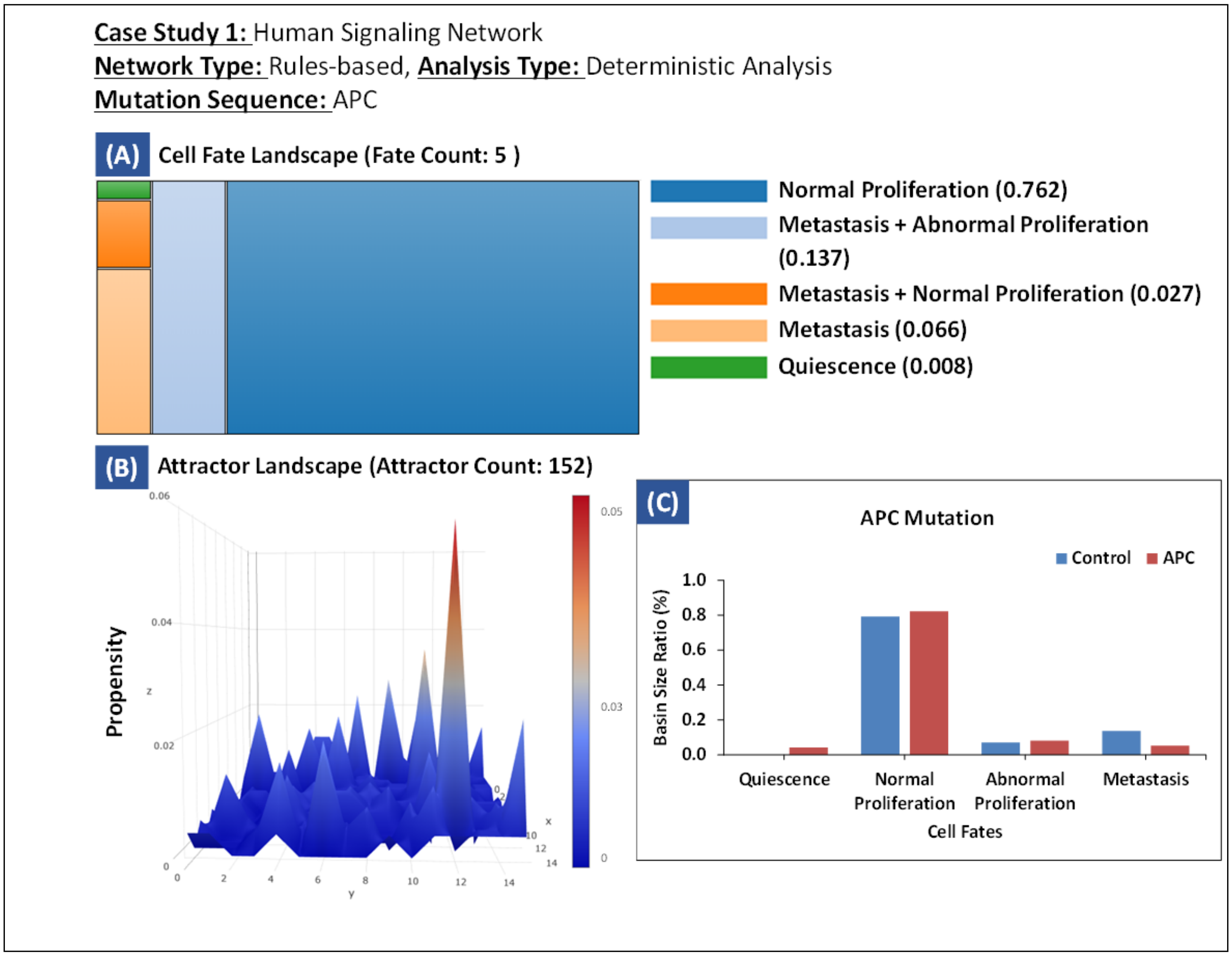

**Figure S25 – Case Study 2: Human Signaling Network Analysis with the Incorporation of APC Mutation** (A) Cell Fate Landscape representing five most likely attractors, (B) Attractor Landscape with 152 cell fate attractors, and (C) Comparison of basin size ratios of cell fates between the control and APC mutation model.

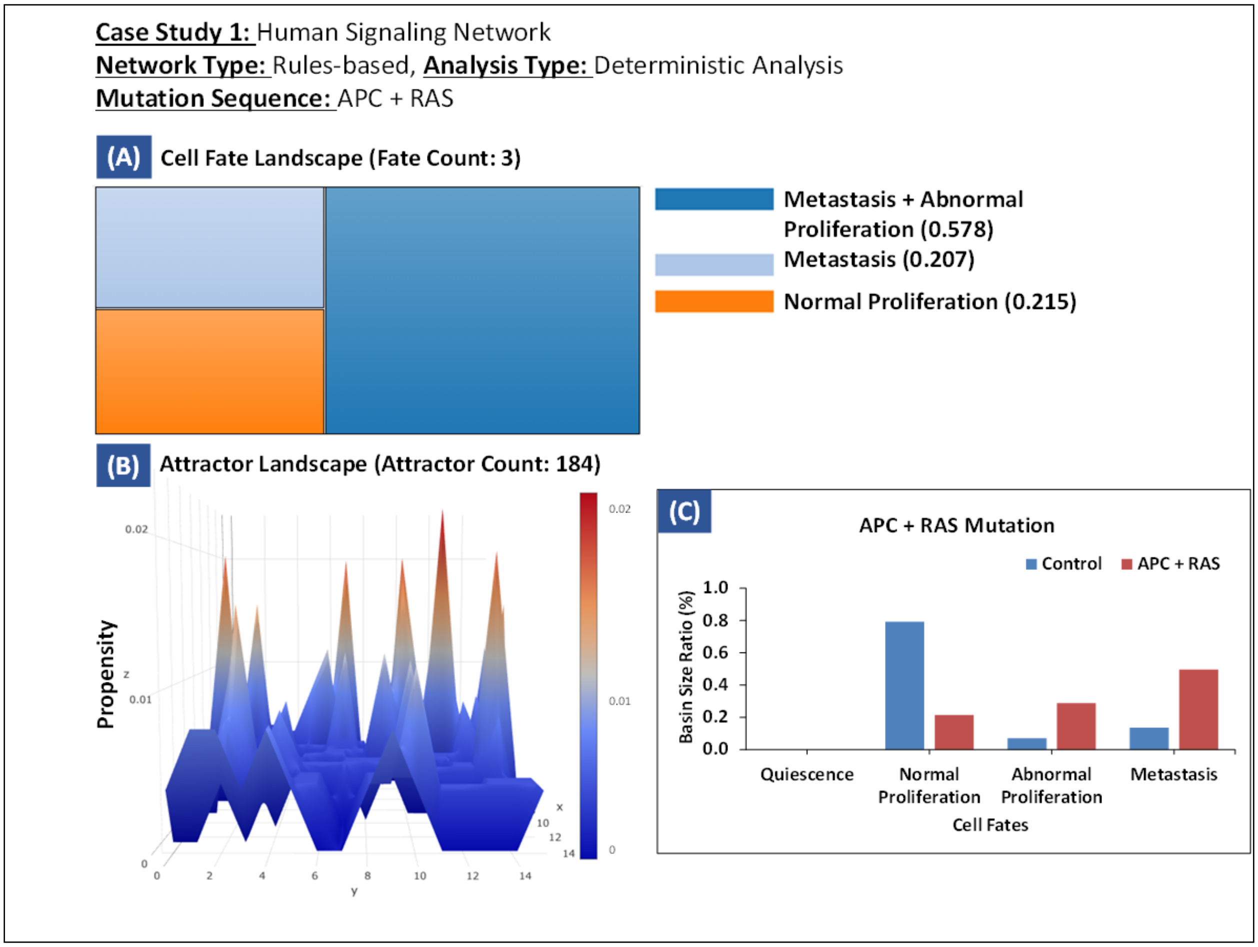

**Figure S26 – Case Study 2: Human Signaling Network Analysis with the Incorporation of APC and RAS Mutations**. (A) Cell Fate Landscape representing three most likely attractors, (B) Attractor Landscape with 184 cell fate attractors, and (C) Comparison of basin size ratios of cell fates between the control and APC+RAS mutation model.

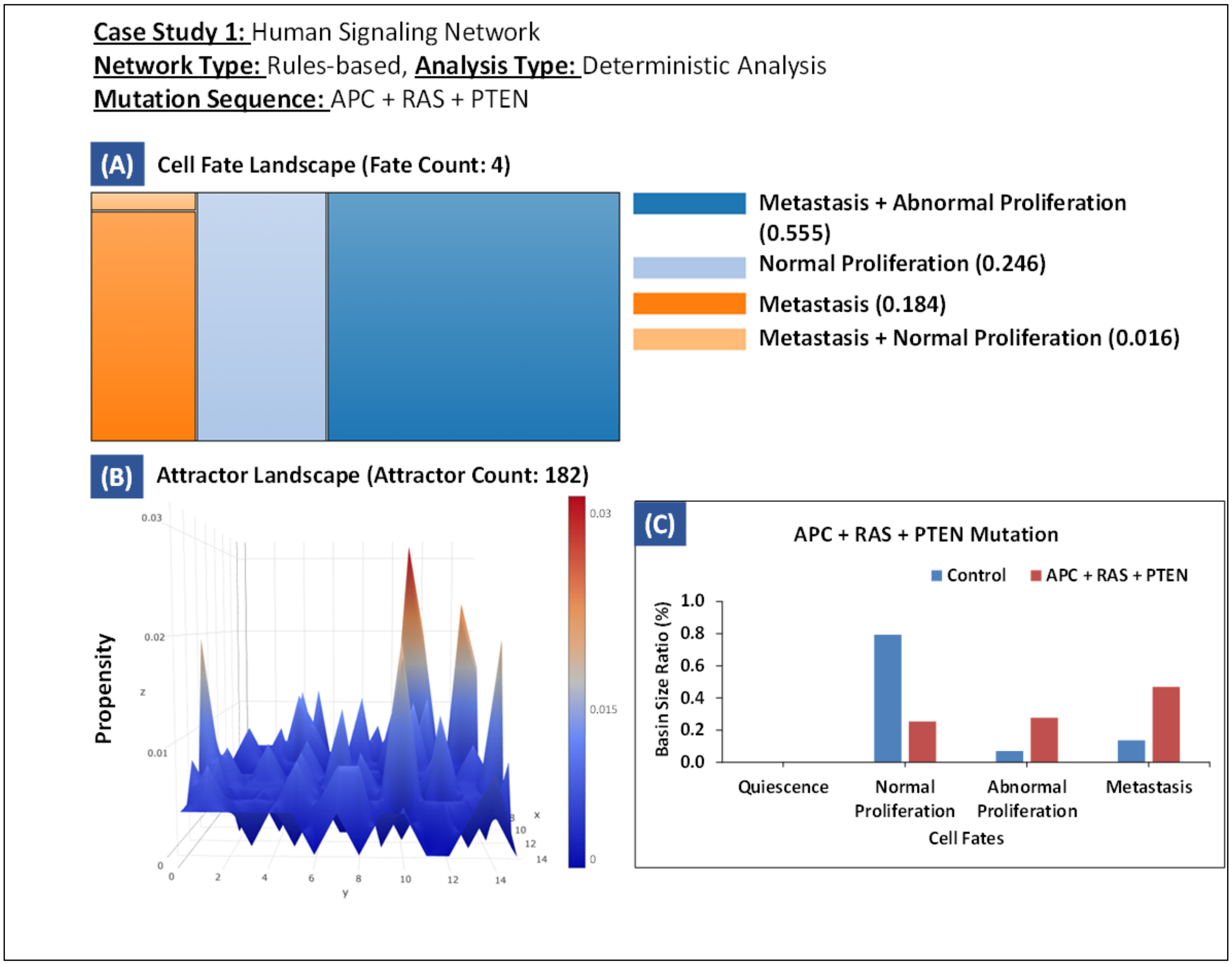

**Figure S27 – Case Study 2: Human Signaling Network Analysis with the Incorporation of APC, RAS, and PTEN Mutations.** (A) Cell Fate Landscape representing four most likely attractors, (B) Attractor Landscape with 182 cell fate attractors, and (C) Comparison of basin size ratios of cell fates between the control and APC+RAS+PTEN mutation model.

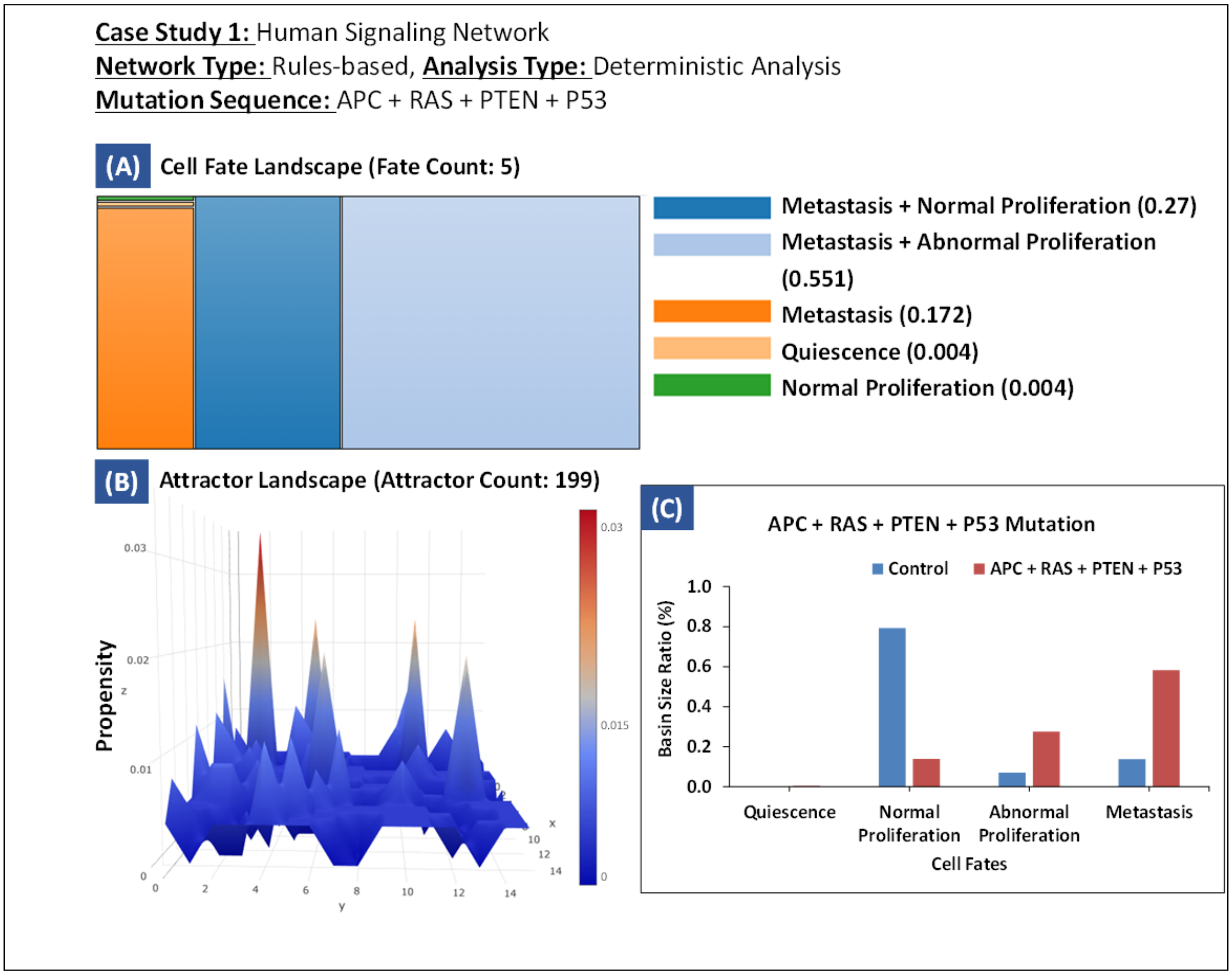

**Figure S28 – Case Study 2: Human Signaling Network Analysis with the Incorporation of APC, RAS, PTEN, and P53 Mutations**. (A) Cell Fate Landscape representing five most likely attractors, (B) Attractor Landscape with 199 cell fate attractors, and (C) Comparison of basin size ratios of cell fates between the control and APC+RAS+ PTEN+P53 mutation model.

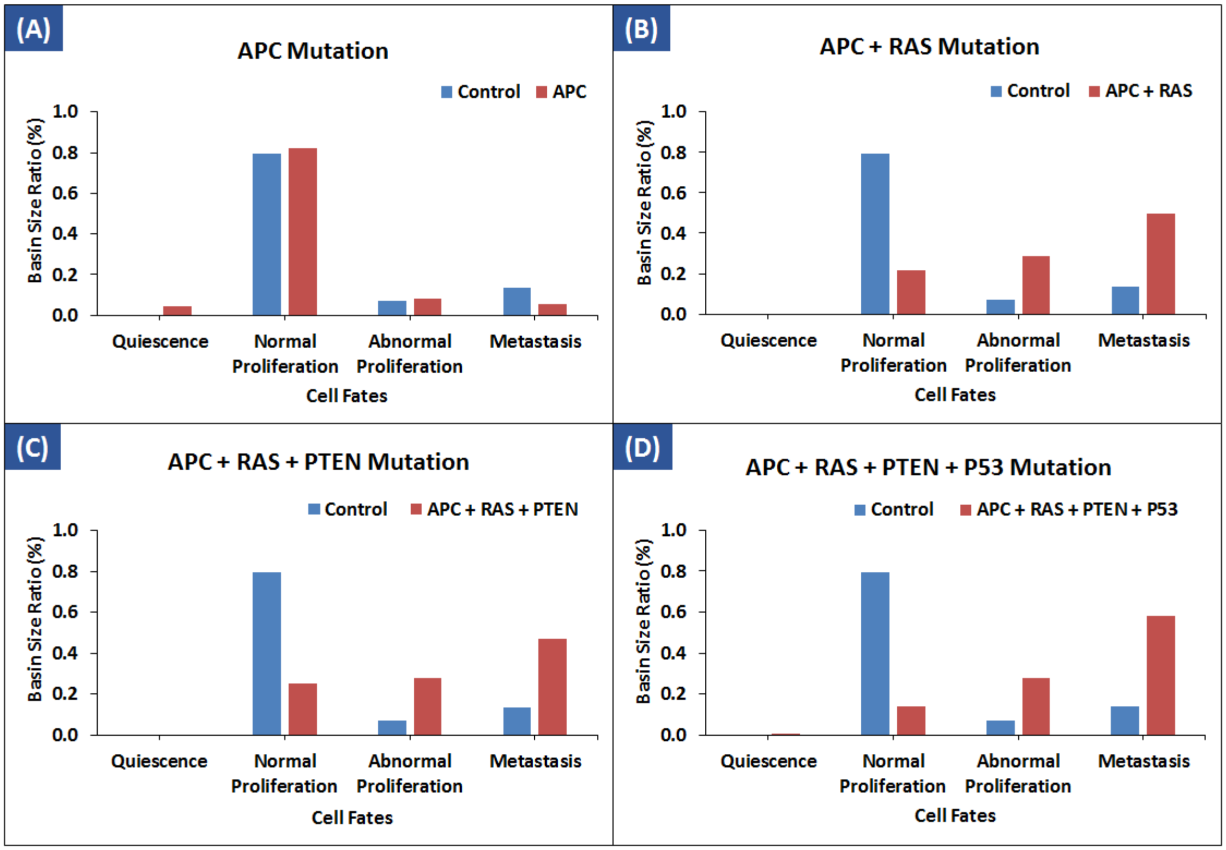

**Figure S29 – Case Study 2: Cell Fates Comparisons between Control and Mutations Induced Therapies.** (A) Comparison of cell fates obtained from control case and APC mutation. The basin size ratio for quiescence in control is (0) and (0.043) in APC mutation, for normal proliferation it is (0.793) in control and (0.822) in APC mutation, for abnormal proliferation it is (0.070) in control and (0.082) in APC mutation and for metastasis the basin size ratio is (0.137) in control and (0.053) in APC mutation, (B) Comparison between the cell fates of control therapy model and APC+RAS mutations model. The attractors along with their basin size ratios are: quiescence (0) in both control and APC + RAS mutation, normal proliferation (0.793) in control and (0.215) in APC + RAS mutation, abnormal proliferation (0.070) in control and (0.289) in APC + RAS mutation and metastasis (0.137) in control and (0.496) in APC + RAS mutation, (C) Comparison between the cell fates of control therapy model and APC+RAS+PTEN mutations model. The attractors along with their basin size ratios are: quiescence (0) in both control and APC + RAS + PTEN mutation, normal proliferation (0.793) in control and (0.254) in APC + RAS + PTEN mutation, Abnormal Proliferation (0.070) in control and (0.278) in APC + RAS + PTEN mutation and metastasis (0.137) in control and (0.470) in APC + RAS + PTEN mutation. (D) Comparison between the cell fates of control therapy model and APC+RAS+PTEN+P53 mutations model. The attractors along with their basin size ratios are: quiescence (0) in control and (0.004) APC + RAS + PTEN + P53 mutation, normal proliferation (0.793) in control and (0.139) in APC + RAS + PTEN + P53 mutation, abnormal proliferation (0.070) in control and (0.278) in APC + RAS + PTEN + P53 mutation and metastasis (0.137) in control and (0.583) in APC + RAS + PTEN + P53 mutation.

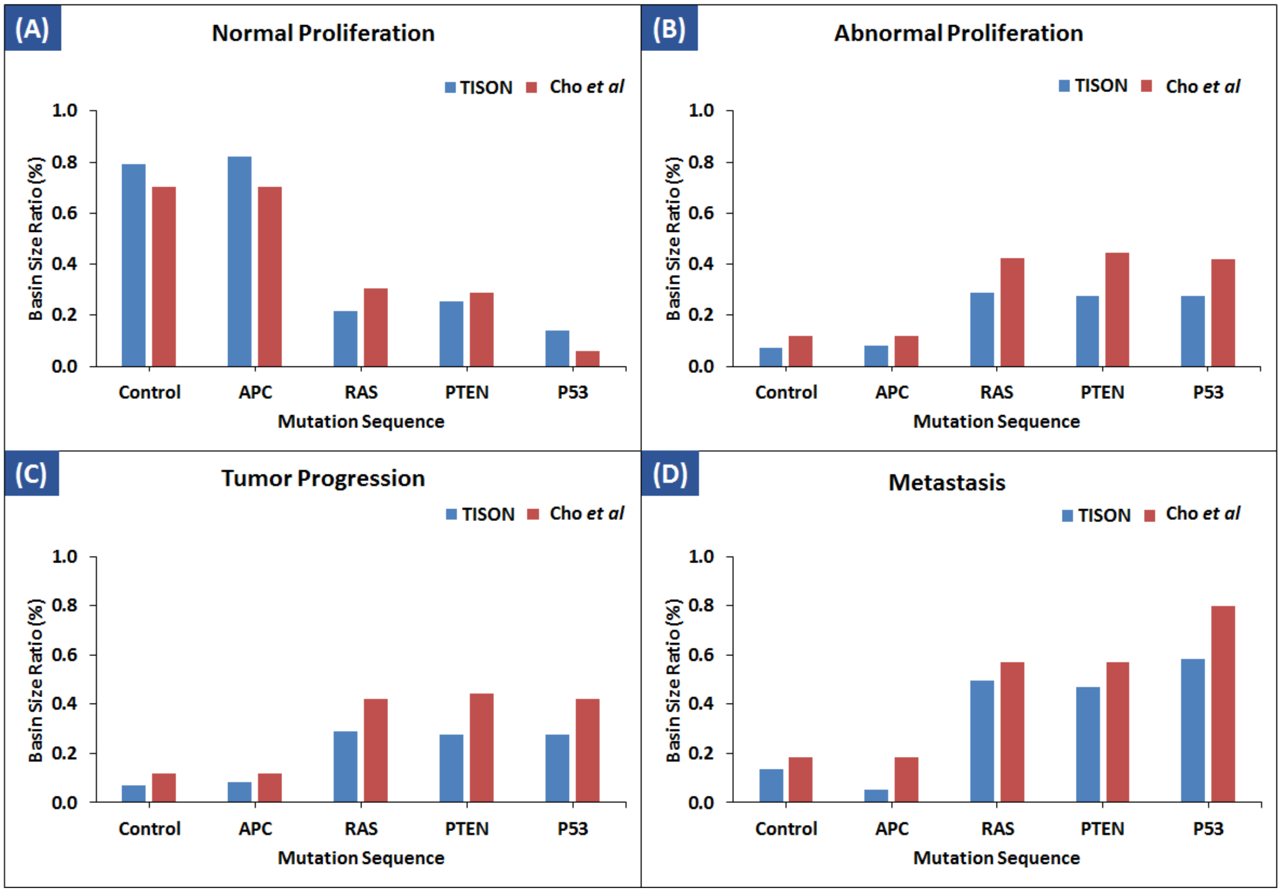

**Figure S30 – Case Study 2: Cell Fates Comparisons between Deterministic Analysis Results of TISON and Cho *et al*.'s Network Models.** (A) Basin size ratios of normal proliferation attractor in control and four mutation models, (B) Basin size ratios of abnormal proliferation attractor in control and four mutation models, (C) Basin size ratios of tumor progression attractor in control and four mutation models and (D) Basin size ratios of metastasis attractor in control and four mutation models.

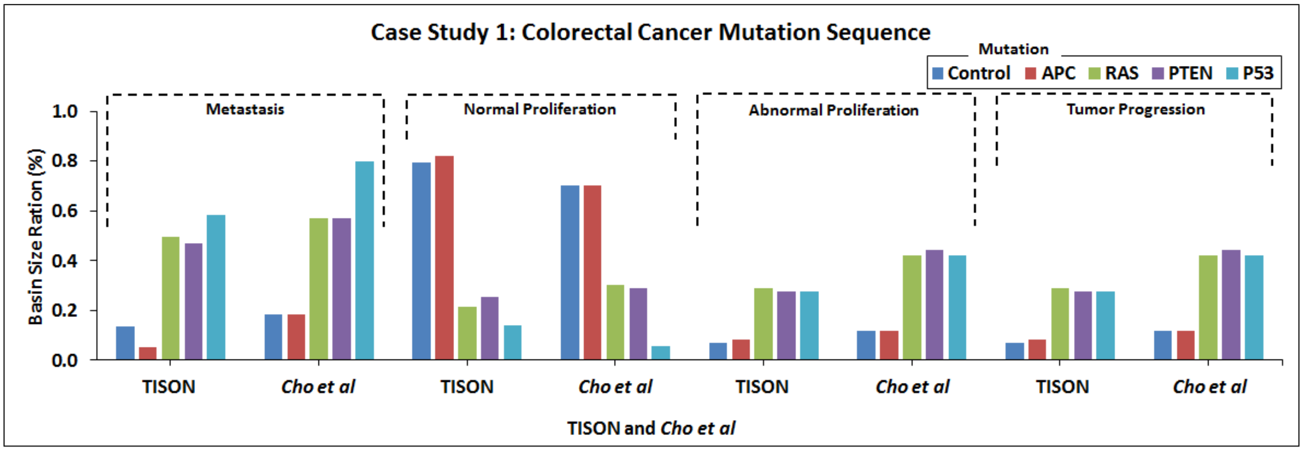

**Figure S31 – Case Study 2: A Summarized Comparison of Cell Fates between the Deterministic Network Analysis Results of TISON and Cho *et al*.'s Network models.** The figure shows the sequential comparison of resultant cell fates (metastasis, normal proliferation, abnormal proliferation, and tumor progression) derived from the network analysis of control and four mutation models.

#### Section 2.2.3. User Manual & Video Tutorial for Therapeutics Editor

**Table S2** - **Reference Table for TE**. Description of supporting information along with the web links have been tabulated below for the TISON home page, user manual, issues table, datasets, and video tutorials on YouTube as a playlist.

| Item | Description | Link |
|---|---|---|
| **Project Home** | URL link for TISON project home | https://tison.lums.edu.pk/ |
| **User's Manual** | Editor's user manual can be found here. | https://tison.lums.edu.pk/Manuals/TherapeuticsManual.pdf |
| **Issues** | Issues and related discussions can be added at the issues page. | https://github.com/BIRL/TISON/issues |
| **Datasets** | Sample Data files are available at the following link. | https://github.com/BIRL/TISON |
| **Playlist of Video Tutorials** | Video tutorials for step-by-step execution of case studies as well as employment of salient features | https://www.youtube.com/watch?v=OyvxS4YgP1I&list=PLaNVq-kFOn0ZUV0kGJCUJBbRw268xpbmo |

### Section 2.3. Environments Editor – An Overview

#### Section 2.3.1. Software Features & Functionalities

A key challenge in cancer modeling is to couple extracellular environments with user-designed intracellular models. TISON's Environments Editor (EE) aims to overcome this challenge by allowing its users to intuitively design cellular microenvironments followed by their seamless integration into cellular models. EE does this by taking a hybrid multi-scale modeling approach (8,9) by incorporating continuum Partial Differential Equation (PDE) (10) models of extracellular environments into discrete agent-based cellular models (ABMs) (9,11).

To access EE, users first have to create a project (using Project Explorer) followed by selecting EE from the Project Home Page or the editor bar at the bottom of the GUI.

- **Physical Time for an Environment**

Each environmental layer created using EE requires a pre-defined physical time to be used in synchronization with other models (see Section 2.3.4 for EE's User Manual). All layers defined within an environment are automatically assigned the same physical time. Users can input any physical time duration which is greater than zero. Time step increment can be set between 0 and maximum physical time defined earlier in the environment.

- **Running Environment**

Once an environment has been designed, users can proceed to simulate it by calculating and updating the concentration matrix within each layer of the model, at every time step. User-defined layer parameters assist in setting up the computation of heat diffusion PDE (shown below). Simulation results are iteratively calculated until the physical time specified for that particular environment.

$$\frac{\partial u}{\partial t} = Dc(\frac{\partial^2 u}{\partial x^2} + \frac{\partial^2 u}{\partial y^2} + \frac{\partial^2 u}{\partial z^2})$$

- **Visualizing and Downloading Environments Editor's Data**

Users can view and download meshes containing diffused environments from EE. Meshes can be visualized as 2D plots (x/y, x/z, and y/z) as well as in the form of a 3D plot. Note that 2D plots are projections of high-dimensional data onto a 2D plane, (i) xy: projection of environment data onto the xy-plane, (ii) xz: projection of environment data onto the xz-plane and (iii) yz: projection of environment data onto the yz-plane (see Section 2.3.4 for EE's User Manual). Furthermore, users can download the final layer matrix for visualization using 3rd party software like MATLAB (12), etc.

#### Section 2.3.2. Materials & Software Development Methodology

To solve the PDE (10), we first discretize them using an Alternate Direct Implicit (ADI) method to obtain a tridiagonal matrix (13,14). ADI is a finite difference method that can be applied for numerical estimation of the 3-dimensional heat equation with a complexity of $O(n^3)$ per time step(15). The equations are then solved using the Thomas algorithm (16). The algorithm is a special case of Gaussian elimination without pivoting for a tridiagonal matrix. It consists of two phases including forward reduction, and backward substitution. The forward reduction sequentially eliminates the lower diagonal of the original matrix, while the backward substitution sequentially solves for unknown variables using known variables and the upper and main diagonals in the resultant matrix.

**Table S3 – Sensitivity Analysis for EE**. The sensitivity analysis between COMSOL and TISON is displayed below:

| Tissue | Initial Condition | Boundary Condition | Mean Error (%) |
|---|---|---|---|
| Skin | 37 | 45 | 0.17 |
| Skin | 37 | 10 | 0.82 |
| Liver | 37 | 45 | 0.20 |
| Liver | 37 | 10 | 0.93 |

#### Section 2.3.3. User Manual & Video Tutorial for Environments Editor

**Table S4** - **Reference Table for EE**. Description of supporting information along with the web links have been tabulated below for the TISON home page, user manual, issues table, datasets, and video tutorials on YouTube as a playlist.

| Item | Description | Link |
|---|---|---|
| **Project Home** | URL link for TISON project home | https://tison.lums.edu.pk/ |
| **User's Manual** | Editor's user manual can be found here. | https://tison.lums.edu.pk/Manuals/EnvironmentsManual.pdf |
| **Issues** | Issues and related discussions can be added at the issues page. | https://github.com/BIRL/TISON/issues |
| **Datasets** | Sample Data files are available at the following link. | https://github.com/BIRL/TISON |
| **Playlist of Video Tutorials** | Video tutorials for step-by-step execution of case studies as well as employment of salient features | https://www.youtube.com/watch?v=O-bmM3DlNXE&list=PLaNVq-kFOn0YJXrXNXE541QHzcTI8l4lR&ab_channel=BiomedicalInformaticsResearchLab |

### Section 2.4. Cell Circuits Editor – An Overview

#### Section 2.4.1. Software Features & Functionalities

TISON's Cell Circuits Editor (CCE) assists in the development of cell decision systems in the form of finite state machines (FSM) (19,20). Users can access CCE from Project Home Page or the editor bar at the bottom of the GUI. Once a project is opened, the user can select CCE to initiate the construction of cell circuits. A cell circuit initiates with a "start" step and finishes at an "end" step. Users can incorporate biomolecular cues from networks and environments in the form of variables into each circuit. Additionally, users can select and initialize custom variables for equation-based processing and decision making within circuits. Several ready-to-use cell fates have also been provided in the editor including mitosis, quiescence, cell death, migration, and differentiation (see Section 2.4.3 for CCE's User Manual). Upon assembly, a cell circuit becomes available to the user that, upon execution, computes the emergent cell fates through a systematic temporal simulation.

#### Section 2.4.2. Functional Validation of the Cell Circuits Editor

To validate the functioning of CCE, we have reconstructed a literature-based case study in three stages, using  Gerlee and Anderson's "*minimal"* model (9). These case studies can be accessed either through the "Upload Case Study" button in CCE or through the template project tab on the TISON's home page. The stage-wise development of the case study is elaborated in the sub-sections below.

##### Stage 1 – Reconstruction of Gerlee and Anderson*'s "Minimal"* Model

Firstly, Gerlee and Anderson*'s* minimal model was reconstructed in TISON's CCE. The circuit was employed to study cell population growth under normoxia and hypoxia. Local variables were defined for oxygen, and age for onward programming of cell fate outcomes (Figure S32) (see Supplementary Data – CCE for further details).

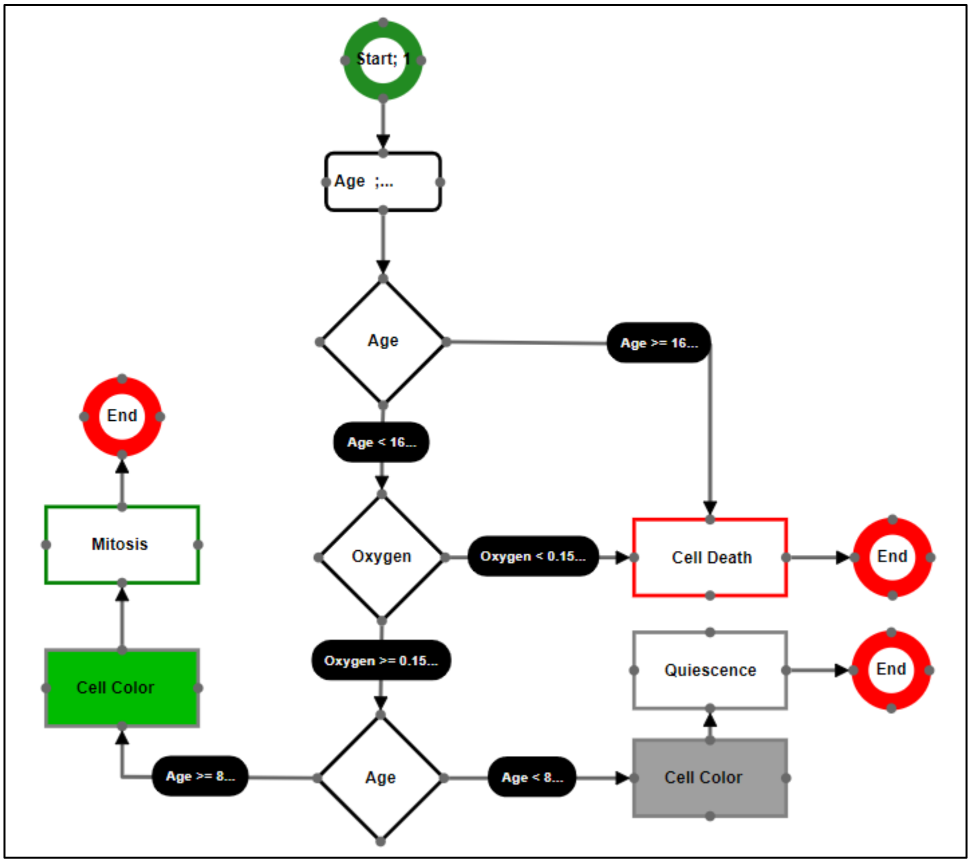

**Figure S32 - Stage 1: Reconstruction of Gerlee and Anderson*'s "Minimal" Model as a* Cell Circuit in TISON**

##### Stage 2 – Incorporation of Environment into the "*Minimal*" Model

In the second stage, we incorporated an extracellular environmental layer containing oxygen with an initial concentration of 1 and diffusion constant of 1.8 X $10^{-9}$ $cm^2$ $s^{-1}$, under Dirichlet boundary condition to study cell population growth under normoxia and hypoxia. The cell circuit programmed cells to either divide, go into quiescence, or perform cell death based on environmental conditions (Figure S33) (see Supplementary Data – CCE for further details).

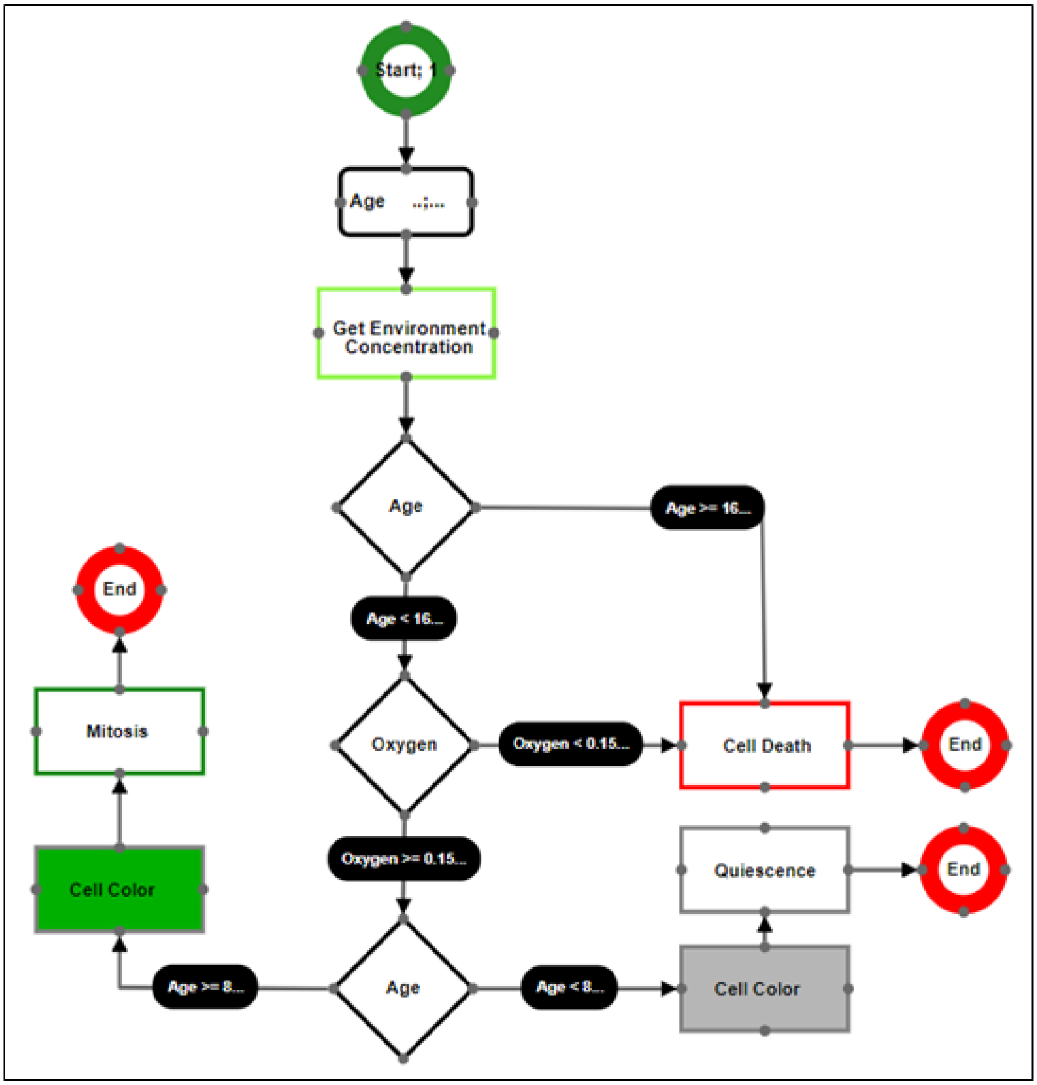

**Figure S33 - Stage 2: Integration of Environment into Gerlee and Anderson's "*Minimal"* Model using TISON's CCE.**

**Stage 3 – Integration of Biomolecular Networks having DNA Damage ON and OFF along with Extracellular Environments into the "*Minimal*" Model**

Finally, the "*minimal*" model was enhanced to include both network and environment variables from NE and EE, respectively. The network imported for this case study was Choi *et al.*'s p53 network having two variants including DNA damage ON and OFF (4). Oxygen was diffused with a diffusion constant of 1.8 X $10^{-9}$ $cm^2$ $s^{-1}$ in line with Gerlee and Anderson's "*minimal*" model (Figure S34-35) (see Supplementary Data – CCE for further details).

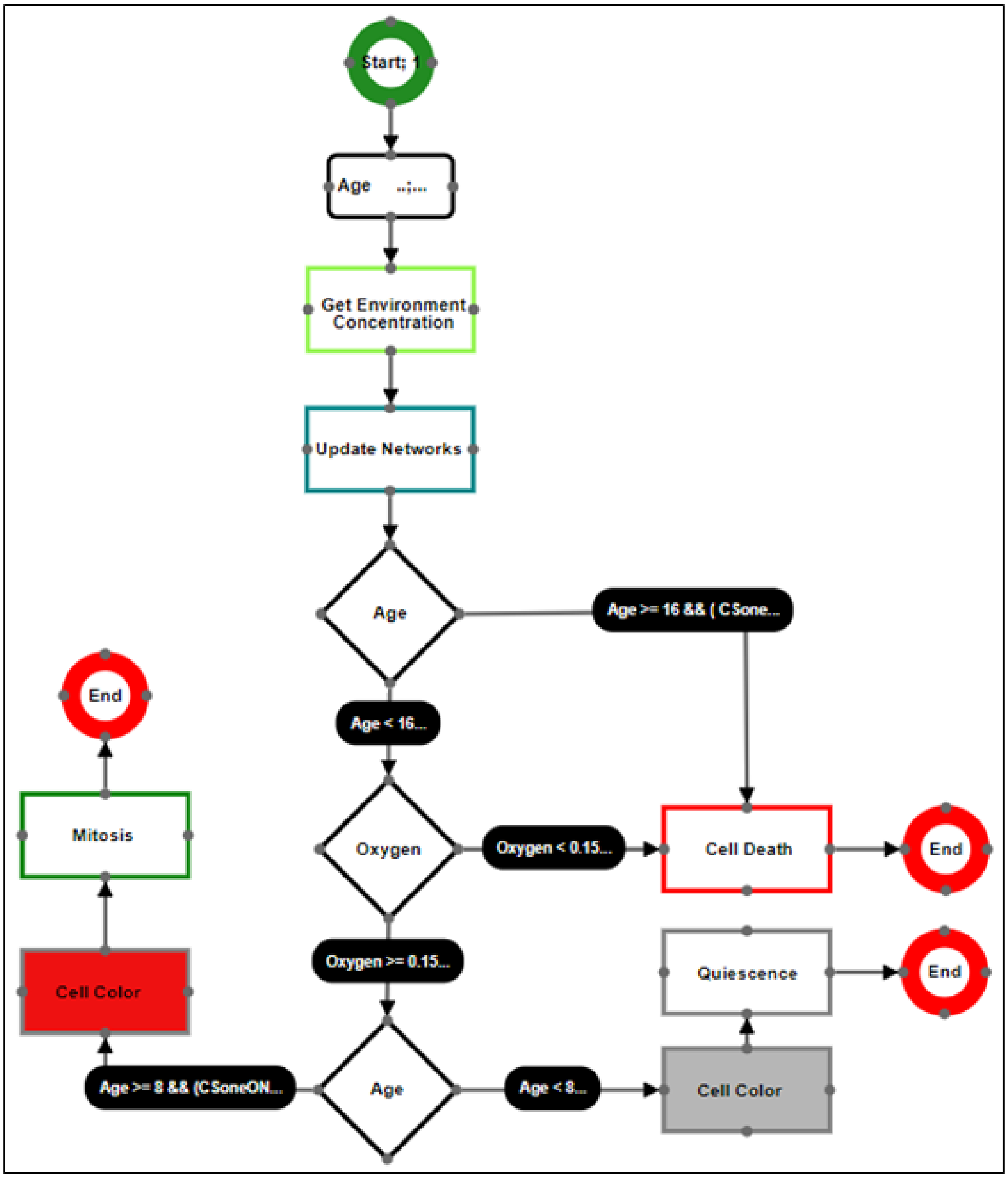

**Figure S34 - Stage 3(a): Cell Circuit of Gerlee and Anderson's "*Minimal*" Model after Incorporation of Network Regulation (DNA Damage ON) and Environment in TISON's CCE.**

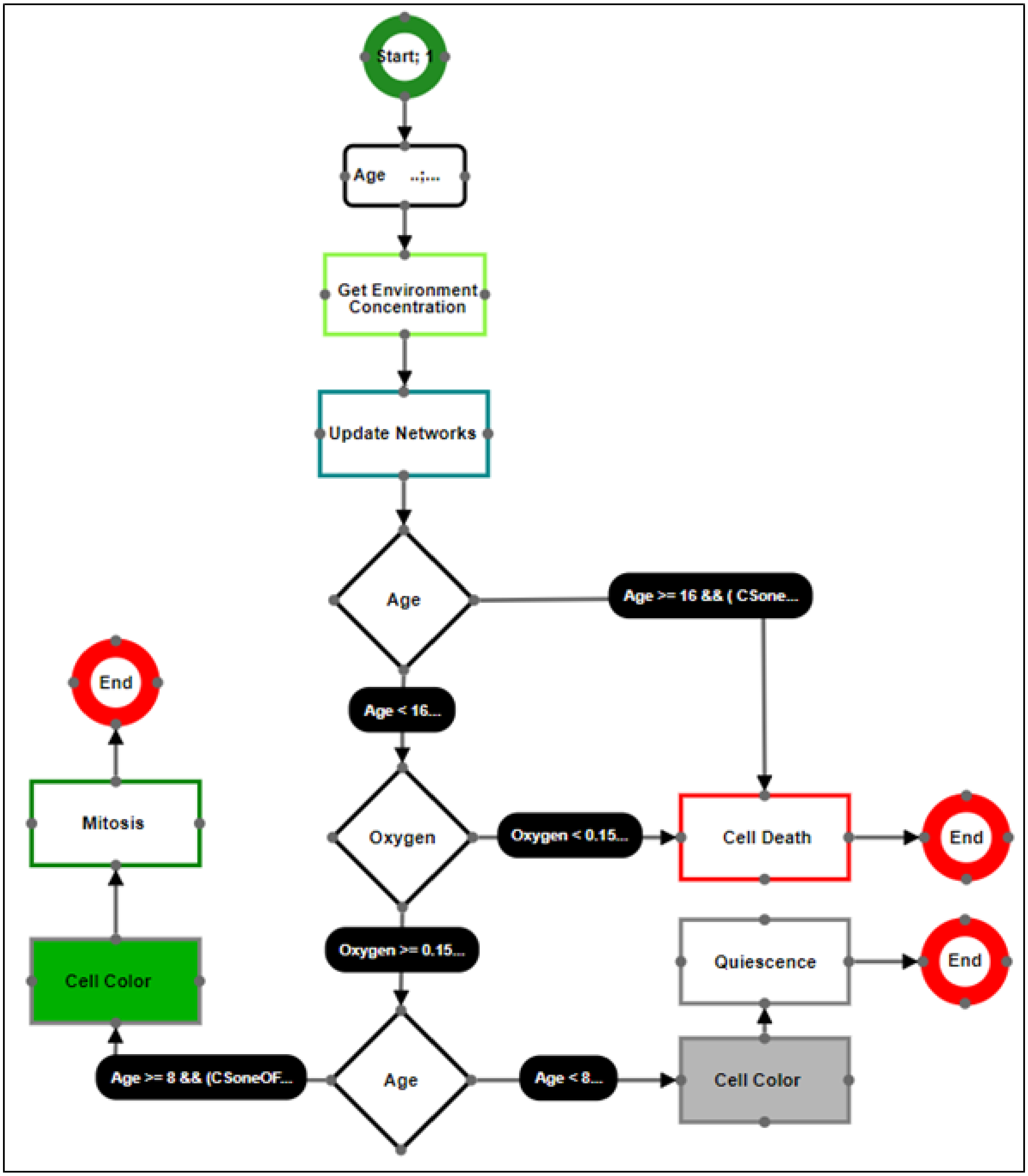

**Figure S35 - Stage 3(b): Cell Circuit of Gerlee and Anderson's "*Minimal*" Model after Incorporation of Network Regulation (DNA Damage OFF) and Environment in TISON's CCE.**

#### Section 2.4.3. User Manual & Video Tutorial for Cell Circuits Editor

**Table S5** - **Reference Table for CCE**. Description of supporting information along with the web links have been tabulated below for TISON home page, user manual, issues table, datasets, and video tutorials on YouTube as a playlist.

| Item | Description | Link |
|---|---|---|
| **Project Home** | URL link for TISON project home | https://tison.lums.edu.pk/ |
| **User's Manual** | Editor's user manual can be found here. | https://tison.lums.edu.pk/Manuals/CellCircuitsManual.pdf |
| **Issues** | Issues and related discussions can be added at the issues page. | https://github.com/BIRL/TISON/issues |
| **Datasets** | Sample Data files are available at the following link. | https://github.com/BIRL/TISON |
| **Playlist of Video Tutorials** | Video tutorials for step-by-step execution of case studies as well as employment of salient features | https://www.youtube.com/watch?v=SuSJF5pg_nM&list=PLaNVq-kFOn0a-jdi6oaitmNsuz3Y3mNue&ab_channel=BiomedicalInformaticsResearchLab |

### Section 2.5. Cell Lines Editor – An Overview

#### Section 2.5.1. Software Features & Functionalities

TISON's Cell Lines Editor (CLE) can be accessed by creating a TISON project (using Project Explorer) followed by selecting CLE either from the Project Home Page or from the editor bar at the bottom of the GUI.

Each cell line is assigned a specific cell circuit designed using CCE (details in Section 2.4). This enables the user to associate the cell line phenotype with a corresponding network and environment through variables from the NE and EE, respectively (detailed in Section 2.1 and 2.3, also see Section 2.5.3 for CLE's User Manual).

#### Section 2.5.2. Functional Validation of the Cell Lines Editor

We have undertaken the functional validation of CLE by constructing two cellular phenotypes representing cancerous and benign breast cell lines. These case studies can be accessed through the "Upload Case Study" button in CLE or through the template project tab on TISON's home page.

**Case Study 1 – Creating an *in silico* MCF-7 breast cancer epithelium cell line with DNA damage ON**

For this case study, we undertook the *in silico* replication of the MCF-7 breast epithelium cell lines using TISON. The resulting cell line was coupled with a cell circuit linked to a biomolecular network under DNA damage (Figure S36) (see Supplementary Data – CLE for further details.

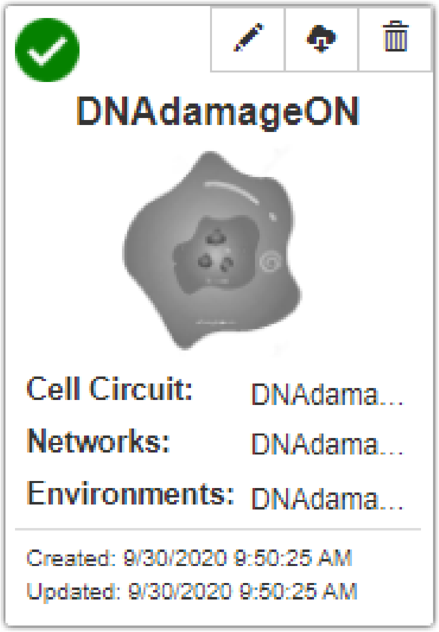

**Figure S36 – Case Study 1: MCF-7 with DNA Damage ON**

**Case Study 2 – Creating an *in silico* MCF-7 breast cancer epithelium cell line with DNA damage OFF**

In this case study, we constructed an in silico MCF-7 cell line representing a benign breast epithelial cell. The cell line was associated with a cell circuit containing an embedded biomolecular network without any DNA damage (OFF) (Figure S37) (see Supplementary Data – CLE for further details).

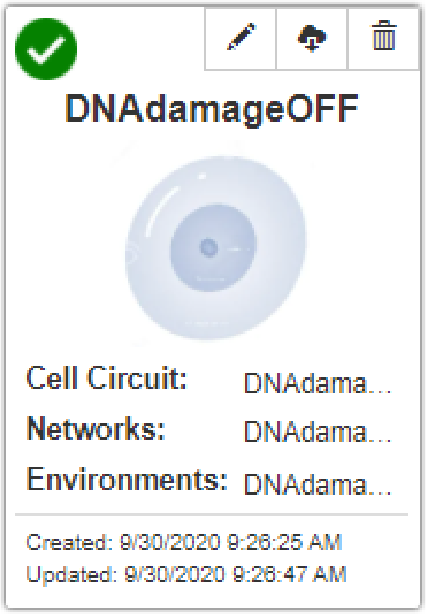

**Figure S37 – Case Study 2: MCF-7A with DNA Damage OFF**

#### Section 2.5.3. User Manual & Video Tutorial for Cell Lines Editor

**Table S6** - **Reference Table for CLE**. Description of supporting information along with the web links have been tabulated below for the TISON home page, user manual, issues table, datasets, and video tutorials on YouTube as a playlist.

| Item | Description | Link |
|---|---|---|
| **Project Home** | URL link for TISON project home | https://tison.lums.edu.pk/ |
| **User's Manual** | Editor's user manual can be found here. | https://tison.lums.edu.pk/Manuals/CellLinesManual.pdf |
| **Issues** | Issues and related discussions can be added at the issues page. | https://github.com/BIRL/TISON/issues |
| **Datasets** | Sample Data files are available at the following link. | https://github.com/BIRL/TISON |
| **Playlist of Video Tutorials** | Video tutorials for step-by-step execution of case studies as well as employment of salient features | https://www.youtube.com/watch?v=acoWwrI_f50&list=PLaNVq-kFOn0Z5ugt9uqA1ALfSfpPoRQn-&ab_channel=BiomedicalInformaticsResearchLab |

### Section 2.6. Organoids Editor – An Overview

TISON's Organoids Editor (OE) can be used to assemble three-dimensional *in silico* tissue organoids using *in silico* cell lines (detailed earlier in Section 2.5). OE can assist in designing multi-tissue organoid systems wherein multiple organoid elements can come together to form up to a whole organ. Users can access OE either from Project Home Page or from the editor bar at the bottom of the GUI.

#### Section 2.6.1. Software Features & Functionalities

- **Creating Organoids Elements**

To design three-dimensional *in silico* tissue organoids in the form of cell geometries, users first have to create "organoids elements" by using cell lines developed in CLE (detailed in Section 2.5). Each cell line comes ready with an embedded cell circuit (detailed in Section 2.4), associated biomolecular networks (detailed in Section 2.1), and an extracellular environment (detailed in Section 2.3). Organoid elements further come together to form multi-tissue systems. To create tissue elements, users can employ one of three methods including (i) a range, (ii) through an equation, or (iii) directly uploading a file with spatial coordinates of the organoid. Specifically, "drawing with a range" allows users to position cells within a coordinates bounds. The "draw with equation" feature allows users to draw at the vertices (x, y, and z coordinates on the surface) of any geometrical shape as calculated by a mathematical equation. The "draw with coordinates" enables users to input a set of x, y, and z coordinates to design a custom three-dimensional organoid. TISON also allows setting up of colors for each organoid element for ease of visualization (see Section 2.4.3 for OE's User Manual).

- **Resupply Points**

Users can build a link between organoid and environment editor (detailed in Section 2.3) using the "resupply points" option. This feature enables the user to specify coordinate traces for resupply locations wherein each trace will be assigned a resupply type. The user can draw the resupply traces using ranges, equations, or coordinates (detailed above). Currently resupply types available in TISON include: (i) blood vessel, (ii) lymphatic vessel, and (iii) nerve (see Section 2.4.3 for OE's User Manual).

- **Presets**

OE provides a set of predefined organoid templates for ready-use within the editor. These presets are provided as mathematical equations for the editor (see Section 2.4.3 for OE's User Manual).

#### Section 2.6.2. Functional Validation of the Organoids Editor

We have undertaken functional validation of OE by constructing two organoids representing cancerous and benign breast organoids. These case studies can be accessed through the "Upload Case Study" button in OE or through the template project tab on TISON's home page.

**Case Study 1 – MCF-7 Organoid with DNA damage ON**

In the first case study, we have constructed a breast organoid using the MCF-7 cell line developed previously in CLE. The cell lines embed the DNA damage ON network (detail in Section 2.5) (see Supplementary Data – OE for further details).

**Case Study 2 – MCF-7 Organoid with DNA damage OFF**

In the second case study, we have designed breast organoid with MCF-7 *in silico* cell line. The cell lines have the embedded DNA damage OFF network (detail in Section 2.5) (see Supplementary Data – OE for further details).

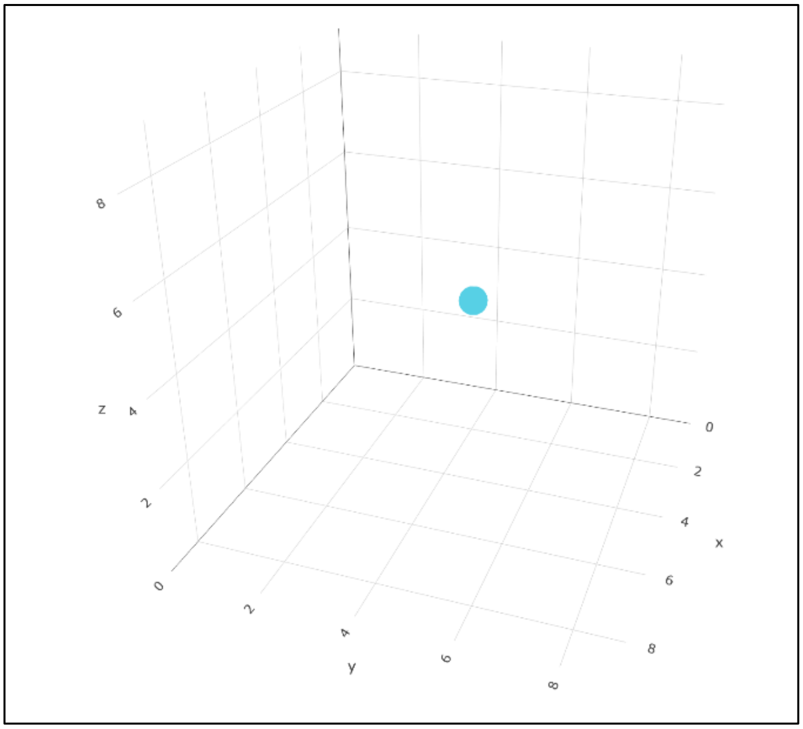

**Figure S38 – MCF-7 Organoid Placement in OE**

#### Section 2.6.3. User Manual & Video Tutorial for Organoids Editor

**Table S7** - **Reference Table for OE**. Description of supporting information along with the web links have been tabulated below for the TISON home page, user manual, issues table, datasets, and video tutorials on YouTube as a playlist.

| Item | Description | Link |
|---|---|---|
| **Project Home** | URL link for TISON project home | https://tison.lums.edu.pk/ |
| **User's Manual** | Editor's user manual can be found here. | https://tison.lums.edu.pk/Manuals/TissuesManual.pdf |
| **Issues** | Issues and related discussions can be added at the issues page. | https://github.com/BIRL/TISON/issues |
| **Datasets** | Sample Data files are available at the following link. | https://github.com/BIRL/TISON |
| **Playlist of Video Tutorials** | Video tutorials for step-by-step execution of case studies as well as employment of salient features | https://www.youtube.com/watch?v=2TY7iLri48E&list=PLaNVq-kFOn0b6numWR0VT9nQ4gi8y0AGM&ab_channel=BiomedicalInformaticsResearchLab |

### Section 2.7. Simulations Editor – An Overview

#### Section 2.7.1. Software Features & Functionalities

TISON's Simulations Editor (SE) assists its users to simulate the spatiotemporal evolution of organoids, constructed using OE (detailed earlier in Section 2.6). SE integrates data from TISON's editors, which includes the biomolecular regulatory networks and environments associated with cell decision circuits, and organoids. SE thereby enables TISON users to decipher the multifactorial interplay at each scale within these multiscale models (see Section 2.7.3 for SE's User Manual). Users can access SE either from Project Home Page or from the editor bar at the bottom of the GUI. Simulation generated in SE can be replayed, aborted, exported as well as users can choose to replay simulation using the simulation draggable slider at the bottom of the page. Once a simulation is being played, users can observe the exact time step and clock time for the duration of the simulation to see the temporal and spatial evolution of the modeled organoid.

#### Section 2.7.2. Functional Validation of the Simulations Editor

To exemplify the features provided by SE, we have constructed four literature-based case studies on MCF-7 breast epithelium cell lines including: (i) DNA damage ON, and (ii) DNA damage OFF. For that, two variants of Choi *et al.'s* p53 network (4), with DNA damage 'ON' and DNA damage 'OFF' were created using NE. An oxygen microenvironment was designed using EE and coupled with the networks through a "*minimal"* cell circuit model reported by Gerlee and Anderson (9), using CCE. The resultant cell circuits (having networks with DNA damage ON and OFF, in the presence and absence of therapies) are used to create two *in silico* cell lines in CLE. Next, breast epithelium organoids are designed in OE using these *in silico* cell lines and simulated in SE. These case studies can be accessed through the "Upload Case Study" button in SE or through the template project tab on TISON's home page. The case studies are discussed below in detail:

**Case Study 1 – Results from MCF-7 with DNA damage ON**

The first case study comprises the DNA damage ON - variant of Choi *et al.'s* p53 network, along with oxygen microenvironment integrated into a cell circuit. This circuit was used to create a cell line, followed by the development of an organoid model (see Supplementary Data – SE for further details).

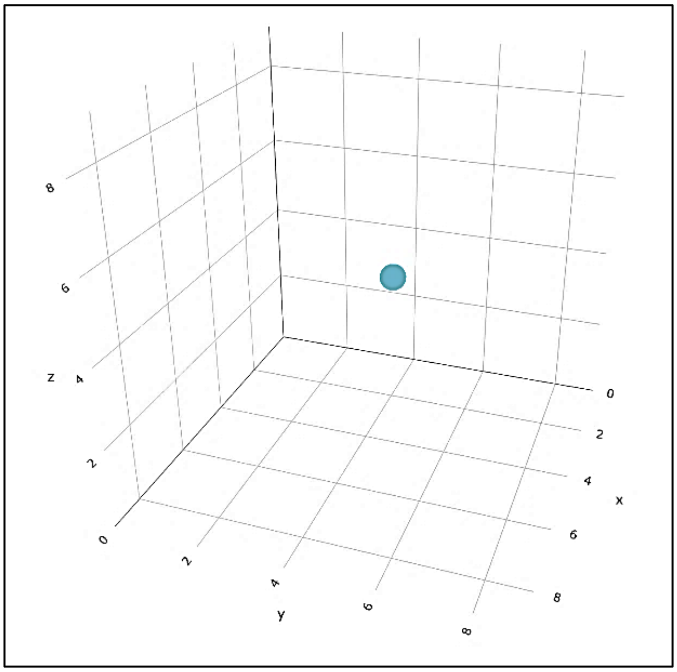

**Figure S39 – MCF-7 Organoid with DNA Damage ON, Simulated for 30 Time Steps.**

**Case Study 2 – Results from MCF-7 Organoid with DNA damage OFF**

The second case study employs the DNA damage OFF - a variant of Choi *et al.'s* p53 network, along with oxygen microenvironment integrated into a cell circuit. This circuit was used to create a cell line, followed by the development of an organoid model. (see Supplementary Data – SE for further details).

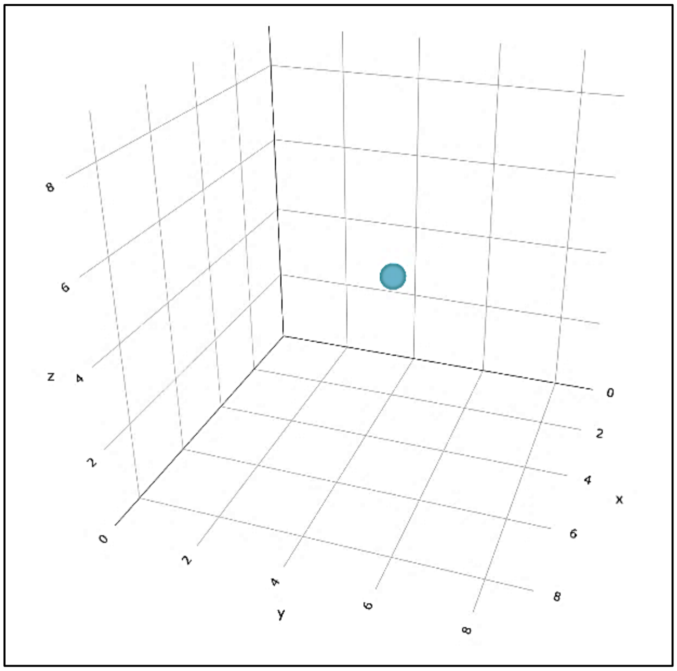

**Figure S40 – MCF-7 Organoid with DNA Damage OFF, Simulated for 30 Time Steps.**

#### Section 2.7.3. User Manual & Video Tutorial for Simulations Editor

**Table S8** - **Reference Table for SE**. Description of supporting information along with the web links have been tabulated below for: TISON home page, user manual, issues table, datasets, and video tutorials on YouTube as a playlist.

| Item | Description | Link |
|---|---|---|
| **Project Home** | URL link for TISON project home | https://tison.lums.edu.pk/ |
| **User's Manual** | TISON User manual can be found here. | https://tison.lums.edu.pk/Manuals/SimulationsManual.pdf |
| **Issues** | Issues and related discussions can be made at the issues page for TISON. | http://203.135.63.101:81/Projects/Index |
| **Datasets** | Sample Data files are available at the following link, for TISON. | https://github.com/BIRL/TISON |

| | | |
|---|---|---|
| **Playlist of Video Tutorials** | Video tutorials for step-by-step execution of case studies as well as employment of salient features of TISON. | https://www.youtube.com/watch?v=P6tWw7leuPE&list=PLaNVq-kFOn0aDz08h0waoOuJJwwpjaOqZ&ab_channel=BiomedicalInformaticsResearchLab |

1

### Section 2.8. Analytics Editor – An Overview

#### Section 2.8.1. Software Features & Functionalities

TISON's Analytics Editor (AE) allows users to visualize data obtained from the temporal and spatial simulation of cancer models simulated in SE. Users can perform multiple queries on the data generated from SE and observe the underlying biomolecular regulations involved in processes such as tumorigenesis and cancer growth. Users can access AE from the Project Home Page or the editor bar at the bottom of the GUI. Depending on the simulation results, users can perform the following queries in AE: (i) Cell Population (Total), (ii) Cell Population w.r.t Cell Line, (iii) Organoid Center (iv) Oragnoid Mobility (v) Oragnoid Parameters, (vi) Environment Consumption (vii) Cell Circuit Variables, and (viii) Cell Count w.r.t Color (see Section 2.8.3 for AE's User Manual).

#### Section 2.8.2. User Manual & Video Tutorial for Analytics Editor

**Table S9** - **Reference Table for AE**. Description of supporting information along with the web links have been tabulated below for TISON home page, user manual, issues table, datasets, and video tutorials on YouTube as a playlist.

| Item | Description | Link |
|---|---|---|
| **Project Home** | URL link for TISON project home | https://tison.lums.edu.pk/ |
| **User's Manual** | Editor's user manual can be found here. | https://tison.lums.edu.pk/Manuals/AnalyticsManual.pdf |
| **Issues** | Issues and related discussions can be added at the issues page. | http://203.135.63.101:81/Projects/Index |
| **Datasets** | Sample Data files are available at the following link. | https://github.com/BIRL/TISON |
| **Playlist of Video Tutorials** | Video tutorials for step-by-step execution of case studies as well as employment of salient features | https://www.youtube.com/watch?v=vGFV0vNtvXA&list=PLaNVq-kFOn0aEAGHZE3PzmcTEfzpdbAoY&i |

| | | |
|---|---|---|
| | | ndex=2&t=0s&ab_channel=BiomedicalInformaticsResearchLab |

### Section 3. Step-by-step Development of Multi-scale Models using TISON's Eight Editors

#### Section 3.1.1. Step-by-step Development Pipeline in TISON

**Step 1.** We will construct biomolecular regulatory networks for each case study, using existing literature and online patient databases in TISON's ***Networks Editor*** **(NE)**. These include Catalogue of Somatic Mutations in Cancer (COSMIC) (21), Genotype-Tissue Expression (GTEx) (22), The Cancer Genome Atlas (TCGA) (23,24), Metabolic gEne Rapid Visualizer (MERAV) (25), Genomic Data Commons (GDC) (26), The Cancer Proteome Atlas (TCPA) (27), Human Protein Atlas (HPA) (28) etc. Next, we will carry out robustness analysis of six models with minor perturbations and evaluate them against random signals. Once, networks are confirmed to be biologically plausible showing high robustness against random perturbations, we will validate the resulting networks using physiological/normal, stress and cancerous conditions. The cell fates obtained will be matched with literature and the output node propensities will be tallied with experimental values taken from database. Cell fate validation of the networks will be performed using TISON. NE will be used to perform topological, deterministic and probabilistic analyses for eliciting emergent properties from biomolecule cancer networks.

**Step 2.** The biomolecular targets identified during step 1 will be subjected to therapeutic screening using TISON's ***Therapeutics Editor*** **(TE).** TE will then help to develop therapeutic screens for identification of novel drug targets, drug repurposing and personalized therapeutics. Presence of gene expression data integration from various spatiotemporal scales, will render therapeutic screening strategies reliable and coherent with experimental data on large-scale networks.

**Step 3.** To design the extra-cellular matrix (ECM) (29) of a cell, we will employ TISON's ***Environments Editor*** **(EE)** which allows its user to create environmental models to mimic normal or tumor microenvironments (30) and later integrate them into cellular models. The environmental milieu is reflective of the conditions surrounding a cell and is designed to elicit exact biological responses. EE assists in the creation of diffusive microenvironments towards modelling the dynamical engagement of environmental cues with cellular tissue organoids. EE will implement continuum models of extra-cellular environments using Partial Differential Equations (PDE) (10).

**Step 4.** To couple the biomolecular networks with extra-cellular entities like drugs, nutrients and cellular secretions (ligands or ROS), cellular circuits will be developed using finite state automata theory (19) in TISON's ***Cell Circuit Editor*** **(CCE)**. CCE can integrate information from biomolecular networks and extra-cellular environments (constructed using NE and EE, respectively) and assemble them in the form of "*Cell Circuits*". Each node in a circuit defines the execution of specific processes such as environmental uptake of nutrients and drugs.

**Step 5.** Next, TISON's ***Cell Lines Editor*** **(CLE)** will be used to create *in silico* cell lines such as MCF7; breast cancer cell line, HCT-116; colon cancer cell line etc. Each cell line will be assigned a specific cell circuit design using CCE. This will enable the user to associate the cell line phenotype with a corresponding environment and network from the EE and NE, respectively. CLE will also provide stand-alone repositories with data on *in silico* cell lines for creating enhanced and more accurate *in silico* cell models.

**Step 6.** To assemble the *in silico* cell lines into three-dimensional organoids, termed "*organoids*" TISON's ***Organoids Editor*** **(OE)** will be used. The organoids can be designed to resemble nerve tissue, lymphatic vessels, epithelium lining etc. OE will also provide an intuitive biopsy tissue image processing feature that will seamlessly convert clinical images into *in silico* organoids.

**Step 7.** To simulate organoid growth and evolution over time, TISON's ***Simulations Editor*** **(SE)** will be employed. SE will simulate the tissue organoid models for calculating their temporal evolution under specific environmental conditions and allow users to visualize the results. The editor will also allow its users to observe organoid progression according to the variables and rates defined in the organoid and circuit editors. Users can then investigate the morphology of the organoid given the temporal and spatial nature of the environment using the clock time and time steps feature of SE. SE can be used to investigate organoid development as well as undertake comparisons between disease and homeostatic organoid evolution.

**Step 8.** Lastly, simulation data generated in SE will be queried in the ***Analytics Editor*** **(AE)** for onward quantitative data analysis. The resultant multi-scale models will be analyzed to view patient specific therapeutic responses in AE which will then allow for the statistical analysis and visualization of data generated by the SE. AE will enable its users to view temporal variations in cell population as well as cell line specific properties from simulation data. Depending on the simulation results, users will be able to plot (i) cell and phenotype populations (total) over time, (ii) statistics on cell-type specific properties and variables, (iii) concentration of extra-cellular environments, (iv) cellular phylogeny amongst others.

### Section 4. References

1. Ermentrout GB, Edelstein-Keshet L. Cellular automata approaches to biological modeling. J Theor Biol. 1993;160(1):97–133.

2. Kim Y, Choi S, Shin D, Cho KH. Quantitative evaluation and reversion analysis of the attractor landscapes of an intracellular regulatory network for colorectal cancer. BMC Syst Biol. 2017;11(1):1–22.

3. Cho S-H, Park S-M, Lee H-S, Lee H-Y, Cho K-H. Attractor landscape analysis of colorectal tumorigenesis and its reversion. BMC Syst Biol. 2016;10(1):96.

4. Choi M, Shi J, Jung SH, Chen X, Cho K-H. Attractor Landscape Analysis Reveals Feedback Loops in the p53 Network That Control the Cellular Response to DNA Damage. Sci Signal [Internet]. 2012;5(251):ra83--ra83. Available from: https://stke.sciencemag.org/content/5/251/ra83

5. Han B, Wang J. Quantifying robustness and dissipation cost of yeast cell cycle network: the funneled energy landscape perspectives. Biophys J. 2007;92(11):3755–63.

6. Li C, Wang J. Quantifying Cell Fate Decisions for Differentiation and Reprogramming of a Human Stem Cell Network : Landscape and Biological Paths. 2013;9(8).

7. Guo J, Lin F, Zhang X, Tanavde V, Zheng J. NetLand: Quantitative modeling and visualization of Waddington's epigenetic landscape using probabilistic potential. Bioinformatics. 2017;33(10):1583–5.

8. Osborne JM, Walter A, Kershaw SK, Mirams GR, Fletcher AG, Pathmanathan P, et al. A hybrid approach to multi-scale modelling of cancer. Philos Trans R Soc A Math Phys Eng Sci. 2010;368(1930):5013–28.

9. Gerlee P, Anderson ARA. An evolutionary hybrid cellular automaton model of solid tumour growth. J Theor Biol. 2007 Jun;246(4):583–603.

10. Villadsen J, Michelsen ML. Solution of differential equation models by polynomial approximation. Vol. 7. Prentice-Hall Englewood Cliffs, NJ; 1978.

11. Bonabeau E. Agent-based modeling: Methods and techniques for simulating human systems. Proc Natl Acad Sci. 2002;99(suppl 3):7280–7.

12. MATLAB [Internet]. Available from: https://www.mathworks.com/products/matlab.html

13. Chang MJ, Chow LC, Chang WS. Improved alternating-direction implicit method for solving transient three-dimensional heat diffusion problems. Numer Heat Transf Part B Fundam. 1991;19(1):69–84.

14. 3-D Heat Equation Numerical Solution [Internet]. Available from: https://www.mathworks.com/matlabcentral/fileexchange/59336-3-d-heat-equation-numerical-solution

15. Douglas J, Rachford HH. On the Numerical Solution of Heat Conduction Problems in Two and Three Space Variables. Trans Am Math Soc. 1956;82(2):421.

16. Zhang R. Applied Contaminant Transport Modeling: Theory and Practice. Vol. 25, Journal of Environment Quality. 1996. p. 927.

17. Comsol AB. COMSOL multiphysics reference manual. Version. 2007;

18. Rivera MJ, Molina JAL, Trujillo M, Romero-García V, Berjano EJ. Analytical validation of COMSOL Multiphysics for theoretical models of Radiofrequency ablation including the Hyperbolic Bioheat transfer equation. In: 2010 Annual International Conference of the IEEE Engineering in Medicine and Biology. IEEE; 2010. p. 3214–7.

19. Mohri M. On some applications of finite-state automata theory to natural language processing. Nat Lang Eng. 1996;2(1):61–80.

20. Oishi K, Klavins E. Framework for Engineering Finite State Machines in Gene Regulatory Networks. ACS Synth Biol [Internet]. 2014 Sep 19;3(9):652–65. Available from: https://doi.org/10.1021/sb4001799

21. Bamford S, Dawson E, Forbes S, Clements J, Pettett R, Dogan A, et al. The COSMIC (Catalogue of Somatic Mutations in Cancer) database and website. Br J Cancer. 2004;91:355–8.

22. Lonsdale J, Thomas J, Salvatore M, Phillips R, Lo E, Shad S, et al. The Genotype-Tissue Expression (GTEx) project. Vol. 45, Nature Genetics. Nature Publishing Group; 2013. p. 580–5.

23. Lee HJ, Palm J, Grimes SM, Ji HP. The Cancer Genome Atlas Clinical Explorer: A web and mobile interface for identifying clinical-genomic driver associations. Genome Med [Internet]. 2015;7(1):1–14. Available from: http://dx.doi.org/10.1186/s13073-015-0226-3

24. Wang Z, Jensen MA, Zenklusen JC. A practical guide to The Cancer Genome Atlas (TCGA). In: Methods in Molecular Biology. Humana Press Inc.; 2016. p. 111–41.

25. Shaul YD, Yuan B, Thiru P, Nutter-Upham A, McCallum S, Lanzkron C, et al. MERAV: A tool for comparing gene expression across human tissues and cell types. Nucleic Acids Res. 2016;44(D1):D560–6.

26. Jensen MA, Ferretti V, Grossman RL, Staudt LM. The NCI Genomic Data Commons as an engine for precision medicine. Blood. 2017;130(4):453–9.

27. Li J, Lu Y, Akbani R, Ju Z, Roebuck PL, Liu W, et al. TCPA: A resource for cancer functional proteomics data. Vol. 10, Nature Methods. Nature Publishing Group; 2013. p. 1046–7.

28. Pontén F, Jirström K, Uhlen M. The Human Protein Atlas—a tool for pathology. J Pathol. 2008 Dec;216(4):387–93.

29. Warrick JW, Murphy WL, Beebe DJ. Screening the cellular microenvironment: a role for microfluidics. IEEE Rev Biomed Eng. 2008;1(1):75–93.

30. Whiteside TL. The tumor microenvironment and its role in promoting tumor growth. Oncogene. 2008 Oct;27(45):5904–12.
